# Subdural CMOS optical probe (SCOPe) for bidirectional neural interfacing

**DOI:** 10.1101/2023.02.07.527500

**Authors:** Eric H. Pollmann, Heyu Yin, Ilke Uguz, Agrita Dubey, Katie Elizabeth Wingel, John S Choi, Sajjad Moazeni, Yatin Gilhotra, Victoria A. Pavlovsky, Adam Banees, Vivek Boominathan, Jacob Robinson, Ashok Veeraraghavan, Vincent A. Pieribone, Bijan Pesaran, Kenneth L. Shepard

## Abstract

Optical neurotechnologies use light to interface with neurons and can monitor and manipulate neural activity with high spatial-temporal precision over large cortical extents. While there has been significant progress in miniaturizing microscope for head-mounted configurations, these existing devices are still very bulky and could never be fully implanted. Any viable translation of these technologies to human use will require a much more noninvasive, fully implantable form factor. Here, we leverage advances in microelectronics and heterogeneous optoelectronic packaging to develop a transformative, ultrathin, miniaturized device for bidirectional optical stimulation and recording: the subdural CMOS Optical Probe (SCOPe). By being thin enough to lie entirely within the subdural space of the primate brain, SCOPe defines a path for the eventual human translation of a new generation of brain-machine interfaces based on light.

## Introduction

Electrode-based interfaces dominate clinical applications of brain-machine interfaces that record from and stimulate the nervous system(*1*). However, basic neuroscience research in animal models has been transformed by optical techniques(*2*) which can precisely target defined populations of neurons for recordings using genetically encoded calcium(*3*) or voltage indicators(*4*) and for stimulation using light-sensitive opsins(*5*). Electrode recordings are governed by the screening of electric fields in cerebral spinal fluid, which causes a rapid fall off in the electrophysiological signal with screening lengths on the order of *λ_AP_* ≈ 20 μm (*6, 7*). In contrast, optical techniques are governed by the scattering and absorption of light allowing larger distances between sensors and neurons of more than 100 μm. This means that optical imaging approaches can interrogate neurons at greater distances than electrodes, enabling simultaneous recordings from a larger and more complete volumes of neurons(*8*). In this way, optical neurotechnologies deliver advantages over more conventional electrical interfacing approaches.

To-date, however, even miniaturized optical neurotechnologies have generally relied on microscopy techniques requiring bulky instrumentation. A proliferation of lens-based miniature microscopes(*9*) have been deployed in many applications, including high-speed volumetric imaging(*10*), wireless imaging(*11*), mesoscale field-of-view imaging(*12*), and two-photon imaging(*13*). Recently, lens-less computational miniature microscopes have improved upon the volumetric field-of-view by performing single-shot 3D imaging, using either a micro-lens array(*14, 15*), amplitude mask(*16*), or phase-mask(*17*). While more volumetrically efficient than table-top microscopes, current lens-less approaches use commercial off-the-shelf (COTS) instrumentation and need an opening in the dura and skull that matches or exceeds the field-of-view (FoV). These constraints ultimately preclude fully implanted translational applications.

In this work, we develop the SCOPe (Subdural CMOS Optical Probe) system, a miniaturized, mechanically flexible, less than 200-μm-thick, lens-less miniature microscope and optical stimulator. SCOPe dramatically improves on the volumetric efficiency of implantable optoelectronics for neural interfaces and introduces a new class of brain-machine interface devices that are thin enough to be contained within the subdural space of the primate brain. This radical improvement in volumetric efficiency is achieved through the design of a custom complementary metal-oxide-semiconductor (CMOS) application-specific integrated circuit (ASIC) which is capable of both fluorescence imaging and optogenetic stimulation. The integrated circuit chip, thinned to less than 15 μm so as to be mechanically flexible(*18–20*) and conform to the cortical surface, is packaged on a flexible polyimide printed circuit board along with the heterogeneous integration of the light sources, filters, and lens-less masks required for a high-performance optical neural interface. We demonstrate basic functionality of the SCOPe device for imaging and optical stimulation *in vivo* in the mouse model over a subset of the array. We then validate full-FoV application by imaging neurons expressing GCaMP8m in the motor cortex of the non-human primate (NHP) and demonstrating the ability to decode reach movement speed trajectories moment-by-moment.

## Results

### Displacement factor

To compare the invasiveness of various miniaturized optical implantable systems, we define the displacement factor (DF) as being the ratio of the volume of the implanted device to the imaged tissue volume. Using this definition, we contrast the SCOPe form-factor with the conventional miniature microscope(*10*) (**Fig. 1**A), quantitatively calculate the DF of our device (**Fig. 1**B), and compare it with lens-based and lensless miniature microscopes(*9–15, 17*) (**Table S1**). Most notably, the SCOPe is capable of imaging a FoV that is greater than its own volumetric displacement, occupying approximately 7.5 mm^3^ while being able to image a 3D volumetric FoV of 15 mm^3^. This corresponds to a DF of 0.5, while lens-less fluorescence imagers(*17*) have DF of 66 and lens-based fluorescence imagers(*10*) have DF of 1428. While the miniaturization of SCOPe does tradeoff resolution for FoV, its resolution is comparable to other miniature mesoscopes(*12*) providing similar FoVs at significantly larger volumes (DF in excess of 350).

**Fig. 1.**
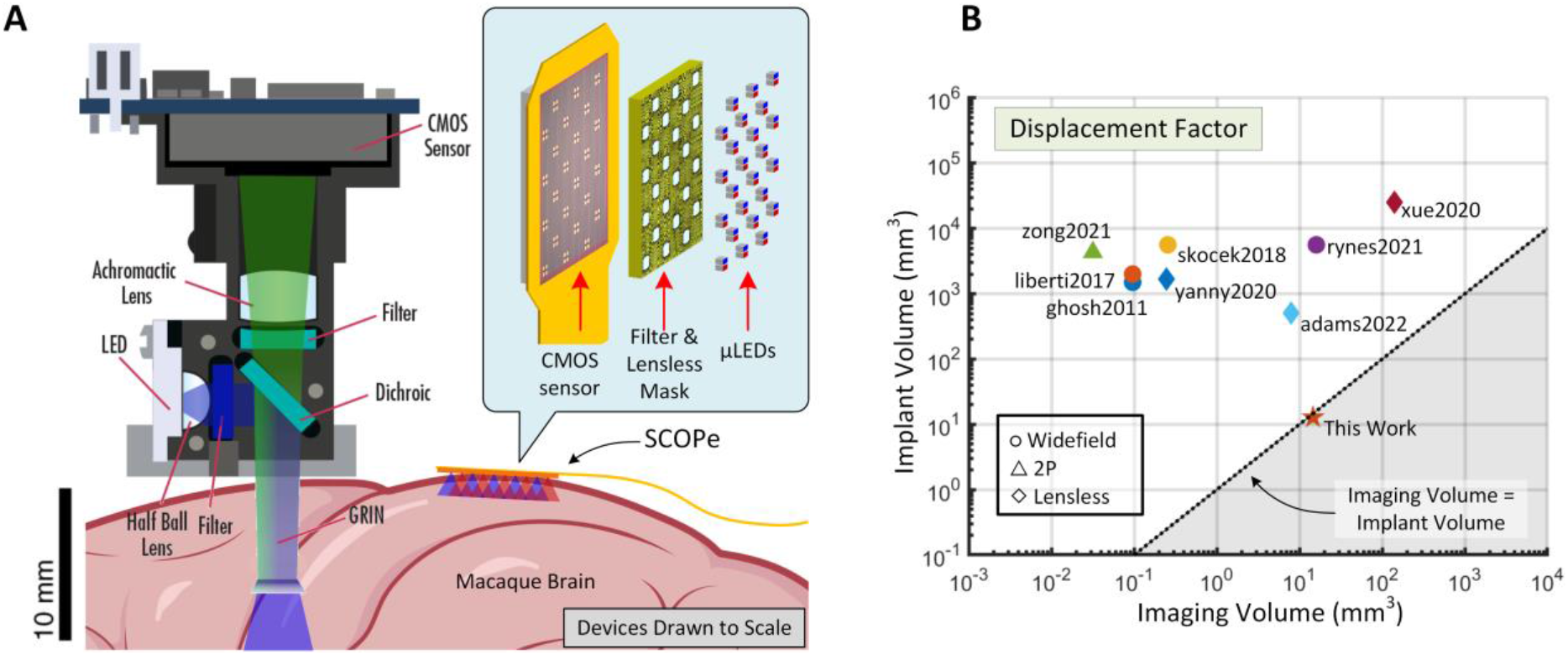
Comparison of displacement factor of miniature microscopes. (A) To-scale size comparison of the Miniscope(*10*) and the SCOPe developed here. (B) Graph showing comparison of displacement factors of SCOPe along with current state-of-the-art(*9–15, 17*) miniscopes, plotting implant volume on the y-axis and imaging volume on the x-axis. Devices in the bottom-right of the graph are more desirable.

### CMOS ASIC design

The use of single-photon avalanche photodiodes (SPAD) has increased in recent years due to their miniaturization and monolithic integration into CMOS imaging arrays(*21*). They have been explored in many biophotonics applications including endoscopic FLIM(*22*), super-resolution microscopy(*23*), NIROT(*24*), and PET(*25*). The SCOPe system incorporates a custom CMOS ASIC that consists of an array of 192×256 monolithic SPAD detectors (**Fig. 2**A) for low-light-intensity imaging and dual color (470 nm blue and 590 nm red) flip-chip bonded micro-light-emitting diodes (μLED) as light sources for fluorescence excitation and optogenetic stimulation, respectively. The 6.4 × 7.8 mm^2^ integrated circuit and major sub-blocks are fabricated in a 130-nm high-voltage CMOS process with support for a custom SPAD design (**Fig. 2**B). The ASIC is equipped with lead-free solder bumps for flip-chip bonding to a flexible printed-circuit board (**Fig. 2**C, **Fig. S1**, and **Section S1**) which routes the reference clock (100 MHz, for maximum frame rate operation of 200fps), the scan-chain configuration signals, the voltage supplies (VDD of 1.5V, DVDD of 1.8V, VLED of 2.5V, and VSPAD of 20.9V), and the raw data stream to a field-programmable gate array (FPGA) which interfaces with a host computer to store and process the data (**Fig. 2**D). The drivers for the fluorescence-exciting blue μLEDs are configured for continuous-wave (CW) illumination for high signal-to-noise-ratio (SNR) imaging whereas the drivers for the optogenetic red μLEDs are configured on a frame-by-frame basis allowing for pulsing durations in increments of 5 ms at 200 fps.

**Fig. 2.**
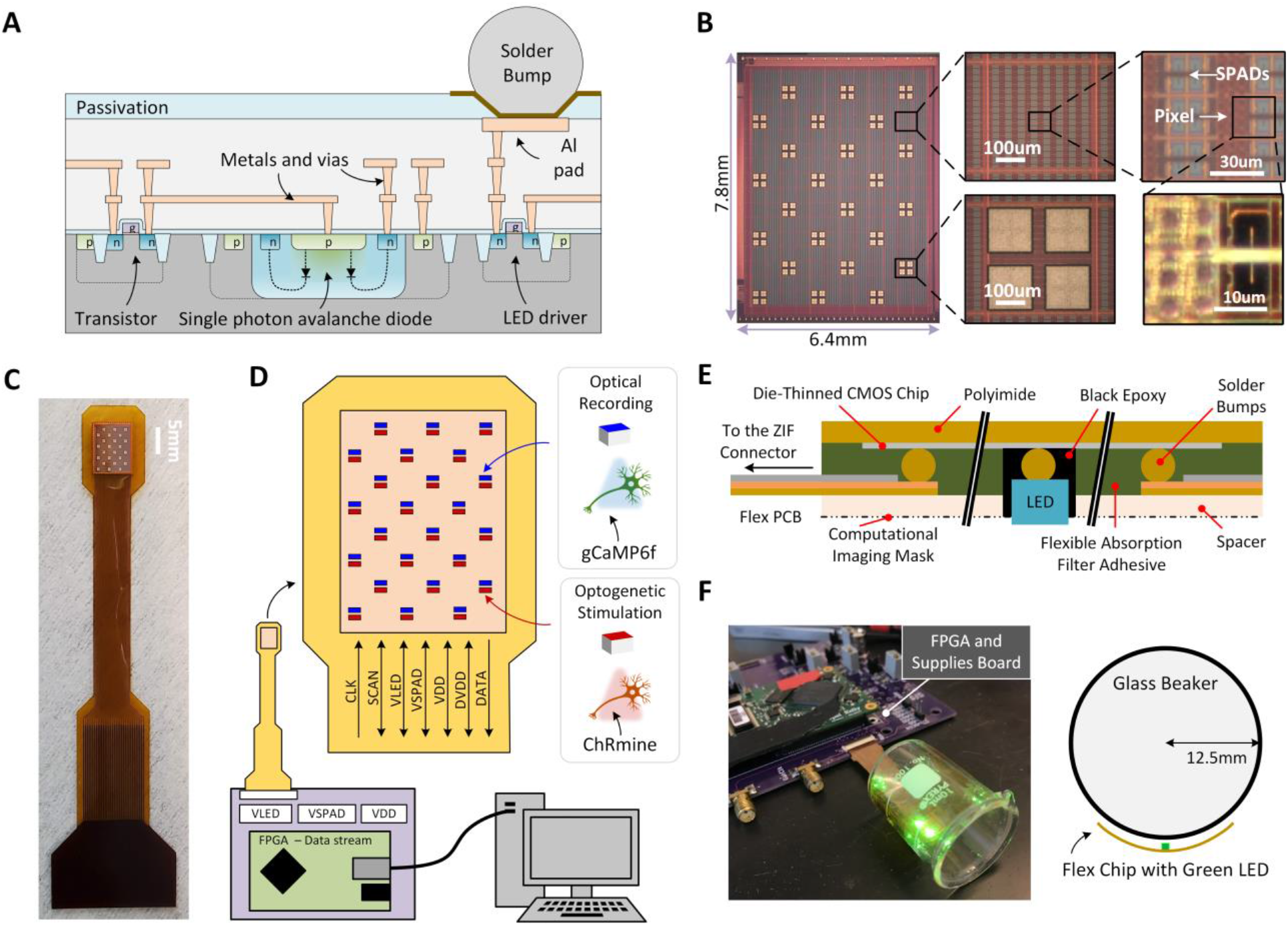
SCOPe device packaging and data acquisition system. (A) Cross section of integrated circuit showing monolithic integration of SPADs with planar bulk CMOS technology and solder bump pads for flip-chip packaging of μLED dice. (B) Die micrographs of the 6.4×7.8mm CMOS ASIC showing arrays of 16×16 SPADs, flip-chip bonded μLED pads, a cluster of pixels, and a single pixel. (C) Packaged ASIC in a flexible, conformable PCB. (D) Fluorescence and optogenetic stimulation configuration modes with individually multiplexed blue and red μLEDs, data acquisition system including scan-chain input-output configurability, power supply generation, and data stream-out. (E) Flexible packaging incorporating the filters, LEDs and computational mask in a less than 250-μm-thick form-factor. (F) SCOPe device curved around a labware beaker with 25-mm diameter and the controller and LEDs illuminated to demonstrate device functionality.

### Single photon avalanche diodes

A more detailed overview of the chip block diagram is given in **Fig. S2** with a more detailed discussion of the circuit design in **Section S1**. In photon counting mode, SPAD performance is governed by noise equivalent power (NEP) and dynamic range (DR)(*26*). NEP is the minimum signal intensity required for an SNR of 1 within a 1 Hz bandwidth and is derived from the photon-detection probability (PDP), fill factor, and dark count rate (see Methods). We can also definite a photon-detection efficiency (PDE) which is the product of the PDP and fill factor (see **Section S5**). A lower NEP denotes better performance, and our sensor achieves 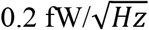 at 520 nm given a measured PDP of 9% at 550 nm and excess bias of 1.5 V, a fill-factor of 11.5%, and a dark-count-rate rate (DCR) of 17 Hz. The DR of the sensor is important in fluorescence microscopy preparations since a large amount of specular reflection occurs at the tissue surface and insufficient spectral filtering can result in image saturation. The DR of SCOPe is 60dB as determined by the per-pixel 10-bit counters. Further description of the characterization of these SPADs is given in **Fig. S3**. We compared our SPAD sensor with other SPAD arrays(*19, 27–31*) (see **Table S2**). At 200 fps, our sensor shows higher dynamic range and lower power consumption than other comparable SPAD imaging systems(*30, 31*).

### Sensor power consumption

The inherent low-noise performance of SPADs, a result of low DCR and zero read noise, enables low-power sensing which is important for implantable applications to limit tissue heating. Since the SPAD is an activity-based high-gain sensor (**Fig. S4**), power consumption scales with light intensity. This contrasts with conventional CMOS image sensors (CIS) which use photodiodes and rely on electronic amplification at a power overhead to get low-noise performance at low light levels. The SPAD image sensors on SCOPe at 200 fps consume only 18 mW from 1.5V VDD supply and 160 μW from the 20.9V VSPAD supply in the dark. As light intensity increases, the dynamic power consumption on VDD increases in the in-pixel counters and data transmission blocks. Then, as the light levels increase and the counters start to rollover, dynamic power consumption on VDD saturates at approximately 21 mW and only the SPAD power consumption continues to increase with increasing intensity. Sensor performance can be compared across circuit architectures and technologies by defining a circuit figure-of-merit (FoM) which is the ratio of noise-power product to the pixel rate (*32*). The noise is defined as the square root of the dark count rate, and the pixel rate is the sensor readout rate considering sensor array size and readout speed (**Section S5**). As shown in **Fig. S4**, at light levels of < 5 nW/mm^2^, comparable to the that produced by fluorescent emission in SCOPe applications, the SPAD image sensor shows comparable or better FoM to state-of-the-art CIS across various circuit architectures.

### System packaging and mechanical flexibility

Designing mechanically flexible imagers with high resolution have typically required the use of flexible organic materials paired with polycrystalline silicon thin-film transistors(*33*). Unfortunately, such devices lack the resolution, power, and noise performance of CMOS devices. In SCOPe, we achieve mechanical flexibility while still employing CMOS. The full packaging of the SCOPe system is shown in **Fig. 2**E. These packaging components include the μLED light sources, flexible optical filters, and computational imaging mask on a thin polyimide spacer, all mounted atop a flexible polyimide printed-circuit board (PCB). The Methods and **Fig. S5** detail the packaging flow that renders the final device. Prior to device implantation, the chip is conformal coated with 6 μm of parylene (**Fig. S6**).

The laminated components of the implant are all made of flexible materials except for the CMOS die and dielectric interference filter (**Table S3**) which are rendered flexible by thinning. Flexure becomes limited by the stresses introduced into the die with curvature. For a Young’s Modulus (*E*) of 85 GPa, Poisson’s ratio (*v*) of 0.21, and a substrate thickness of 15 μm, the minimum radius of curvature (*R_min_*) is 4mm, assuming a maximum stress of 200 MPa (70) for bulk silicon (see Methods). In practice, this radius of curvature is closer to 12.5 mm (**Fig. 2**F), because the CMOS die with interference filter is a more complex materials stack and defects created by backside thinning become nucleation site for fracture. This curvature, however, is sufficient to conform to brains as small as the mouse(*34*), although the device is too large for subdural implantation in mice.

### Light sources

For fluorescence excitation, coherent laser light sources have traditionally been utilized in conjunction with fiber-optic bundles(*35*), tapered optical fibers(*36*), or nanophotonic silicon probes(*37, 38*) to provide narrow spectral linewidths; however, this approach requires complex packaging and precludes a fully-integrated implanted device because of the need for external light sources. Other integrated laser sources, such as vertical-cavity surface-emitting lasers (VCSELs), could be considered, but coherence is not a requirement in this application and the more spectrally broad light emission of LEDs can be managed with excitation filters. We, therefore, focus on the use of these simpler non-coherent LED sources. Organic LEDs(*39–41*), which provide an extremely scalable and ultrathin form-factor, still typically have optical power densities of less than 0.2 mW mm^−2^ at 5V(*41*), which are insufficient for fluorescence excitation, and suffer spectral linewidths in excess of 77nm(*42*). Instead, we choose solid-state μLEDs based on the InGaN/GaN materials system, which offer high external quantum efficiencies (EQE) and optical power densities up to 70 mW mm^−2^ at 3V (**Fig. 3**A). Various options for optical sources are compared in **Fig. S7**. When packaged, these μLEDs have significant light leakage directly from the sides and back of the μLED, which bypasses the filters and is captured directly by the adjacent SPAD detectors. To reduce this effect to an immeasurable level, we underfill the μLED with a black optically absorbing epoxy after attachment (see Methods).

**Fig. 3.**
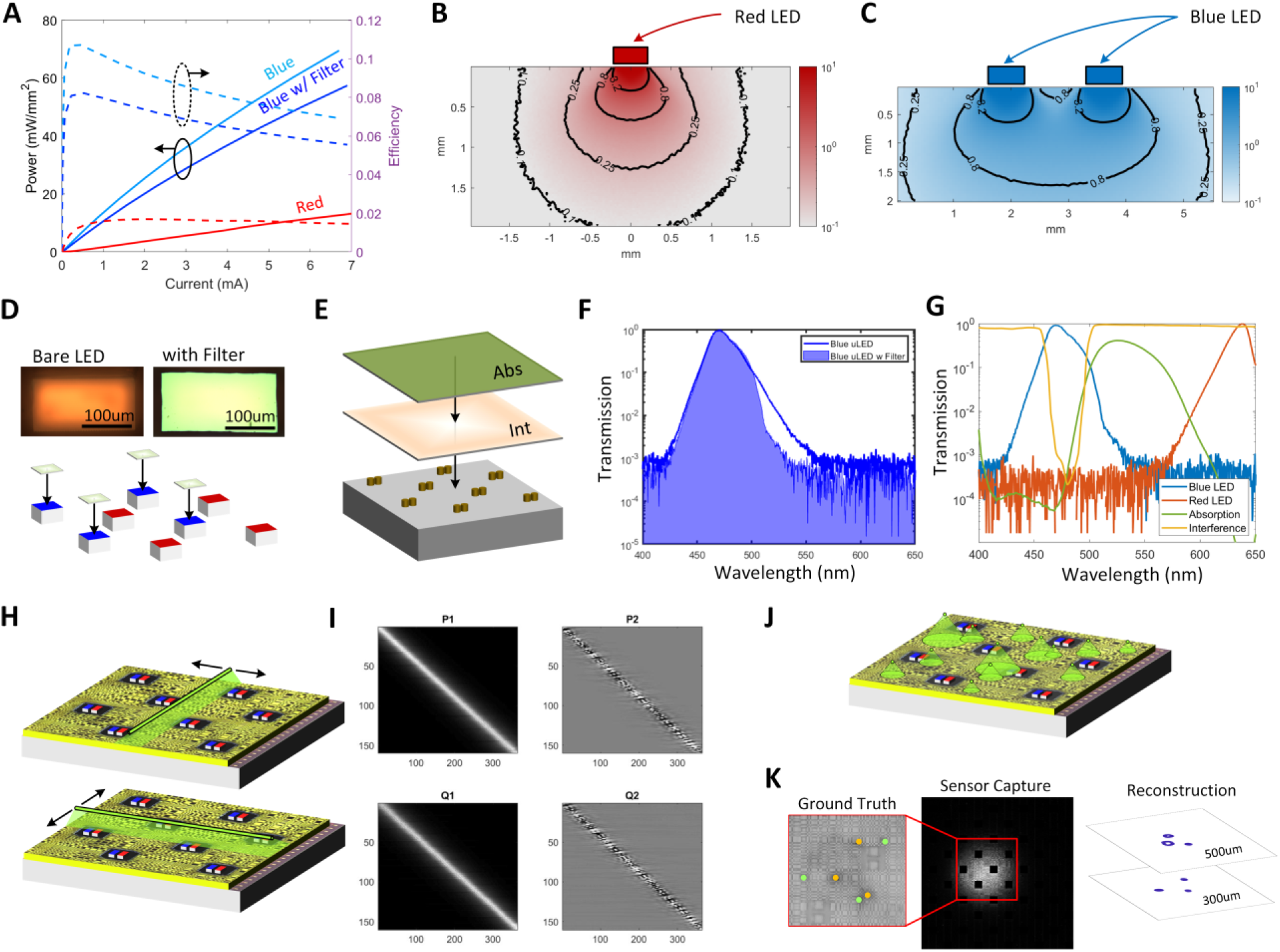
Co-integrated optical components of SCOPe. (A) Optical power density as a function of forward current for both blue and red μLED light sources. (B) Simulation of light propagation in scattering tissue for a single red (640nm) μLED used for optogenetic stimulation. (C) Simulation of depth of penetration of light for two blue (470nm) μLEDs for fluorescence imaging. (D) Micrograph images of the μLED with and without the interference filter directly deposited and a graphic showing selective deposition onto only blue μLEDs. (E) Graphic showing direct deposition of absorption (abs) and interference (int) filters onto CMOS chip. (F) Spectral plot of the 500-nm-short-pass excitation filter deposited on top of blue μLED. (G) Spectral plot of the 500-nm-long-pass hybrid absorption/interference emission filter. (H) Calibration process that sweeps a fluorescent line slit in both row and column directions for each z-plane. (I) Separable transfer functions where P1, P2 operate on the rows and Q1, Q2 operate on the columns. P1, Q1 show the open mask with no apertures and P2, Q2 show the coded mask. (J) Single-shot image capture and reconstruction process with fluorescent sources at varying z-depths above the sensor. (K) 3D imaging from a single image capture of six 45μm fluorescent beads, three of the beads (denoted in green) are located at 300 μm from the mask and three of the beads (denoted in yellow) are located at 500 μm. Contour lines show the normalized reconstructed images with a threshold of 0.6.

### Light propagation modeling

The measured optical power density of the red and blue μLEDs determines the programmable forward current levels needed to drive the μLEDs for optogenetic stimulation and fluorescence excitation, respectively (**Fig. 3**A). The SCOPe system incorporates two separate arrays of 24 blue (470nm) and 24 red (640 nm) μLEDs with 1.25-mm lateral pitch and 0.6 mm diagonal pitch for fluorescence excitation and optogenetic stimulation (**Fig. S8**), respectively. The CMOS drivers and red and blue InGaN μLEDs achieve currents in the excess of 7mA and optical output power densities of 10mW/mm^2^ and 60mW/mm^2^, respectively (**Fig. S9**). One of the key advantages of the SCOPe device is optical proximity, which allows significant reductions in the required optical power.

For optogenetic stimulation, a Monte Carlo model of light propagation(*43*) from a single red stimulation μLED is shown in **Fig. 3**B using optical parameters of cortical tissue at 640 nm, namely a scattering coefficient (*μ_s_*) of 9mm^−1^, an absorption coefficient (*μ_a_*) of 0.02mm^−1^, an anisotropy (*G*) of 0.89, and an index of refraction (*n*) of 1.36)(*44*). Near the tissue surface, higher fluence is observed due to scattering and internal reflection at the tissue-fluid interface(*45*). For ChRmine, a highly sensitive red-shifted opsin, the optogenetic neuronal depolarization threshold(*46*) light intensity is ~0.1 mW/mm^2^. This requires a minimum of 0.02 mW/mm^2^ (achieved with a forward current of 250 μA) at the surface of the red μLED to robustly activate the neuron at a depth of 200 μm directly over the μLED.

For fluorescence imaging using the blue μLEDs, the illumination pattern is determined by the Lambertian radiation profile, tissue scattering, and the pitch between the μLEDs. This pitch is chosen to maximize the photodetection area relative to light emission area on the CMOS image sensor (87.5%) (**Fig. S10**) while achieving uniform illumination (to within 75%) at a target depth of 200 μm in tissue. In this design we were limited by the μLED sizes available commercially. Shallower depths of uniform illumination could have been achieved by going to smaller μLED sizes while maintaining the same 87.5% photodetection-to-emission-area ratio. Given the 94-dB DR of the sensors, required optical filter is determined by SNR performance requirements. Noise is dominated by the photon-shot noise of the unrejected backscattered illumination light. The amount of fluorescence signal detected is a function of the light propagation of both the excitation and fluorescence light (**Fig. S11**) as well as the brightness of the fluorescent probe (**Section S2**). A Monte Carlo model of light propagation(*43*) estimates the light path losses at a cortical depth of 200 μm (**Fig. 3**C) given the optical properties of cortical tissue at 470 nm, namely, *μ_a_*=0.21mm^−1^ (**Section S2**), *μ_s_*=10mm^−1^, *g*=0.9, and *n*=1.38(*47*). From these simulations, the amount of green fluorescence light returning to the detectors is 0.074% of the illumination intensity, while the amount of backscattered light is 7.5% of this same intensity (**Fig. S11**). The minimum required optical density (OD) of the filter can then be determined from the DR of SCOPe and the SNR requirements (see **Section S5** and **Fig. S12**). In particular, for an SNR of 5 and restricting the dynamic range to only 40 dB, which is less than 10% of full-scale and corresponds to the excitation light levels employed in these experiments, an OD of 2 required.

### Filters

Achieving this degree of spectral filtering in the form factor of SCOPe is very challenging. In traditional fluorescence microscopy systems(*9–15, 17*), a cascade of collimating lenses and high-quality dichroic interference filters are used to isolate the emitted fluorescent light from the high-power excitation. However, these optical components are bulky and require focal lengths and working distances that are larger than the thickness of the SCOPe device. To achieve a truly implantable form factor without the use of collimating lenses or external light-sources, we designed an interference-based excitation filter, deposited directly to the flip-chip bonded μLEDs (**Fig. 3**D), and a hybrid absorption/interference emission filter, deposited and packaged directly on the surface of the imager (**Fig. 3**E). The excitation filter reduces the spectral bandwidth of the LED light source, providing an additional 10× rejection of the 525-nm out-of-band emission, which would cause undesired mixing with the fluorescence signal (**Fig. 3**F). The emission filter rejects the 470-nm blue excitation light in favor of 520-nm green emission. This hybrid interference and absorption filter(*48*) combines the higher rejection (peak OD > 5) provided by the long-pass interference filter at lower incidence angles (generally < 25°) with the wide-angle rejection of a bandpass absorption filter (**Fig. 3**G). The absorption filter provides rejection of both the blue excitation light (OD 4) as well as the red optogenetic stimulation light (OD 5) (**Fig. S13**). The μLED spectrum overlaps with the excitation spectrum of gCaMP6f, and the emission filter overlaps with the emission spectrum of gCaMP6f (**Fig. S14**). The spectral transmission of the gCaMP6f emission illumination, and the intrinsic sensor spectral responsivity is shown in **Fig. S15.** The total OD filtering achieved is better than 1.9, meeting the requirements outlined above.

### Computational imaging

SCOPe is designed to collect fluorescence at an effective focal plane at a controllable depth within the cortex. Traditional lens-based optics suffer from a fundamental tradeoff between size and performance; as the lens shrinks in size, the FoV does as well. A variety of lens-less computational imaging optical metasurfaces have been demonstrated to avoid these limitations of refraction optics (**Fig. S16**), such as angle-sensitive pixels(*18, 49*), random diffuser masks(*50*), micro-lens arrays(*14, 15*), coded-aperture amplitude masks(*16, 51*), and phase-masks(*17, 52*). We choose to use a separable coded-aperture amplitude mask(*16, 51*) because of small working distances required. This separability reduces the calibration and reconstruction complexity(*16*) as compared with other computational imaging methods (**Fig. S17** and **Section S3)**. The amplitude mask is incorporated 100 μm from the SPAD detectors and contacts the cortical surface. Unlike a conventional image sensor, our imager is missing 12.5% of the pixels due to the embedded μLEDs. The reconstruction accounts for these missing sensor pixels during reconstruction. Calibration is performed by sweeping a vertical and horizontal fluorescence line source across the imager FoV at each depth, *d*, of interest (**Fig. 3**H) to determine the separable transfer functions (**Fig. 3**I) required for reconstruction. The resolution of the imager is 60-μm at a depth of 200 μm as determined by a double line slit experiment and a USAF target, despite the loss of information due to the μLED positions (**Fig. S18**). To demonstrate the single-shot volumetric imaging of the SCOPe system (**Fig. 3**J), we took an image capture of six 45-μm fluorescent beads, in which three are located at d=300 μm and three are located at d=500 μm (**Fig. 3**K); this imaging is performed by computational refocusing of multiple z-depths.

### Power dissipation and heating

While heating is usually disregarded in the development of conventional miniscopes due to the thermal isolation of the heat source (μLEDs and CMOS image sensor) from the tissue, this is not appropriate for fully implanted devices. The design of a custom low-power integrated circuit allows us to limit the extraneous power consumption found in commercial general-purpose CMOS imaging systems, as well as to engineer the sensor front end for low-light biological scenes by using SPADs, as described earlier. In addition, the proximity of the optical device to the tissue interface allows for efficient optical interrogation with the biological scene without the efficiency losses of lenses and apertures. In our *in vivo* experiments, SCOPe achieves *in-vivo* imaging at 40 frames per second (fps) with a power consumption of less than 10 mW for the sensor and <110 μW per μLED, of which approximately 10 μW is the optical emission. (**Fig. S9** and **Table S4**). This contrasts with miniature mesoscopes(*12*) with similar FoVs and resolution which consume 226 mW in the image sensor and 31mW in the excitation LEDs. This translates to more than 13× less power consumption from our illumination μLEDs and 22× less power consumption from our image sensor. The heating of the device with an 200-μm-thick overlaid cortical slice was characterized using a FLIR infrared camera (see Methods) to be less than 1°C(*53*) up to electrical powers of 20mW and optical powers of 1.2mW, which is approximately two orders of magnitude larger than needed to image and stimulate neural activity (**Fig. S19**).

### *In vivo* validation in the mouse

To characterize the performance of SCOPe for both stimulation and imaging, we performed experiments in transgenic GCaMP6f mice with expression throughout the cortex (see Methods). We performed a 6×6 *mm*^2^ craniotomy (**Fig. S20**) and placed the SCOPe device face-down over somatosensory cortex (SSC) and visual cortex (VC) (**Fig. 4**A). To generate activity with known spatiotemporal dynamics, we implanted flexible electrode shanks (see Methods and **Fig. 4**B) in the SSC down to Layer 5 to deliver 20-100 μA amplitude, biphasic, 200-μs electrical microstimulation pulses directly beneath the imager that are known to produce reproducible neuropil induction as determined by two-photon (2p) microscopy(*54*). Since the SCOPe device is a poor fit to the mouse brain which is relatively small, we only studied a 2.1 × 3 mm portion of the array with six μLEDs, at locations 07B, 08B, 09B, 10B, 13B, and 14B (**Fig. S8**, **Fig. 4**C).

**Fig. 4.**
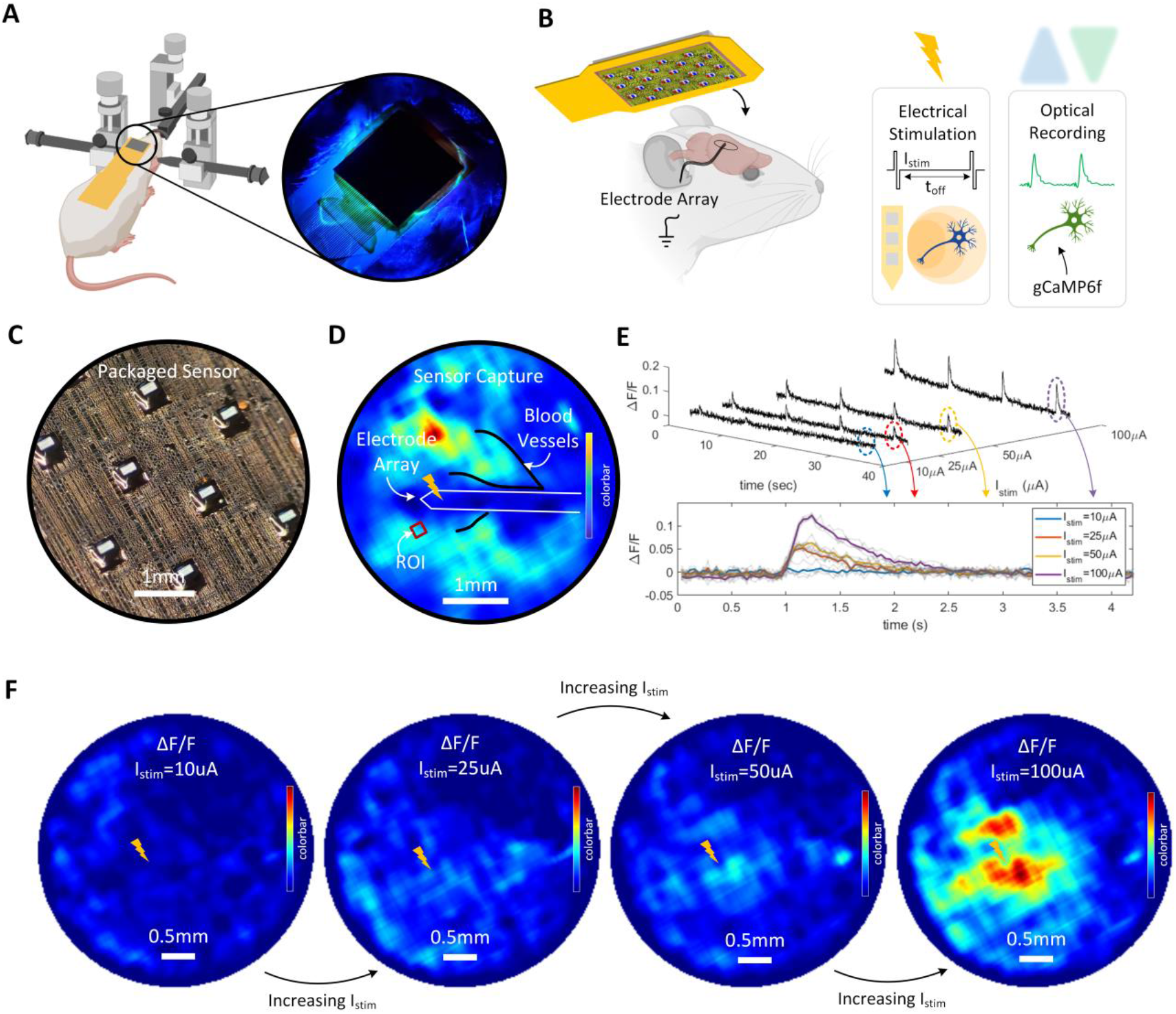
*In-vivo* imaging of calcium dynamics with electrical stimulus in the mouse model. (A) Experimental setup with the craniotomy and the SCOPe device placed face-down on the cortical surface. (B) *In-vivo* optical recording with electrical stimulus using a flexible polyimide electrode recording shank inserted into the cortical tissue. (C) Region of the SCOPe device with μLEDs, black underfill epoxy and computational mask that is in contact with the mouse brain. (D) Averaged SCOPe capture at a reconstructed image depth of 100 μm with blood vessels and stimulation sites outlined. The electrode array is laminated across the surface of the brain with the stimulation penetrating electrode sites inserted at depth located at the yellow lightning bolt. Color bar is 0-1023 counts. (E) ΔF/F averaged over the ROI denoted by the red rectangle in (D). The inset shows the responses to different current amplitudes of 10, 25, 50, 100 μA, (F) ΔF/F images reconstructed at a depth of 100 μm temporally averaged over 0.5 seconds of the first stimulation pulse. Color bar is 0-0.2 ΔF/F.

We imaged using the SCOPe device at 40 fps (**Fig. 4**D) while delivering electrical microstimulation pulses. Images are reconstructed at a depth of 100 μm. The microstimulation site aligns with LED pair location 10 (**Fig. S8**). Vasculature is evident in the sensor capture since blood vessels act as absorbers in the fluorescent scene (**Fig. S21**). We took these measurements with an output optical power from each of the blue μLEDs of 27 μW (and an electrical input power of 370 μW). Total chip power consumption was less than 10 mW, with the SPAD array consuming only 210 μW. The dynamics of neuropil response are measured by SCOPe and are synchronized with applied pulses. For 10-μA stimulation pulses, induced calcium transients are below the noise floor; increasing the stimulation amplitude between 25 and 100 μA results in responses that are directly proportional to the current amplitude (**Fig. 4**E). The 40-fps frame rate is sufficient to resolve the resulting GCaMP6f dynamics. This fluorescence response spatially varies over the full area of the cranial window imaged by SCOPe (**Fig. 4**F). A ΔF/F increase of 0.2 associated with stimulation is observed directly over the electrodes, near to LED site 10 (**Fig. S8**). This response decreases with increasing distance from the stimulation site and is undetectable at distances greater than 1.2 mm (see **Movie S1**).

The integration of two μLED colors on SCOPe allows us to use μLEDs for optogenetic stimulation concurrent with imaging. To characterize the optical power densities require for optogenetic stimulation from SCOPe, we prepared a ChR2(*55*) transgenic mouse with two craniotomies (**Fig. S22)**, one in each hemisphere (3 mm cranial window) and placed SCOPe device with one μLED activated on the surface of the right hemisphere (**Fig. 5**A). In each of the craniotomies we implanted flexible shanks, each containing 10 electrodes with pitch of 100 μm. We set the μLED optical power density to 20 mW/mm^2^ such that the power at the most-superficial electrode 50 μm from the surface in the left shank is approximately 60 mW/mm^2^ due to fluence build-up and at the deepest electrode 1 mm from the surface, approximately 5 mW/mm^2^ due to power spreading, scattering, and absorption. Neural recordings revealed the highest modulation at the most superficial site and a significant response as deep as 1 mm (**Fig. 5**B). Local field potentials (LFPs) measured on the most superficial electrode varied with μLED power (**Fig. 5**B) and differed from spontaneous activity under deep anesthesia, which is primarily in the alpha and beta (1-10 Hz) frequency bands. We observed a major increase in both frequency bands after delivering optical stimulation that is proportional to the applied emission power. On the right side (control), the time-frequency response for the second shank 3 mm away from the stimulation site shows only minor modulation, confirming stimulation selectivity. Additionally, in a separate experiment containing a wild type mouse preparation, we show that there are no photovoltaic effects associated with the optical stimulation (**Fig. S23**).

**Fig. 5.**
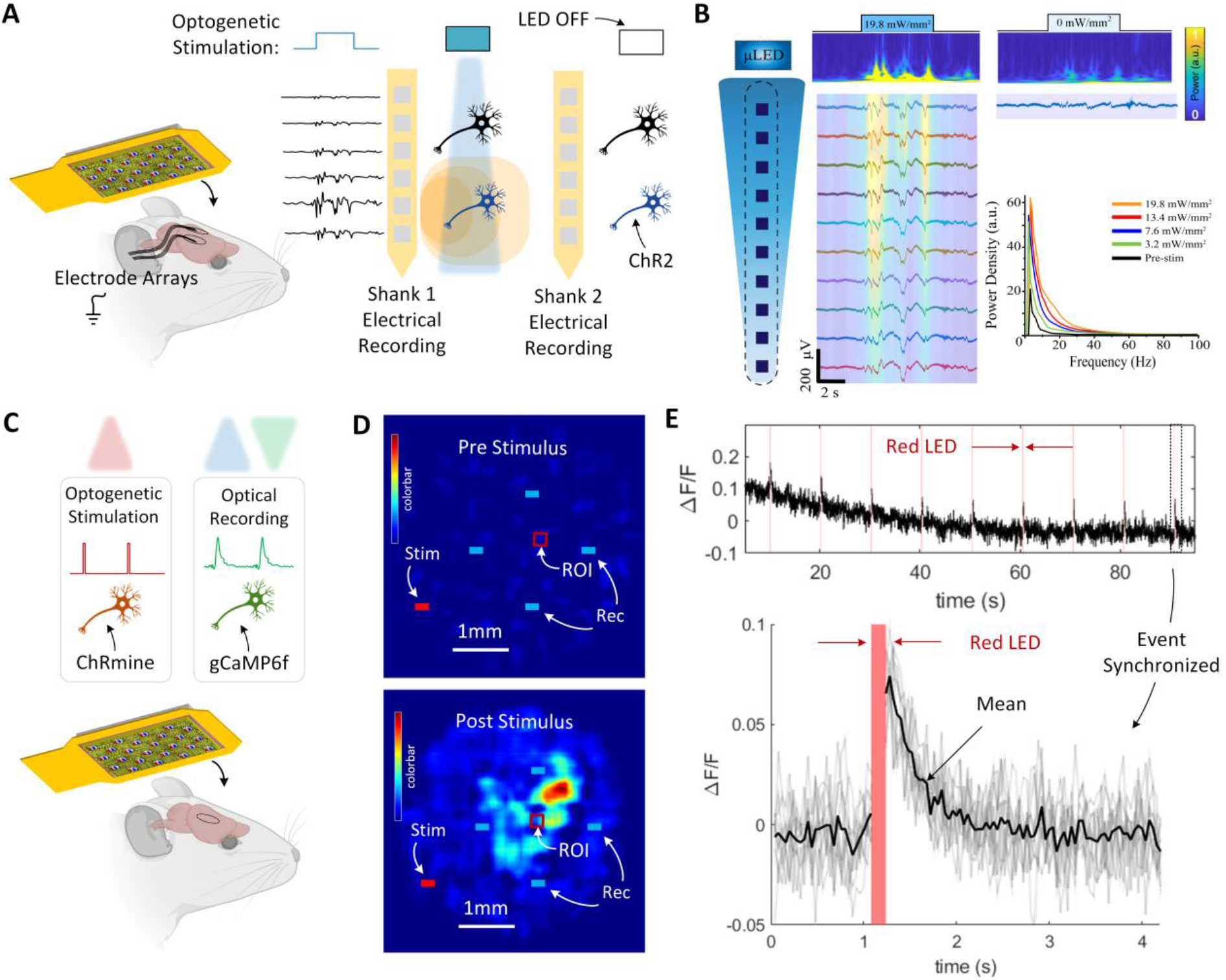
All-optical *in-vivo* imaging of calcium dynamics with optical stimulus in the mouse model. (A) Optical stimulus and electrical recording setup with two craniotomies performed in each hemisphere in a ChR2 transgenic mouse to allow a single SCOPe device to interact with both hemispheres. (B) Electrophysiological recordings on implanted electrodes showing modulation of activity in the left hemisphere underneath the blue stimulation μLED. (C) These are GCAMP6f transgenic mouse injected with ChRmine opsins. Optical stimulation and optical recording setup using a single 6×6 mm^2^ cranial window. (D) ΔF/F image capture reconstructed at a depth of 100 μm pre- and post-stimulus. Color bar is 0-0.2 ΔF/F. (E) ΔF/F time traces over a 100 second time window with the red LED stimulation frames highlighted for a total of 9 stimulation cycles. The time trace is taken by averaging the pixels in the ROI denoted by the red rectangle in (D). The inset shows the synchronized and overlaid time traces drawn with transparent gray lines over a four-second window for each of the red LED stimulation pulses. The mean of the nine time traces is denoted by a black line.

To demonstrate the all-optical bidirectional capabilities of SCOPe, we studied the same GCaMP6f mice with the red-shifted opsin ChRmine(*56*) injected in the motor cortex. We positioned a SCOPe device with dual wavelength emission capability at 590 nm and 470 nm over the cortical surface (**Fig. 5**C). For each test cycle, we applied stimulation pulses for four frames (~100 ms) from LED 08R and imaged neural responses for 400 frames (~10 s) with illumination from LEDs 10B, 13B, 14B, and 16B (**Fig. S8**). The red μLED emits an optical power of 193 μW with an input power of 12.25 mW (1.6% EQE). The optical power density at the site of the red μLED is 9.65 mW/mm^2^, which, according to the Monte Carlo model (**Fig. 3**B), provides a half-sphere with a radius of approximately 1 mm of power density > 1 mW/mm^2^ in neural tissue. This is greater than the activation threshold for ChRmine (0.1 mW/mm^2^)(*46*). While red-shifted in spectral responsivity, ChRmine still remains very sensitive to blue wavelengths (**Fig. S24** and **Section S4**). We minimized crosstalk by limiting the intensity of blue excitation light that is used for fluorescence imaging. We also confirmed responses were not due to crosstalk because the lateral distance between the red stimulation LED 08R and the midpoint between the four blue recording LEDs is 1.9 mm, which is where the greatest ΔF/F modulation is observed (**Fig. 5**D and **Movie S2**). Transients following optogenetic stimulation also had comparable time decays to those following electrical stimulation (**Fig. 5**E).

### *In vivo* validation in non-human primate (NHP)

To fully demonstrate the subdural form factor of the large FoV device and ability to decode motor behavior on single trials, we performed a sequence of behavioral experiments in a macaque monkey with a targeted expression of GCaMP8m in the dorsal premotor (PMd) and primary motor (M1) cortices (**Fig. 6**A and Methods). The SCOPe device was inserted in the subdural space between an artificial skull (**Fig. 6**B) and cortical surface. In contrast to anesthetized *in vivo* mouse experiments, experiments in NHP were performed awake and during task behavior. This gave us the opportunity to assess the ability of the SCOPe device to resolve motor function and support a brain-machine interface.

**Fig. 6.**
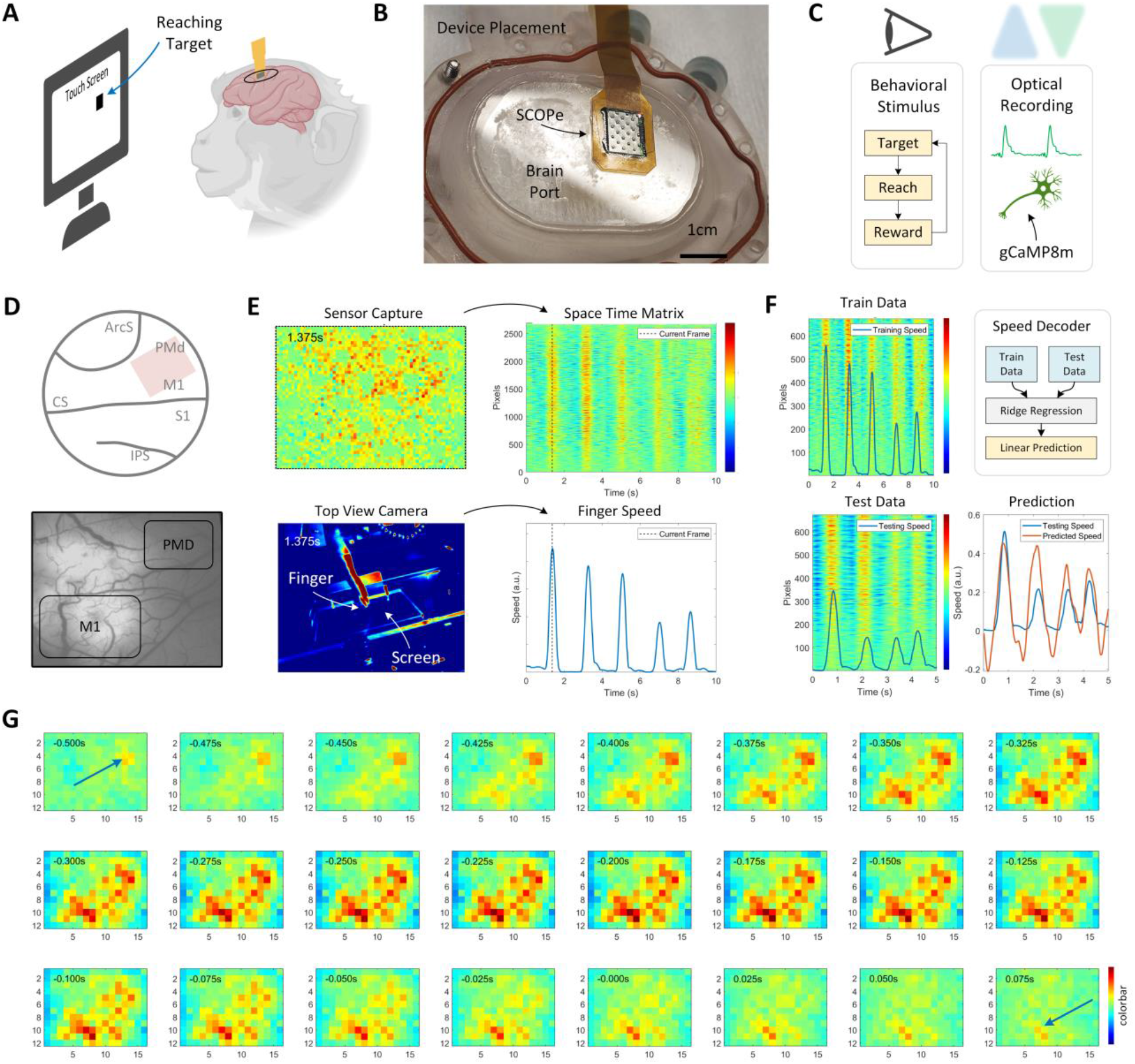
*In-vivo* imaging of calcium dynamics in the non-human primate. (A) Calcium fluorescence imaging in PMd/M1 regions of the NHP cortex. (B) Device placement on the artificial skull over a cranial window, referred to as a Brain Port. (C) Experimental setup with behavioral stimulus in which the animal touches the screen in the correct location, holds the touch for 400 ms, receives a juice reward, and retracts the arm in preparation for the next trial. During this behavior, there is simultaneous optical recording of neural activity through gCaMP8m reporters with the SCOPe device. (D) Placement of the SCOPe device in the Brain Port relative to the PMd and M1 regions. (E) Processed image capture with space versus time representation for temporal analysis of simultaneously capture infrared recordings as a ground truth of reaching activity. The finger tracking allows for direct comparison of ground truth with the sensor capture. Color bar is −1.5 to 1.5 ΔF/F. (F) Speed-feature-decoder pipeline with example training and testing datasets. The predicted velocity correlates well (*ρ* > 0.4) for randomly selected training and testing datasets swept temporally throughout the experiment. Color bar is −1.5 to 1.5 ΔF/F. (G) Spatiotemporal representations of neural activity during a reaching task showing the propagation from PMd to M1. Color bar is −1.5 to 1.5 ΔF/F.

We placed the device into a craniotomy and durotomy and directly onto the M1 and PMd (**Fig. 6**D). We sampled images at 40 fps using 22 μLEDs delivering a total optical power of 200 μW (9 μW per LED) corresponding to a power density of 450 μW/mm^2^ per μLED. We measured heat dissipation using a FLIR camera pointed at the implanted device and confirmed that the temperature did not change by more than 1°C during the experiments.

We first obtained an anatomical image focused to a depth of 200 μm to reveal the overall structure of the GCaMP8m expression and the vasculature within the 5.1 × 6.8 mm^2^ FoV (**Fig. S25**). Initial analysis of the activity revealed prominent physiological artefacts, a dominant low-frequency oscillation at 0.1-Hz associated with respiration(*57, 58*) and a 3-Hz oscillation due to the heart beat (*59*) (**Fig. S26**). We suppressed these artefacts by sequentially applying a high-pass (with a cut-off of 0.5-Hz) and smoothing with a moving average filter of 10 samples (cut-off of 2-Hz). Motion artefacts present in miniscope configurations associated with relative movement of the sensor and the brain(*60*) do not occur with the SCOPe device because it is in direct contact with the cortical surface.

We then examined the relationship between SCOPe recordings and reaching behavior. We positioned three machine vision cameras to track arm and hand movements (**Fig. S27**) and instructed the monkey to perform reach movements to visual targets presented on a touchscreen display. The sensor-wide neural activity captured with the SCOPe device clearly revealed detailed spatial-temporal dynamics. Synchronizing sensor data with behavior observed by the video revealed peaks in neural activity which correlated with reach movement speed (**Fig. 6**E and **Movie S3)**. To confirm the activity was due to reaching movements, we compared the neural activity in M1 during control recordings when the reaching arm was restrained and during a baseline period of quiet wakefulness. In each case, we observed the neural activity in M1 was inactive (**Fig. S28**). Interestingly, however, PMd remained somewhat active in the arm-restrained case, see below.

To quantify the relationship between SCOPe images and behavior, we analyzed images captured by the SCOPe device just over the region of M1 (corresponding to sensor regions in the quadrant enclosed by LEDs 14B, 16B, 17B, 19B, 20B, 22B, and 23B from **Fig. S8**) to build a feature decoder and decode movement speed trajectories during the reaching task (**Fig. 6**F). We specifically built a feature decoder for the speed of movement of the index finger of the reaching arm by estimating a ridge regression function that returns the coefficient estimates of the predictor data (see **Section S6**). We first considered the case in which a 400-sample subset of one recording (Recording 36) is used for training. We then tested the model by decoding a 200-sample subset from a different recording (Recording 39) obtained while the monkey performed the same reaching task (**Fig. 6**F). We observed good predictive capability (*ρ* = 0.66). We then swept the training and testing datasets in the selected recordings and computed the correlation to generate a two-dimensional 19 × 23 correlation matrix (**Fig. S29**). In all cases, the speed feature decoder revealed a correlation prediction of better than 0.4 (*ρ* > 0.4) swept across time in the entire recording. These results demonstrate that the SCOPe devices could successfully decode the moment-by-moment trajectories of movement speed from the motor cortices and support a brain-machine interface.

We then decoded information from spatial organization of neural activity across the entire FoV of SCOPe. To do so, we first applied a low-pass spatial filter by binning groups of 16 × 16 pixels to improve the imaging signal-to-noise ratio. For each trial, we then averaged the spatiotemporal structure of the neural response across each reach movement, 80 samples centered on an individual reaching event identified by peak arm speed (**Fig. S30**). We used ground truth white-light camera images taken of the device before and after device placement overlaid with the corresponding structural widefield fluorescence images of PMd and M1 to localize the SPAD detector imaging sites (**Fig. 6**D). Across all trials, we observed that there was a peak in movement-related neural activity across the motor cortical surface that first appeared in PMd, ~ 500 ms before movement onset. We then observed the peak propagates across the cortical surface to M1 for ~300 ms until movement offset, consistent with prior electrophysiological studies(*61*). We leveraged the large image FoV to investigate the relative timing of spatial-temporal dynamics across PMd and M1 by correlating the response of the PMd and M1 regions as a function of the time lag between them. When the monkey performed the reaching task, we observed a positive correlation at a lag of 10-time samples (250 ms) between the PMd and M1 regions. In contrast, when the monkey was arm-restrained experiments, we did not observe any correlation (**Fig. S31**). Neural activity in PMd correlated with intent to move the arm but did not propagate into M1 in the absence of movement. Overall, these results demonstrate that the SCOPe device can resolve functionally significant multiregional brain dynamics across the motor cortices to reveal the gating of communication between brain regions during behavior.

## DISCUSSION

Here, we engineer and validate an implantable optical recording and stimulating device, SCOPe, that is thin enough to be accommodated in the subdural space, creating the opportunity for new generation of brain-machine interfaces (BMIs) based on such volumetrically efficient designs. While electrode-based interfaces have achieved similar miniaturization, optical interfaces have suffered from excessive heat dissipation and lagged in aggressive miniaturization due to the need for bulky optical elements and light sources. By aggressive exploiting custom CMOS ASICs and advanced heterogenous integration. SCOPe realizes a fully implantable optical neural interface that is less than 200-μm in total thickness and generates very little heat (< 1°C) while covering a FoV of more than 5.1 × 6.8 mm^2^.

BMIs, in general, are enabled by the ability to record and stimulate neural activity as well as encode or decode sensory or motor function (*62*). We have demonstrated SCOPe device as a full-function BMI in the NHP by imaging motor function using virally mediated expression of a genetically encoded calcium indicator in the primate motor cortex and by predicting speed trajectories moment-by-moment. Because SCOPe consumes less than 10 mW in 40 fps operation while producing off-chip data rates of less than 20 Mbps, it is very amenable to the addition of wireless powering and wireless data telemetry in future iterations directly on the SCOPe ASIC, eliminating the wires through the skull required.

The SCOPe device operates in the mesoscopic regime due to its reliance on lens-less computational imaging approaches. Mesoscopic imaging offers a powerful tool to investigate and extract information from large-scale neuronal networks spanning multiple brain regions at higher resolutions than traditional brain imaging methods such as fMRI (**Section S4**). Moreover, the spatial resolution we demonstrate still exceeds surface electrode approaches, highlighting the strength of using light to mediate neural interfaces. We used AAV to transfect neurons in NHP, an approach which also allows for cell-type selectivity for projection neurons and interneurons. This selectivity is already applied clinically in gene therapy in the central nervous system, for example, to treat Parkinson’s Disease. When single-cell resolution is required, the number of cells that can be separated becomes limited by the total number of detector pixels, usually requiring sparse labelling. Single-cell approaches also benefit significantly from the use of soma-localized reporters(*63*) to reduce background neuropil fluorescence. Strategies to further increase cellular-level resolution include thin metasurfaces(*64*), computational imaging techniques such as blind-source separation(*8*) and deep-learning-enabled interpolation(*65*), and further increasing the number and density of pixels in SCOPe.

In summary, the SCOPe device leads the way to a new class of BMIs that can leverage genetically encoded reporters and actuators along with a subdural-form-factor implantable device.

## Acknowledgments

We gratefully acknowledge TSMC for chip fabrication and their support in the use of experimental SPAD devices. All schematics were created using BioRender.

## Funding

DARPA under Contract N66001-17-C-4012 (VAP, KLS)

National Science Foundation under Grant 1706207 (KLS)

NIH NS-103518 (BP)

Circuits and Systems Society Fellowship (EHP)

## Author contributions

Conceptualization: EHP, KLS

Circuit design and thinning: EHP, SM, YG, AB

Optical packaging: EHP, HY, SM

Computational imaging algorithms: VB, AV, JR

*In-vivo* mouse experiments: EHP, HY, IU, VAP

*In-vivo* NHP experiments and neural data analysis: EHP, AD, KEW, BP

NHP surgery: BP, JSC

Funding acquisition: VAP, KLS, BP

Supervision: KLS

Writing – original draft: EHP

Writing – review & editing: EHP, BP, KLS

## Data and materials availability

All imaging data, all scripts used for image processing, and the decoder model are available at https://github.com/klshepard/scope. All other relevant data are available from the corresponding authors upon reasonable request.

## Supplementary Materials

Materials and Methods

Supplementary Text

Figs. S1 to S30

Tables S1 to S4

References (*##*–*##*)

Movies S1 to S3

## Supplementary Materials

### SUPPORTING INFORMATION

Supporting information includes **Fig. S1** through **Fig. S31**, **Table S1** through **Table S5**, Supplementary **Movie S1** through **Movie S3**, and extended methods and Supplementary **Section S1** through **Section S6**.

### ACKNOWLEDGEMENT

This work was supported in part by DARPA under Contract N66001-17-C-4012, by the National Science Foundation under Grant 1706207 and by the NIH Brain Initiative under U01-NS103518.

### AUTHOR CONTRIBUTIONS

E.H.P and K.L.S. conceptualized the study. E.H.P, S.M., and Y.G. designed the circuits. E.H.P, H.Y., I.U, and V.A.P performed the *in-vivo* mouse experiments. E.H.P, H.Y., A.D., and E.K.W performed the in-vivo NHP experiments. A.B. performed the CMOS die-thinning. V.B, A.V., and J.R. wrote the computational imaging algorithm. B.P. and J.S.C. performed the surgical procedures in NHP. E.H.P., A.D., and B.P. analyzed the neural activity in NHP. E.H.P, B.P. and K.L.S. wrote the paper and edited the manuscript. K.L.S. provided overall supervision and guidance.

### CORRESPONDING AUTHORS

Correspondence and requests for materials should be addressed to K.L.S.

### Materials and Methods

#### Application-specific integrated circuit design and fabrication

This chip was designed in a custom mixed-signal design flow using Mentor and Cadence tools. The analog frontend was designed and simulated using Virtuoso Schematic Editor (Cadence) and Virtuoso ADE (Cadence) simulation suite with the SPAD behavior modeled using experimentally verified Verilog-A models. Post-layout verification was performed with Calibre Design Rule Check (nmDRC, Siemens EDA) to ensure that the design meets foundry specifications, Calibre Layout-vs-Schematic (nmLVS Siemens EDA) to cross-reference the layout with the schematic netlist, and Calibre Resistance-Capacitance Extraction (xRC, Siemens EDA) to generate a netlist including RC parasitics and model layout-related non-idealities. The digital register-transfer logic backend was written in Verilog HDL and simulated using NC Verilog (Cadence). The gate-level netlist is synthesized using Design Compiler (Synopsys), and place and routed level netlist is generated using Encounter (Cadence). The digital logic and timing are simulated and checked against various testbenches after each of these steps to ensure functionality. The generated GDSII stream format (GDSII) is streamed out and includes the digital core interfaced with the analog imager array. Final chip-level simulations and nmDRC, nmLVS, and xRC checks are performed before exporting the GDSII to foundry for fabrication.

The chip is fabricated in a 0.13-μm bipolar-CMOS-DMOS (BCD) technology at a commercial semiconductor foundry (Taiwan Semiconductor Manufacturing Company, Hsinchu City, Taiwan). This technology also supports integration of the SPADs. The chips are also solder-bumped for both interconnection from the chip to the flexible PCB and for mounting of the μLEDs. Chips are initially thinned to 305-μm thickness and diced for assembly in SCOPe.

#### Package and board design

The boards used for transmitting data from the chip to the host computer consistent of a rigid host board and a flexible chip board. The host board contains an FPGA module (XEM7010, Opal Kelly) for programming the chip and relaying the data coming back over USB to the host computer for storage. The Opal Kelly integration module has a 5V power jack which is used to generate all the required chip voltages using low-drop-out (LDO) linear regulators. The SPAD voltage of ~21V is supplied using an external Triple Output Programmable DC Power Supply (E36312A, Keysight Technologies) with a BNC connector. The power-on sequence turns on this voltage last to prevent any damage to the low voltage digital logic on the chip.

The flexible polyimide chip PCB is designed in electronic design automation software (Altium Designer, Altium Ltd) and manufactured at a commercial printed circuit board electronics manufacturer (EPEC Engineered Technologies). The full stack up of the one-layer flex PCB includes coverlay, coverlay adhesive, copper (conduction layer), and polyimide for a total package thickness of 52μm (**Fig. S1**). Stiffener is present on one end of the board to form a 200-μm-thick zero-insertion-force (ZIF) connector to the host board. The copper layer within this connection edge was finished with electroless-nickel-electroless-palladium-immersion-gold (ENEPIG). To accommodate the chip, there is a laser-cut opening in the polyimide substrate such that the face of the integrated circuit is exposed when bonded to the board’s back side. The chip is assembled as a 305-μm-thick die using a pick-and-place assembly tool (Finetech, Fineplacer Lambda, Berlin, Germany). The pick-and-place tip (4.8mm ×4.8mm ×0.5mm) used for bonding the CMOS chip to the flex PCB was chosen to approximately match the size of the die. We use 0.2mm washer to match the 0.5-mm thickness of the tip. The process program pre-heats the flexible PCB to 100 °C prior to alignment to account for warping of the flexible PCB during the temperature ramp. The pick and place tip temperature is ramped to peak temperature of 250C, which allows for the reflowing of the lead-free solder bumps on the CMOS die (**Fig. S1**).

**Fig. S5** shows the packaging flow for SCOPe, including the integration of the μLED light sources, filters, and computational mask onto the CMOS substrate. To prepare the chrome-polyimide computation imaging mask, a two-inch silicon wafer was spin coated with two layer of low-stress thick polyimide (PI-2611, HD MicroSystems™). After curing in a N_2_ oven under 350 °C for 30 minutes with 2 - 4 °C/minute ramp, a 20-μm-thick polyimide film results. A 200-nm-thick chromium (Cr) mask was then electron-beam-deposited on top of this film by using an Angstrom EvoVac Multi-Process Evaporator. A 1-μm-thick S1818 photoresist was then spin coated on top of the Cr film following by 1-minute soft bake at 115 °C. After 30-minutes of baking, the S1818 photoresist was exposed using a Suss MA6 Mask Aligner with a dose of 150 mJ/cm^2^ and developed with AZ300MIF for one minute. The sample was then soak inside Transene Chromium Etchant Type 1020 for two minutes to wet etch the computational pattern into the Cr layer. Finally, after removing the photoresist, a 273-nm ArF Excimer laser (IPG Photonics) was used to cut out the mask from the substrate and make the opening for the μLEDs for SCOPe packaging. **Fig. S5** outlines the process steps which (A) start with the polyimide (25-μm thick) substrate fabrication, followed by (B) patterning of the Cr amplitude mask and cutting (C) of both the μLED openings and the over mask using the excimer laser.

In parallel, (D) we start with a CMOS (305-μm thick) chip with exposed solder bumps for the μLED bond pads, then (E) deposit a thin-film 500nm long-pass interference filter (11-μm thick) with the μLED bond sites protected, and finally (F) flip-chip bond the μLEDs (80-μm thick) using a Eutectic solder reflow process. Combining the two processes, (G) we align the polyimide substrate with computational mask with the μLED cutouts to the CMOS chip using pick-and-place tool and bond the two substrates together using our custom-made 500nm long-pass absorption filter epoxy (~20-μm thick). After curing, (H) we use a Model PLI-11A Pico-Injector (Warner Instruments) to underfill the μLEDs with EPOTEK 320-lv black optically absorbing epoxy. When all the packaging steps are completed, (I) the chip is die-thinned (~7.5-μm thick) to become mechanically flexible.

Die thinning is done on the X-prep Precision Milling/Polishing system (Allied High-Tech Products). Initial thickness measurement of the chip is done by using the X-Prep Vision – Substrate Measurement Instrument which uses spectroscopy to find the bulk silicon thickness of the chip, a process which is repeated after each processing step. The backside of the packaged SCOPe device, when first placed into the X-prep milling system, is leveled, and calibrated to the grinding mount. The grinding mount has an attached diamond grinding disc which grinds down the excess silicon to ~5μm greater than the desired thickness. This grinding step is done in multiple passes to remove excess debris. On each grinding step, grinding fluid is used to lubricate the grinding surface and to remove heat generated from the mount. After this initial grinding is done, the chip backside sees three polishing steps at grit sizes of 15μm, 1μm, 0.04 μm. The 15-μm and 1-μm steps use diamond grit paste and a polishing mount to smooth the surface of the chip. The 0.04-μm step uses silica as a finishing polish for the chip. These three polishing steps remove approximately 5 μm to reach the target silicon thickness of 10 μm. After thinning, the front-side and back-side of the SCOPe device is coated with ~5-μm-thick parylene to passivate the device and provide additional biocompatibility (**Fig. S6**).

Once thinned, the bending stress in the CMOS die can be modeled using thin plate theory since the thickness of the CMOS die is orders of magnitude smaller than the lateral dimensions(*66*). In this case, the minimum radius of curvature (*R_min_*) before the stresses in the CMOS die exceed the maximum bending stress (*σ_max_*), resulting in fracture, is related to the biaxial Young’s Modulus, the Poisson’s ratio (v), and the thickness of the substrate by:

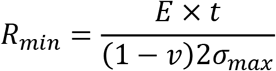

#### Data acquisition interface on the host board

The host board is responsible for all data acquisition, including capturing, streaming, and storing the data from SCOPe on a host computer. The interface with the data stream from the chip, the host board uses the XEM7010-A50 (Opal Kelly) integration module both to program the chip and to convert the data coming from the chip into USB 2.0 format for packet transfer back to the host computer. The XEM7010-A50 module supports 2.7Mb of on-chip FPGA block RAM, 512 MiB DDR3 memory banks, switching power supplies, a multiple-output phase-locked loop (PLL), and up to 120 input-output (IO) connections. FPGA configuration bits are generated and compiled in Vivado Design Suite (Xilinx). The top-level hardware design logic (HDL) is programmed using Verilog and contains the top-level state machine for communicating with both the chip and the FrontPanel HDL for communicating with the Opal Kelly FrontPanel software API.

Using this FPGA interface, the SCOPe ASIC is initialized and programmed using a scan chain, which is ty enabled at chip initialization and whenever the state configuration of the chip needs to be updated. Since the scan chain is a serialized shift register, the amount of time needed to program the chip depends on the number of configuration bits. Data transmission from the SCOPe ASIC is serialized at 100 MHz with three off-chip signals: DATA, CLK, and FLAG. DATA is the raw sensor data with a serialized stream out; CLK is used for synchronizing the transmitter and receiver clock, and FLAG is a logical enable signal used for determining the validity of the data. The effective data rate for this link is the determined by the number of pixels, the bit depth per pixel, and the frame rate: 192 × 256 × 10b × 200Hz = 98.3Mb/s. For the USB2.0 protocol, the Opal Kelly converts this serialized data stream into a 16b wide bus for use with the USB 2.0 pipe out. The conversion is done in two steps using Xilinx’s intellectual property (IP) FIFO Generator with a pass-through 1:8 FIFO follows by an 8:16 FIFO. The pass-through FIFO converts the serialized data stream into an 8b parallel bus and operates on the chip CLK domain. The second FIFO buffers the data as it is streamed to the host computer and operates on the clock domain of the USB 2.0 interface (48 MHz). Chaining FIFOs in this manner works only if the secondary clock is faster than the data clock, such that the FIFOs do not fill. The data is read little-Endian off the chip and bit-reversed prior to sending over USB to simplify data visualization. The data is stored in binary files on the host computer and is read and reformatted into images with 10b precision in MATLAB.

#### SPAD electrical and optical characterization

A single photon avalanche diode (SPAD) is a semiconductor p-n junction with a reverse-bias (*V_SPAD_*) exceeding the breakdown voltage (*V_BD_*) by an amount referred to as the excess voltage (*V_E_*), *V_SPAD_* = *V_BD_* + *V_E_*, such that a single photogenerated carrier initiates an avalanche current through impact-ionization. Upon triggering, the voltage across the SPAD is reduced below *V_BD_* to quench the avalanche and subsequently returned to *V_SPAD_* to detect another photon. To determine the best available SPAD to use as our detector, a design-of-experiments (DOE) was performed with many SPAD variants. The SPADs in this array varied the guard ring structure (standard p-well, high-voltage p-well, virtual p-epi) as well as the depth of the implant region (p+, high-voltage p-body, p-well, high-voltage p-well). The breakdown voltage of each of these devices was determined by performing a current-voltage (IV) sweep using a B1500 Semiconductor Parameter Analyzer (Keysight, USA). PDP measurements were performed by sweeping the wavelength on a Digikröm CM110 monochromator fed into the 819D Series Spectralon integrating sphere, allowing for us to simultaneously measure the incident power illuminating the SPAD with a 1936-R single channel optical power meter (Newport Corporation, USA).

The photon detection probability is calculated as:

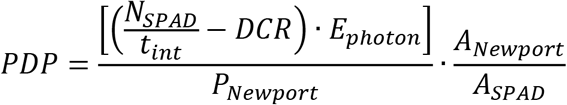

where *N_SPAD_* is the number of SPAD counts collected within an integration time *t_int_, DCR* is dark count rate,, *E_photon_* is the energy of a photon, *P_Newport_* is the photodiode power, *A_Newport_* is the cross sectional active area of the photodiode, and *A_SPAD_* is the cross sectional active area of the SPAD.

#### Flexible electrode shank fabrication

The polyimide electrode shanks used in the mouse studies were custom fabricated. A three-inch fused silica wafer was coated with 2-μm-thick of parylene-C using chemical vapor deposition in a SCS Parylene Coater. For lift-off processing, a 300-nm-thick film of S1811 was spin coated on the parylene substrate and cured at 110°C for one minute. The wafer was exposed to ultraviolet light using a Suss MA6 Mask Aligner and developed with AZ300MIF developer for 30 seconds. A 10-nm-thick Ti adhesion layer, followed by a 100-nm-thick Au layer, was deposited on the patterned photoresist by using an Angstrom EvoVac Multi-Process Evaporator. The resultant wafers were soaked in REMOVER PG (Kayaku Advanced Materials, Inc., Westborough, MA) for 12 hours for lift-off. A second 2-μm-thick parylene-C film was deposited using chemical vapor deposition using 3-(trimethoxysilyl) propyl methacrylate (A-174 silane) as an adhesion promoter. A final layer of parylene-C was deposited following a spin-coating step of soap solution diluted to 1% concentration in de-ionized water. An 8-μm-thick layer of AZ9260 was spin-coated at 3,000r.p.m, baked at 115°C for 2 minutes, exposed using a Suss MA6 Mask Aligner and developed with AZ400K developer. A second dry etch process is performed in a plasma-reactive ion etcher (Oxford Plasmalab 80; 180 W, 60 sccm *O_2_* and 2 sccm *SF_6_*). The PEDOT: PSS dispersion was spin-coated on the wafer and baked at 115 °C for 30 minutes. The sacrificial layer of parylene-C was peeled off to complete patterning of PEDOT: PSS on the electrodes. The contour of the shanks was patterned by using an IPG Photonics excimer laser cutter.

#### Mouse surgery and shank insertion procedure

The Institutional Animal Care and Use Committee (IACUC) reviewed and approved protocols for Columbia University’s program for the humane care and use of animals and inspects the animal facilities and investigator laboratories. Evaluation of the implanted devices was performed in compliance with Animal Welfare and Columbia’s IACUC regulations under approved IACUC protocol AC-AABE5554 “Development of high-density, implantable recording, imaging and stimulating arrays”. All mice were acquired from Jackson Labs (The Jackson Laboratory). For experiments with optogenetics with blue excitation and electrical record, Thy1-ChR2-YFP (Tg(Thy1-COP4/EYFP)9Gfng) mice (6-12 weeks) were utilized. For all other experiments, the strain Vglut1-IRES2-Cre-D was bred with the strain Ai148(TIT2L-GC6f-ICL-tTA2)-D (or Ai148D) to generate offspring possessing both GCaMP6f expression and tTA2.

For experiments with optogenetic stimulation in the red, the CGaMP6f transgenic mice were also transfected with the opsin ChRmine. Sixteen- to twenty-four-week-old GCaMP animals were anesthetized with 1-5% isoflurane. A 0.25 mm opening was drilled on the skull taking special caution not to puncture the dura (−0.5 mm AP, 2 mm ML from Bregma). Using a pulled glass pipette as a needle (50μm tip) and a 10 μL Hamilton syringe mounted on a microsyringe pump (World Precision Instruments, Smartouch), two 500 nL injections of AAV-ChRmine (AAVdj-CaMKIIa-ChRmine-Oscarlet-Kv2.1 (titer; 7.5E+11 vg/ml) were made into the cortex, at 300 and 600μm depth. The injection rate was 80 nL/min. After injections the scalp was sutured. Experiments were carried out 3-4 weeks after injections.

For implantation, the GCaMP6f transgenic mouse was anesthetized using isoflurane with an induction level of 3% and driven by an oxygen flow at the rate of 2 liter per min. Upon induction of anesthesia, the animal was removed from the induction chamber and placed on a stereotactic frame which is attached to a nose cone to maintain a steady flow of isoflurane. The body temperature of the mouse was controlled using a feedback-regulated rodent warmer (ATC-2000) pad set at 32°C. 0.2 ml of bupivacaine was injected subcutaneously under the scalp prior to removal. The skull over V1 was drilled in a 6-mm-diameter circular shape by a dental drill and gently removed. Dura mater was incised using a fine-tip scalpel. The SCOPe device was positioned over the cranial opening directly on the pial brain surface as shown in **Fig. 4**A. For experiments involving either electrical stimulation or record, the flexible polyimide shanks were inserted into cortex before the SCOPe device was positioned.

#### Data analysis of SCOPe data in mouse experiments

In all experiments, a hot pixel mask is applied to remove the hot pixel that occur due to manufacturing defects in the SPAD sensor. In **Fig. 4**D, the averaged sensor capture is the mean of all sensor capture frames, and the image is cropped and rotated to align with the craniotomy viewing in **Fig. S20**B. In **Fig. 4**E-F and **Fig. 5**D-E, a logical mask is used to remove the high variance pixels that have standard deviation counts that are more than 3 × the mean standard deviation which occur due to insufficient filtering or black epoxy adjacent to the LEDs. The ΔF/F is calculated by normalizing to the pixelwise median where N is the raw counts matrix with dimensions 192 × 256 × n, with n representing the sampled time dimension.

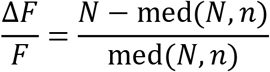

In **Fig. 4**F, **Fig. 5**D, **Movie S1**, and **Movie S2** for visualization purposes, the pixels are spatially binned in 4 × 4 bins. In **Fig. 4**F and **Movie S1** the image is cropped and rotated to align with the craniotomy view in **Fig. S20**B.

#### NHP surgical approach

One adult male rhesus monkey (*Macaca mulatta*) participated in the experiments (Monkey T, 11.4 kg). A large-scale craniotomy and durotomy (4 cm × 2 cm) were made over the frontal motor cortices of the left hemisphere under propofol anesthesia. After recovery, more than 4 weeks following window implantation, pGP-AAV1-hSyn-jGCaMP8m-WPRE (25 uL, AddGene) was then delivered to a single site in the dorsal premotor cortex (PMd) using convection-enhanced delivery (CED)(*67*). All surgical and animal care procedures were done in accordance with National Institute of Health guidelines and were approved by the New York University Animal Care and Use Committee.

#### Behavioral task and data processing

After functional expression was confirmed using fluorescence microscopy, approximately after two months following injection, the SCOPe device was placed on a window insert that spanned PMd and M1 cortices and placed directly onto the cortical surface within the subdural space (**Fig. 6**B). The monkey was then instructed to perform a reaching task to test the ability of the device to measure neural responses to behavior. Each trial of the task started with a green square (2×2 deg) presented on a vertically oriented LED display and touch-sensitive screen (ELO Touch Solutions Inc) placed in front of the monkey while seated in the primate chair. The monkey was trained to reach and touch the target for 400 ms to receive a fluid reward. The reach target was then extinguished and illuminated at another location to instruct another reach movement.

Videos from three machine vision cameras (1024×768 pixels, FLIR) running at ~ 60 Hz were used to localize the 3D position of wrist and distal finger relative to a calibrated origin on the primate chair. Signals were synchronized using a fiducial pulse that was broadcast and recorded together with the video data (Robot Operating System). Camera extrinsics were measured from a checkerboard presented prior to task performance and camera calibration was computed from camera extrinsics and intrinsics using open-source code (OpenCV). Camera 1 provides a viewing angle from the top with the x-direction corresponding with left-to-right arm movements and the y-direction corresponding with forward-to-backward reaching movements. Camera 2 provides a viewing angle from the right with the x-direction corresponding with forward-to-backward arm movements and the y-direction corresponding with up-to-down reaching movements. Camera 3 provides a viewing angle from the left with the x-direction corresponding with forward-to-backward arm movements and the y-direction corresponding with up-to-down reaching movements. We then used an open-source deep learning toolbox (DeepLabCut(*68*)) to track the reach movements during task performance. The wrist, fingertip and landmarks on the primate chair were manually labeled on a subset of frames from each camera and used to train the model. The remaining frames were then labeled automatically from each camera by the trained model.

#### Data analysis of SCOPe data in NHP experiments

Neural data analysis was performed in MATLAB (MathWorks). The hot pixel from the image capture was removed with digital Hadamard mask; since the hot pixel is intrinsic to the pixel sensor in the array, this same hot pixel mask can be used for all the recordings, including dark images. High variance pixels whose activity had a standard deviation greater than twice the mean, due to high μLED leakage and insufficient filtering due to minor defects in the packaging, were removed. We then spatially binned the data by computing the mean of adjacent pixels for a certain number of bins. Initial analyses determined that spatial binning could be done across up to 16×16 pixels before the pixel activity was corrupted by the uninformative LED blocks. After spatial binning, we then high-pass filtered the activity using a high-pass function with normalized passband at 0.5-Hz at a signal sampling rate of 40-Hz. We then temporally-smoothed the data using a moving average filter of 10 samples.

### Section S1. Chip design and packaging

#### ASIC design

The imager is 5.1 mm × 6.8 mm and is comprised of a 3×4 array of macros, each consisting of 14 blocks of 16×16 SPAD pixels with a pitch of 25 μm, and 12.5% of the array is removed for μLED bond pads and drivers. Unlike an earlier design(*19*) in which detected photon counts are stored in shared-row counters, this design(*69*) supports 10-bit pixel-level counters and a global shutter, improving the ratio of integration time to frame period 100-fold from 0.625% to 99.6%, while supporting frame rates of up to 200 fps with a 100 MHz reference clock. The chip’s frame rate is 200fps, when reading the entire array, or 400fps, when reading half the array (128×192); the latter allows for the imaging of fast state-of-the-art voltage indicators(*70*). Each pixel consists of two passively quenched SPADs whose outputs are combined into an in-pixel 10b counter. The data is read off the imager in a rolling column readout structure where the pixel data is buffered with up to 16 repeater cells throughout the array and loaded into a data serializer for streaming off chip. The data transmission module is a 1920b:1b shift register based serializer.

#### Flip-chip CMOS packaging

To be compatible with a flip-chip bonding process, the chip wire bond pads modified via a re-tape out (RTO) to convert to flip chip bond pads via a change in the back end of line (BEOL). The wire bond BEOL stack up includes a passivation via hole between the top metal and the aluminum pad layer as well as a top passivation opening to expose the aluminum pad. The IO pads for digital and supplies are along one dimension of the chip, and there are additional support bump pads on the remaining three sides of the chip. This is an unconventional method of using the flip-chip BEOL methodology, since normally the bumps are equally spaced at high density across the entire die, to get low inductance signal and power IO. However, since the center of our die is an imager that needs to be kept open for sensing, there are only peripheral IO bumps, approximating that of a wire bond packaged die.

#### SPAD architectures: time-gated versus photon-counting

The simplest SPAD quenching circuit is given with a passive quench transistor. The bias transistor operates as a large resistor, discharging the anode of the SPAD until there is a photon-induced avalanche current triggering a pulse that can be registered as a count in the external circuitry. In first implementations (*18, 19*), we designed a custom SPAD-based image sensor to enable time-gated (TG) fluorescence imaging through single-photon counting. Time gating uses the design of a high-speed active-quench, active-reset pixel circuit, which necessitates the implementation of a rolling shutter, shared off-pixel counter in order to maintain a small pixel pitch. In gated operation, the imager is only activated after each excitation pulse. Gating the sensor provides additional background rejection of scattered excitation light. The efficacy of time-gating, however, is limited in practice by a) the turn-off time of the excitation pulse, b) the fluorescence lifetime, and c) the impulse-response function (IRF) of the SPAD detectors. These photogenerated carriers diffuse, or random walk, until they reach the avalanche region of the SPAD causing a much delayed detected photon count(*71*). The minority carrier lifetime in silicon substrate(*72*) is long compared to the nanosecond timescales required during time-gating operation of the SPAD. The ineffectiveness of time-gating to do time-domain filtering of the blue excitation light result necessitates the development of high-quality optical filters.

### Section S2. Modeling of fluorescent scene

#### Modeling light propagation in tissue

We estimate the light propagation in tissue using a combination of analytical solution and Monte Carlo solver(*43*) modeling the light path geometry. The analysis starts from the detector since the sensitivity of the pixel sensor informs the required μLED light excitation. For a chief-ray angle (CRA), *θ*, of the SPAD pixel photodetector, the free-space path loss of an isotropically emitting fluorescent disk scene (**Fig. S11**) with radius of *R* = *d* tan(*θ*) is given by the equation: 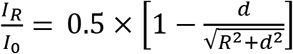. With a CRA of 70°, the total normalized radiated emission from the fluorescent disk captured by the sensor pixel is 0.327. The Monte Carlo simulation uses a modified absorption coefficient of μ_a_=0.21mm^−1^ to model the amount of absorbed blue light due to the presence of a dense fluorescence volume. At a tissue depth of 200 μm, the μLED diagonal spacing of 0.6 mm (**Fig. 3**B) produces an illumination intensity at the mid-point between four μLEDs that is 75% of the normalized μLED power density (mW/mm^2^) as a result of Lambertian emission, angular spreading, tissue scattering, and absorption. The peak fluence values close to the light source locations and near to the tissue interface result in a power density gain due to the excess scattering events. The diffuse optical property of the tissue causes the blue excitation light to scatter back towards the CMOS sensor at approximately 7.5% of the normalized illumination intensity.

#### Fluorescent conversion efficiency

We use literature values for the biophysical properties of the cells to determine the conversion from blue light excitation to green fluorescence emission. First, we determine the amount of spectral overlap, *S_B→gCaMP6f_* = 0.6, between the blue LED emission and the gCaMP6f absorption(*73*) (**Fig. S14**). In this fluorescent disk, the neuronal cellular density available for fluorescence tagging in the mouse visual cortex is approximately 96% (84% neuropil, 12% somas, and 4% blood vessels)(*74*). Over-expression of gCaMP6f can result in damaging side-effects, so moderate to high expression can result is a total overall expression of 45% of all cells(*75*). Combining the cellular density with high expression results in an overall fluorescence of *D_n_* = 43.2% per unit volume of tissue. At high expressions, the neocortical intracellular gCaMP6f concentration, c, in transgenic mouse models is 2-10 fold lower than typical conditions of AAV infections resulting in an intracellular concentration on the order of *c* = 40*μM*(*75*). The ratio of absorbed blue excitation light is then determined by the extinction coefficient(*76*), *ϵ_c_* = 5030 *M*^−1^*mm*^−1^ and application of the Beer-Lambert formula: 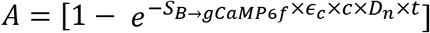 resulting in *A* = 0.5% for a volumetric slice of cortical tissue with *t* = 100*μm*. The amount of green fluorescence signal generated by this fluorescence disk includes the fluorophore quantum yield(*76*), *Q_Y_* = 0.59, resulting in a total blue-to-green conversion efficiency, *η_Blue→Green_* = *A* × *Q_Y_* = 0.003. Cascading the loss due to light path propagation and biophysical blue-green light transduction results in an overall end-to-end fluorescence signal detection chain efficiency given by the following equation:

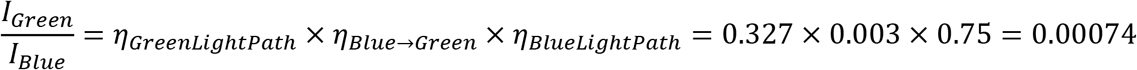

### Section S3. Computational Imaging

#### Texas two-step amplitude mask

Due to mask separability, the local spatially varying point-spread function (PSF) can be fully represented by a superposition of orthogonal terms (**Fig. S17**), where *P_od_* and *P_cd_* operate on the rows of *X_d_*, and *Q_od_* and *Q_cd_* operate on the columns of *X_d_* (the subscripts o, c, and d refer to “open”, “coded”, and “depth”, respectively). As a result, we can capture from a single shot, *Y*, of a superposition of 2D scenes, *X_d_*, which can be computationally “refocused” to retrieve each of the depths.

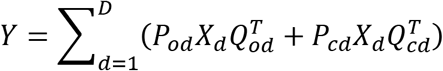

With the calibration matrices *P_od,cd_* and *Q_od,cd_* and a raw image sensor capture, *Y*, the scene is computationally reconstructed (**Fig. S18**), by solving a regularized least-squares problem. where M is a 192×256 binary matrix representing the μLED blocks and ⊙ is the Hadamard product.

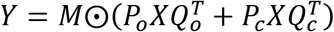

While we were able to achieve good results using high-contrast scenes such as fluorescent beads and double line slits, the amplitude mask is not optimized to work with low contrast biological scenes. Future implementations can head towards phase masks with optimized PSF contours and high-contrast features to provide good protection against noise amplification in these low-contrast scenes and in scattering media similar to epifluorescence imaging(*17*); nevertheless, improvements must be made to further reduce the working distances to allow for fully implantable operation.

#### Imager calibration and reconstruction

The imager is calibrated using a 30-μm crosshair line slit etched into a 100-nm thick Cr deposited on a 100-μm thick glass wafer. The line slit is illuminated, in transmission mode, with a green LED (ThorLabs #M530L4) through an 80° Diffuser (Edmund Optics #47-678) to approximate an isotropic fluorescent source. The line-slit mounted onto an XYZ positional stage (ThorLabs #MTS25-Z8) and is controlled by stepper motors (ThorLabs #KST101). The z-height is determined by bringing the stepper motors down until contacting the chip, and then programming the z-height via MATLAB. The line-slit is swept in 30um steps across the entire imager in both x- and y-directions, and in 100-μm increments in the z-direction. This calibration is only performed once, at the beginning and can be used for subsequent imaging experiments. The double line slit is fabricated using the same method to etch two 30μm line-slits, separated by 45μm into a Cr mask. For fluorescence scenes, we use 45-μm diameter fluorescence beads (Polysciences Fluoresbrite YG Microspheres #18242-2). The images were reconstructed using a Tikhonov regularized least-squares problem.

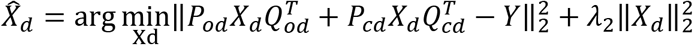

### Section S4. *In-vivo* experimental discussion

#### Optical crosstalk with ChRmine

The ChRmine activation spectrum is very-wide band allowing for optogenetic stimulation with a variety of wavelengths (**Fig. S24**). Given that 80% of the blue LED light directly contributes to induced photocurrent, the light power density used for gCaMP6f imaging needs to be maintained below the threshold for robust neural activation (< 0.1 mW/mm^2^)(*46*). The light propagation losses (**Section S2**) mean that the blue excitation light falls below (< 0.09 mW/mm^2^) the stimulation threshold for neurons located in the tissue. For one photon bidirectional optical experiments, the optical crosstalk can be further reduced by shifting to further, red-shifted opsins such as Chrimson(*77*).

#### Mesoscopic versus single cell

Cellular imaging using two-photon microscopy provides robust recordings of single neuron resolution activity with FoVs restricted to a few hundreds of microns. Alternatively, mesoscopic imaging is a rapidly growing widefield single-photon fluorescence imaging approach that provides extremely large multiple millimeter FoVs at high spatiotemporal resolution. Thus, mesoscopic imaging provides a powerful tool to investigate largescale neuronal networks spanning multiple brain regions at high resolution. One limitation of mesoscopic imaging is that the measured fluorescence signal reported from the superficial layer of the cortex is a weighted integration of neuronal signals by depth(*78*), and that the summed signal include contributions from cell somata and neuropil(*79, 80*). Another limitation is the overlap of green fluorescence with the endogenous activity-dependent absorption caused by blood volume and oxygenation. Typically, a separate green LED light source is used in reflectance mode to measure these hemodynamic signals to subtract them from the fluorescence video capture. However, the contribution of blood volume in GCaMP mice is relatively low compared to the larger activity-dependent fluorescence signals(*81, 82*).

### Section S5. Image sensor properties and filtering requirements

Fill factor (*FF*) is determined by (*A_SPAD_* is the cross-sectional active area of the SPAD, and *A_pixel_* is the cross-sectional area of the pixel):

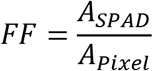

Photon detection efficiency (*PDE*) is given by (*PDP* is the photon detection probability and *FF* is the fill factor):

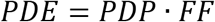

Noise equivalent power (NEP) is a measure of the sensitivity of a photodetector and is given by (*h* is Planck’s constant, *v* is the frequency of the photon, *DCR* is dark count rate, *PDE* is the photon detection efficiency):

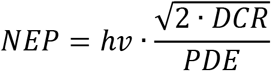

Noise · Power Figure of Merit (*Noise* is the square root of the dark count rate, *Power* is the measured electrical power of the sensor, and *R_P_* is the pixel rate):

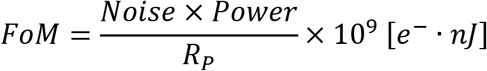

Pixel Rate (*N_H_* is the number of columns, *N_V_* is the number of rows, *r_VO_* is the readout time overhead per row, *r_HO_* is the readout time overhead per column):

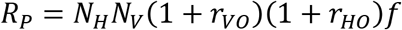

Dynamic range as determined by the number of bits (Q) of the counter is given by:

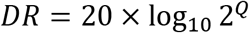

SNR equation (*P_sig_* is the signal power in photoelectrons, and *B* is the background power in photoelectrons):

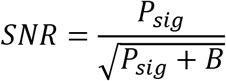

For simplicity, the received photons by the SPAD, *I*, assuming direct coupling can be determined by the number of photons per second per area leaving the LED (light power density normalized by the energy of a photon at 520nm), the cross-sectional area of the SPAD pixel, the photon detection efficiency, and *t_int_* the integration time per frame.

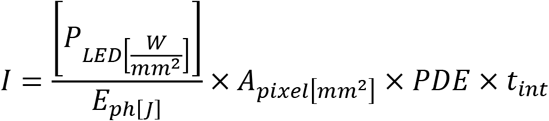

Signal power is determined coupling modeled by the blue to green fluorescence conversion efficiency (**Section S2**) and *I:*

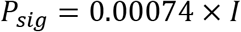

Background power is determined by the backscattering of the excitation light towards the sensor (Methods), *I*, and the optical density, *OD*, provided by the spectral filter stack:

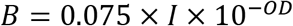

The signal and background powers are limited to the dynamic range of the sensor, which for a single frame, the number of sensor received counts much fall between 0 and 1023. This constraint allows us to solve for *I* as a function of received sensor counts *N_c_*:

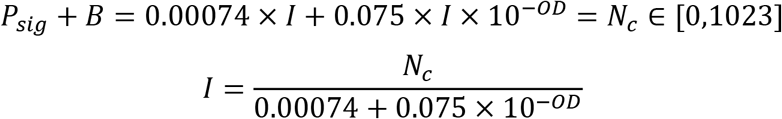

Finally, this result gets plugged into SNR equation and can be plotted for various values of *N_C_* and *OD* (**Fig. S12**):

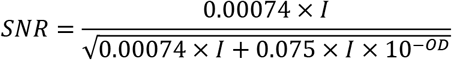

### Section S6. Nonhuman primate experiments

#### Feature decoder

Motor brain machine interfaces (BMI) consist of three main parts: a sensor to provide robust recordings of neural signals, a feature decoder to process the neural signal and predict a response or intention, and a prosthetic to restore lost functionality(*62*). Adaptive algorithms to track changes in recorded signals and predicted intent can also play a significant role. Here, we demonstrate using recordings during awake behavior the ability to deliver accurate recordings of optical neural signals in the motor cortex. To further validate the application of the SCOPe device to BMI, we developed a feature decoder that predicts movement intent from processed neural signals obtained by the SCOPe sensor. The decoder trained a ridge regression function that took as input predictor data a single frame of neural activity imaged by the device and returned as output the response data, a movement speed prediction. *y* denotes the response data specified as a n-by-1 vector, *X* denotes the predictor data specified as an n-by-p matrix, *k* is the ridge parameter that regularizes the regression function, and *I* is the identity matrix. The resulting coefficient estimator matrix, 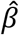, is of dimension (p+1)-by-1.

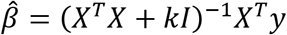

The predicted response data *y_p_*(*t*) is a n-by-1 vector determined by the summation of where *β*_0_ is the first coefficient estimate, *β_j_* are the remaining coefficient estimates (2 to p+1), and *X*_*t*′_ is the testing data specified as an m-by-p matrix at time *t*’.

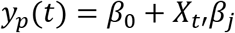

The premotor dorsal and motor cortex have been studied for encoding of kinematic parameters such as direction(*83, 84*), position(*85*), speed, and velocity(*86*). While direction is most strongly encoded in single unit activity, speed is most strongly encoded in high gamma local field potentials(*87*). The mesoscopic calcium fluorescence imaging modality (**Section S4**) employed in our system is able to resolve multiunit activity, but not single unit spiking events. We, therefore, focused on decoding movement speed trajectories. Since the motor cortices predict movements that occur in the future, we varied the relative time of t and t’. To quantify prediction performance, we measured the correlation coefficients for the predicted speed from the feature decoder and the ground truth speed measured via behavior.

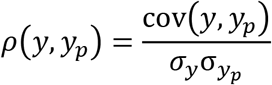

**Table S1.** Comparison table with state-of-the-art miniscopes. Table comparing the type (1P, Lensless, 2P), weight, size (mm^3^), FoV(mm^3^), displacement factor, resolution, power, and bidirectional (recording and stimulation) capabilities of various miniaturized optical microscopes. ^†^Estimated from figures and scale bars. ^‡^Estimated from image sensor data sheet. Other power-hungry blocks such as the LED power consumption are rarely reported. The total system power consumption is underestimated and is likely to be much greater. ^§^Measured total power consumption including image sensor and LED power.

|  | Ref | Ref | Ref | Ref | Ref | Ref | Ref | Ref | This Work |
| --- | --- | --- | --- | --- | --- | --- | --- | --- | --- |
| Name | Original | Wireless | MiniScope | CM<sup>2</sup> | Mini-3D | mScope | 2P MiniScope | FlatScope | SCOPE |
| Type | 1P | 1P | 1P | Lensless | Lensless | 1P | 2P | Lensless | Lensless |
| Weight (G) | 1.9 | 1.8 | 4 | 19 | 2.5 | 4 | 4.2 | 3 | 0.107 |
| Size (mm<sup>3</sup>)<sup>†</sup> | 10×10×15 | 20×10×10 | 25×15×15 | 29×29×30 | 10×10×17 | 25×15×15 | 16×9×30 | 8×8×8 | 6.4×7.8×0.15 |
| FoV (mm<sup>3</sup>)<sup>†</sup> | 0.6×0.8×0.2 | 0.6×0.8×0.2 | 0.8×0.8×0.4 | 8×7×2.5 | 0.9×0.7×0.39 | 8×10×0.2 | 0.42×0.42×0.18 | 4×4×0.5 | 5.1×6.8×0.5 |
| Displacement | 16,000 | 28,000 | 22,000 | 180 | 7,000 | 350 | 140,000 | 64 | 0.5 |
| Res (μm) | 2.5 | 4 | 6 | 7 | 2.76 | 39.37 | 1.13 | 10 | 60 |
| Power (mW)<sup>‡</sup> | 160 | 350 | 363 | 381 | 900 | 226 | N/R | 2,500 | 10<sup>§</sup> |
| Bidirectional | No | No | No | No | No | No | No | No | Yes |

**Fig. S1.**
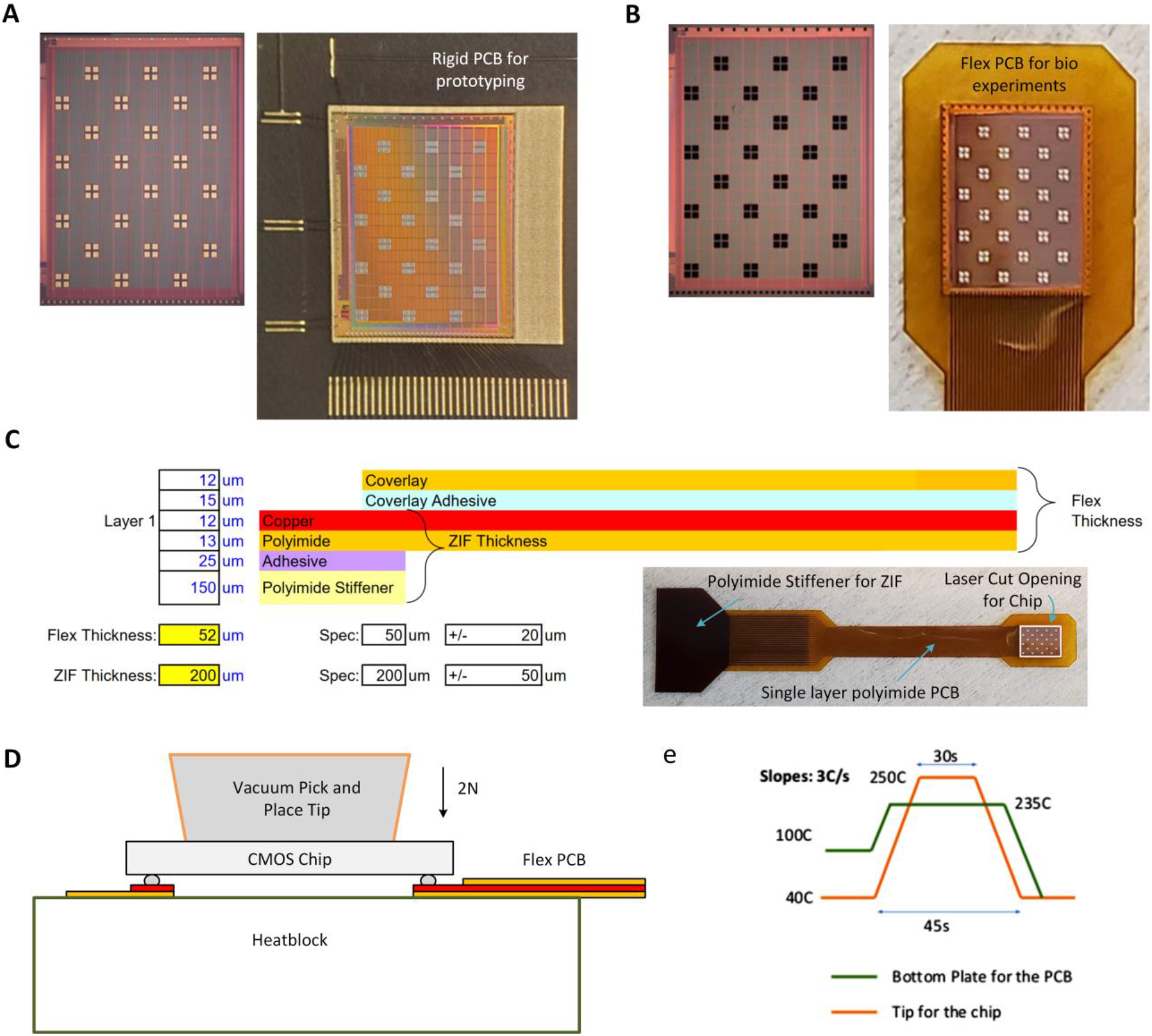
Flip-chip bonding to flexible substrates. (A) Wirebond variant of back-end-of-line (BEOL) CMOS fabrication flow used for electrical and optical prototyping and characterization. The pads are wirebonded to an Electroless Nickel Electroless Palladium Immersion Gold (ENEPIG) surface-finish rigid PCB. (B) Flip chip variant with SnPb solder alloy bumps with 70μm bump diameter and 170μm pitch. (C) Flex PCB is designed for minimum thickness from the specified board house vendor for a single layer polyimide flex PCB. The total flex thickness is 52μm and a polyimide stiffener is added to the connector end of the flex cable for a total ZIF thickness of 200μm. There is a laser cut opening in the flex PCB to leave an optical window for the sensor to directly contact the tissue. (D) Cross section view of the flip chip bonding process using Finetech FINEPLACER lambda for precision die attach to the flex PCB vacuum mounted onto the bottom plate heat block. A force of 2N is applied during the application of a heating program to allow for bump reflow without shorting between adjacent pads. (E) The heating program showing the temperature profiles for the bottom plate (green line) and the pick and place tip (orange trace).

**Fig. S2.**
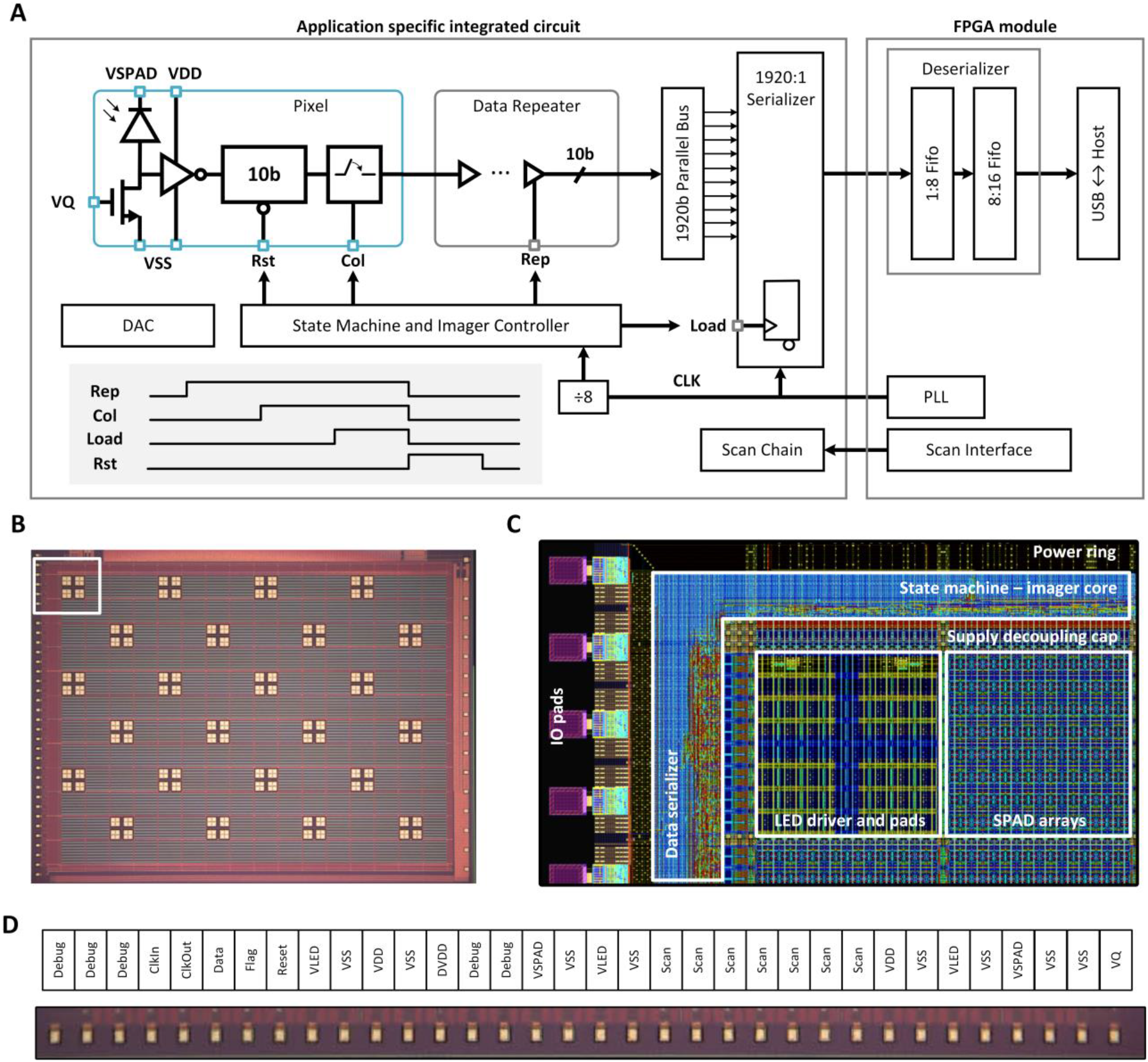
Detailed overview of the CMOS system architecture. (A) The system on chip architecture consists of an analog 192 × 256 SPAD pixel array which interfaces with a digital imager core, data transmission block, and scan chain IO configuration. The pixel has a dual-SPAD architecture where the outputs of two SPADs are summed into a shared 10b in pixel counter. The chip’s controller determines the state machine of the imager core with the readout operation performed in four steps: 1) the thermometer encoded REP signal is asserted for three clock cycles to allow the repeaters throughout the array to buffer the data, 2) the one-hot read column signal is enabled for two clock cycles to pass the data through the in pixel output transmission gates onto the repeater bus, 3) the load signal is used to latch the data into the data transmission serializer flip flops, and 4) REP, COL, and LOAD get de-asserted and RST gets asserted for one clock cycle to clear the data in the pixel counter. The digital clock runs on a clock division by eight of the reference clocks to save power. The data is streamed off-chip at 100MHz using the signals CLK, FLAG, and DATA. (B) Die photo of the fabricated chip. (C) Computer aided design layout (of white rectangle from B) highlighting the analog imager SPAD arrays, the digital core controller containing the imager core and data serializer, the power ring, and the IO pads. (D) A detailed view of the chip’s IO and power signals resulting in a total of 34 pads. Many of the supplies are redundant to reduce the IR droop and minimize inductance and noise.

**Fig. S3.**
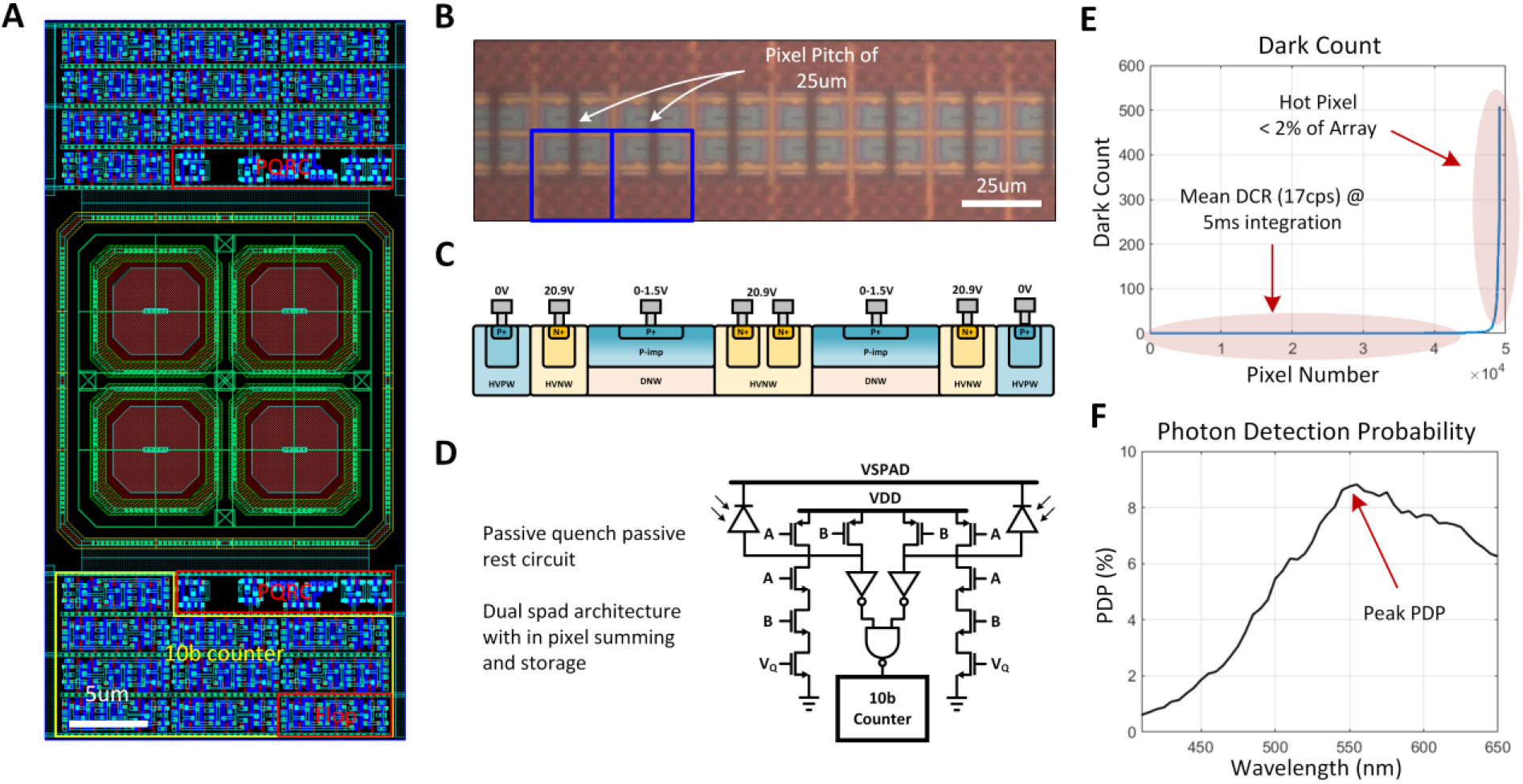
SPAD characterization. (A) The layout of a 2 × 1 SPAD pixel with the top two SPADs summing into the top pixel 10b counter and the bottom two SPADs summing into the bottom pixel 10b counter. Since the standard cell flip flops consume more area and contain buffers for timing and driving large capacitive loads, a custom in-pixel flip flop was designed to minimize area usage so that the 10b asynchronous ripple carry counter could be brought into the pixel. (B) Micrograph of a row of pixels and a pixel pitch of 25μm. (C) Cross section of the SPAD well structure (as seen from the red line in a). The SPAD structure is a P implant diffusion layer sitting in a deep N well optimized for longer wavelength sensitivities. The active diameter of each SPAD is 6μm and N well sharing between the two adjacent chamfered square SPADs, results in a total active area of approximately 72 μm^2^ and a fill-factor (FF) of approximately 11.5%. Each quad of SPADs sits in an isolation ring to prevent substrate noise from increasing the dark count rate. The quenching circuit is passive-quench and passive-reset (PQRC) with dead-time determined by the VQ bias on the quench transistor. An on chip, array shared 3b digital to analog converter (DAC) provides the VQ bias from 450mV to 1.07V in steps of 80mV. (E) The SPAD dark count rate is measured by evaluating the detected counts over a 1s integration in an optical dark room where no visible photons are present. The median dark count rate is calculated to be around 17Hz. Hot pixels are defined as pixels that have average dark count rates above a standard deviation of the median and are typically caused by fabrication defects in the SPAD structure. The total hot pixel account for less than 2% of the array. (F) The SPAD peak photon detection probability (PDP) is 9% at 550nm and an excess bias voltage of 1.5V. The figures are adapted from Ref(*69*).

**Table S2.** SPAD characterization. Table comparing imaging modality (time-resolved or intensity), array size, technology, pixel pitch, PDP, DCR, frame rate, dynamic range, and power consumption with current state-of-the-art SPAD imagers.

|  | **Ref (27)** | **Ref (28)** | **Ref (30)** | **Ref (29)** | **Ref (19)** | **Ref (31)** | **This Work** |
| --- | --- | --- | --- | --- | --- | --- | --- |
| **Name** | Time-Resolved | Time-Resolved | Time-Resolved | Time-Resolved | Time-Resolved | Time-Resolved | Intensity |
| **Array Size** | 160×120 | 256×256 | 512×512 | 256×256 | 160×160 | 128×120 | 192×256 |
| **Technology** | 350nm | 130nm | 180nm | 90/40nm | 130nm | 90/40nm | 130nm |
| **Pitch (Fill Factor)** | 15um, 21% | 16um, 61% | 16um, 10.5% | 9.2um, 51% | 30um, 5% | 8um, 45% | 25um, 11.5% |
| **PDP (Excess Bias)** | -- | 39.5%, 3V | 50%, 6.5V | 23%, 3V | 12%, 3V | 12.6%, 3V | 9%, 1.5V |
| **DCR (Excess Bias)** | 580cps, 3V | 40cps, 1.5V | 75cps, 6.5V | 20cps, 1.5V | 26cps, 3V | 15cps, 1V | 17cps, 1.5V |
| **Frame Rate (fps)** | 486 (8b) | 100,000 (1b) | 274 (8b) | 30 (8b) | 125 (10b) | 15 | 200/400 |
| **Sensitivity** | 1.06 fW/pixel | 0.06 fW/pixel | 2.00 fW/pixel | 0.07 fW/pixel | 1.65 fW/pixel | 0.10 fW/pixel | 0.63 fW/pixel |
| **Dynamic Range** | 74 dB | 64.4 dB | 48 dB | 120 dB | 60.2 dB | 72dB / 126 dB | 94 dB |
| **Power** | 157mW | -- | 26.7 mW | 77.6 mW | 40 mW | 10 mW | 20 mW |

**Fig. S4.**
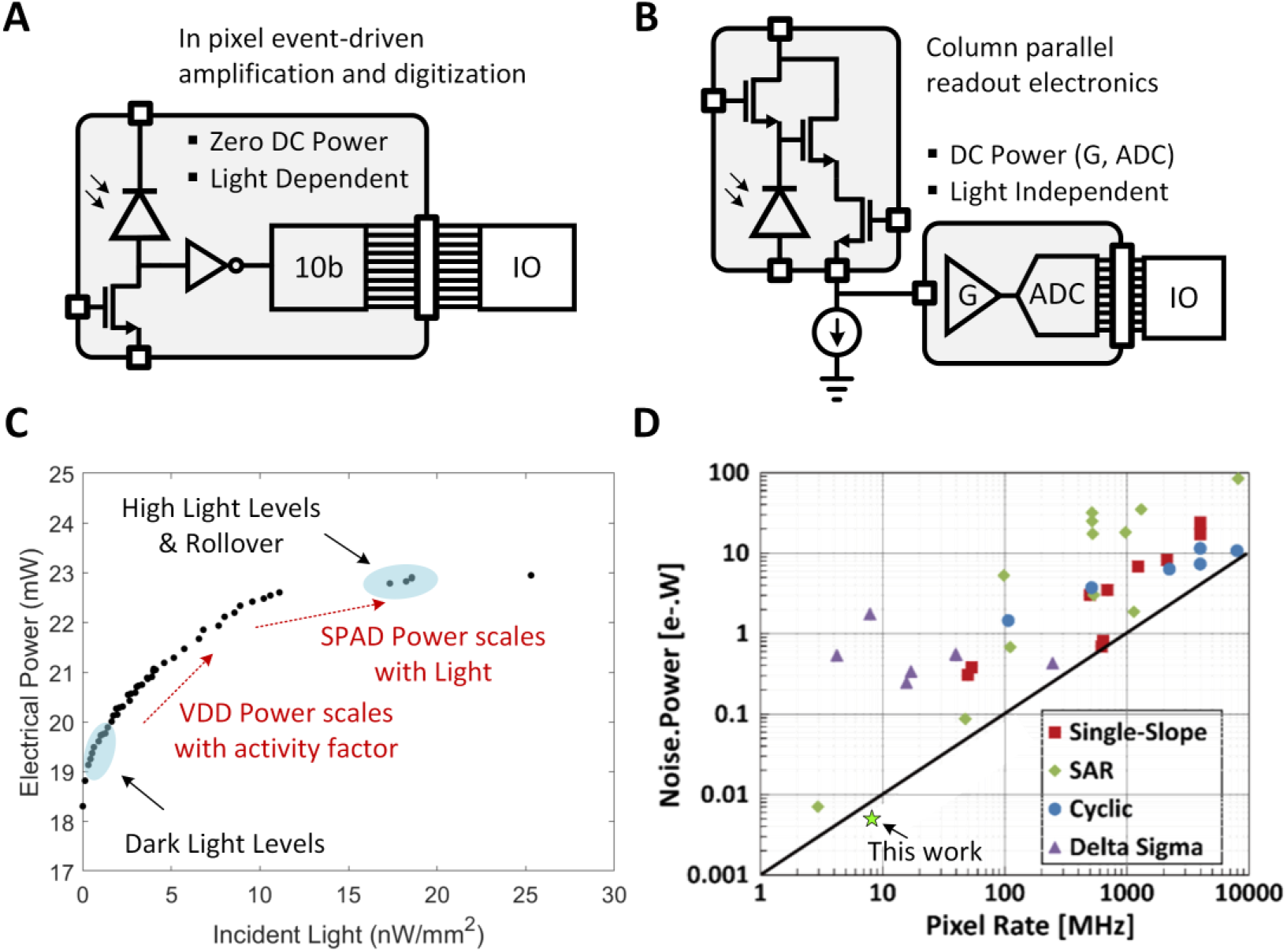
SPAD power consumption. (A) Since the SPAD is an activity-based high-gain sensor, power consumption scales with light intensity, and is a function of the energy consumption of a single avalanche event(*88*) *E_ava_* = *C_SPAD_* × *V_DD_* × *[V_DD_* + *V_BD_]* given an excess bias voltage of *V_DD_* and multiplied by the incident photon flux. This contrasts to (B) photodiode imagers which rely on fixed amplification overhead in the pixel shared column readout amplifiers and analog to digital conversion. (C) Our SPAD image sensor consumes 18.5mW in the dark and 22mW at peak saturation light intensity. (D) CIS- and SPAD-based analog-to-digital signal acquisition is compared at light level of 2 nW/mm^2^ by plotting the product of noise, as defined by 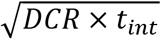, and power on the y-axis and pixel rate (the product of the number of rows, number of columns, and sensor readout rate) on the x-axis. A figure-of-merit (FoM) (*32*) is defined at the ratio of the noise-power product to the pixel rate. Out SPAD sensor (pixel rate = 9.83MHz) shows comparable or better FoM performance than low-noise CMOS image sensors.

**Fig. S5.**
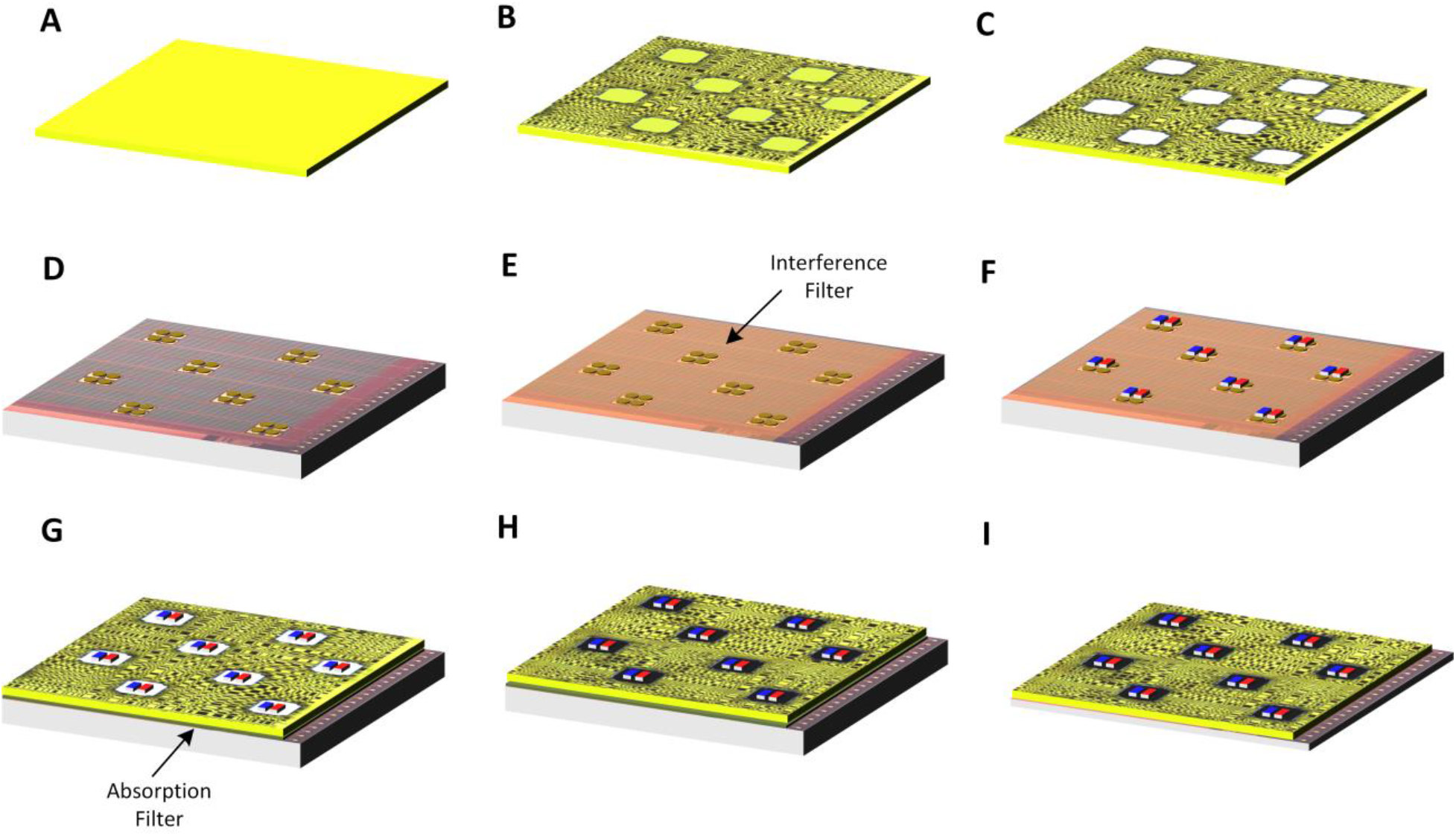
Light sources, filters, and computational mask packaging. Flexible Packaging stack up with less than 250um thickness as well as a functioning die-thinned chip bended along a radius of curvature of 10mm. (A) Polyimide substrate layer (B) Computational mask with Cr patterning, (C) Excimer laser cut-outs, (D) CMOS chip with C4 bumps for μLED contacts, (E) Thin-film dielectric interference filter coating, (F) Flip-chip bonding of μLEDs, (G) Bonding of computational mask to the interference coated and μLED bonded chip using custom absorption filter epoxy, (H) Black underfill epoxy of μLEDs, and (I) CMOS substrate die-thinning. Note: The packaging flow is done in two parallel steps, the mask fabrication in steps (A)-(C) and the interference filter and μLED bonding in steps (D)-(F). The two parallel flows are combined together in step (G) via an absorption filter adhesive bonding step.

**Fig. S6.**
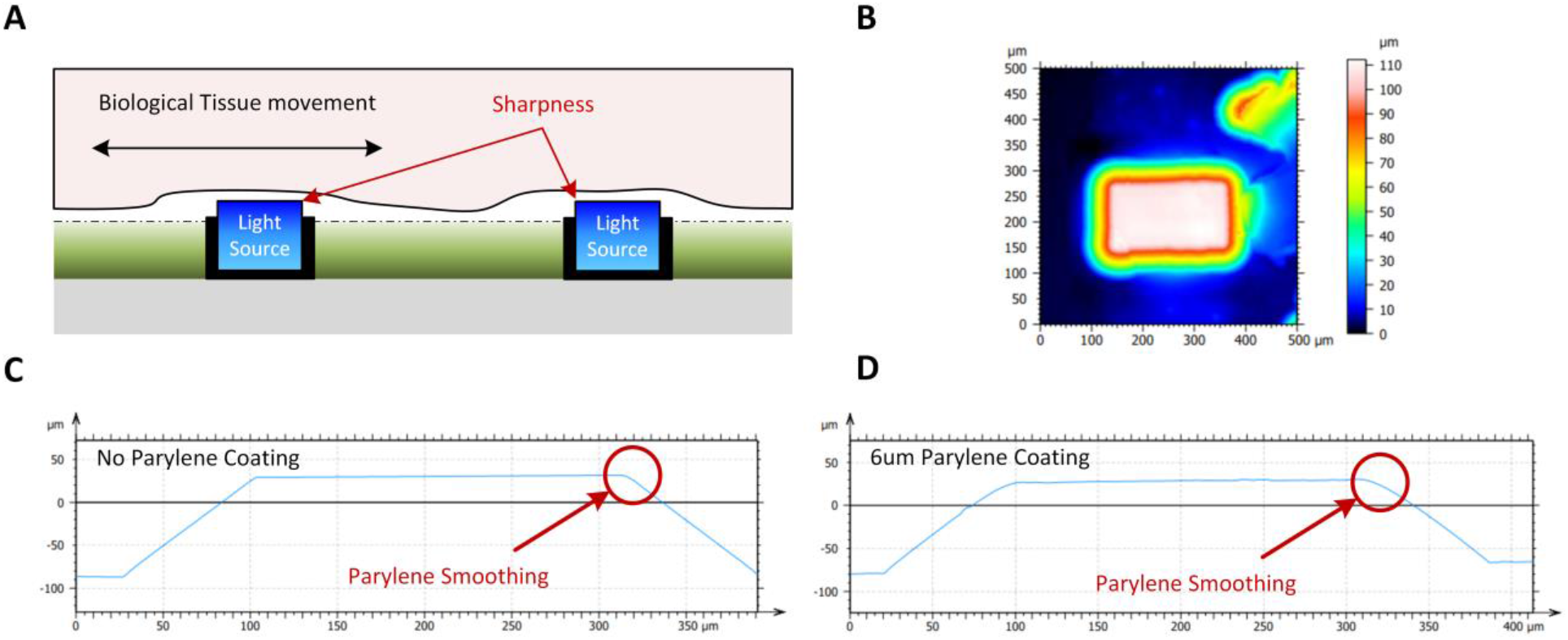
LED sharpness. (A) Cross-section of the CMOS sensor with integrated light sources showing the potential for shearing due to biological tissue movement, (B) Topographical image captured using profilometer, (C) Comparison of smoothness of the μLED before and after 6μm of parylene coating.

**Table S3.** Mechanical properties of materials.

| **Materials** | **Young's Modulus** | **Ref** |
| --- | --- | --- |
| Parylene C | 2.8 GPa | Ref (89) |
| Polyimide | 2.07-2.76 GPa | Ref (90) |
| CMOS (0.5um three metal n-well CMOS process) | ~60GPa | Ref (91) |
| Absorption filter (~20um KMPR photoresist) | 6.5–7.5 GPa | Ref (92) |
| Interference filter | e.g., 0.9085GPa | Ref (93) |

**Fig. S7.**
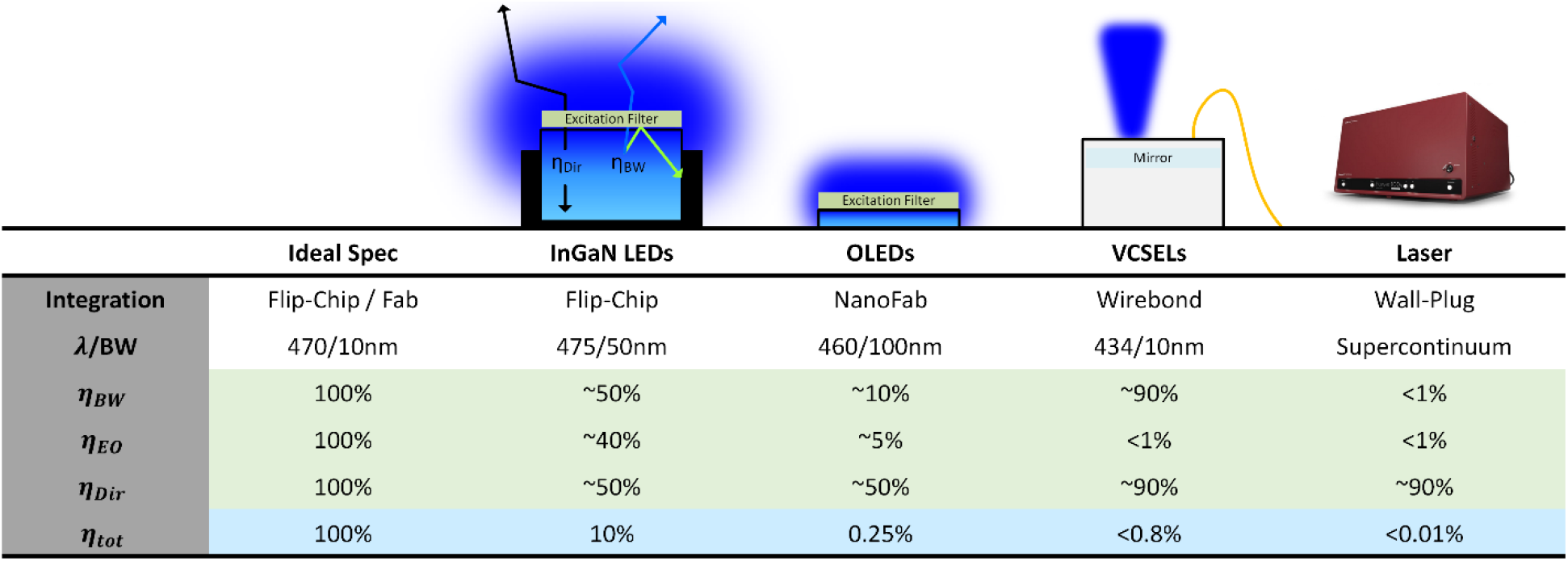
Comparison of integrated light sources. The efficiency of the light source is given by the cascade of spectral (*η_BW_*), electro-optic (*η_EO_*), and directional (*η_Dir_*) efficiencies. Flip-chip InGaN LEDs and OLEDs have relatively wide linewidths necessitating the use of excitation filters. Flip-chip InGaN μLEDs have the best electro-optic efficiency compared with OLEDs and VCSELs. Combining these results, the flip-chip InGaN LEDs have relatively the best efficiency, additionally with the ease of packaging of not having to wirebond and encapsulate.

**Fig. S8.**
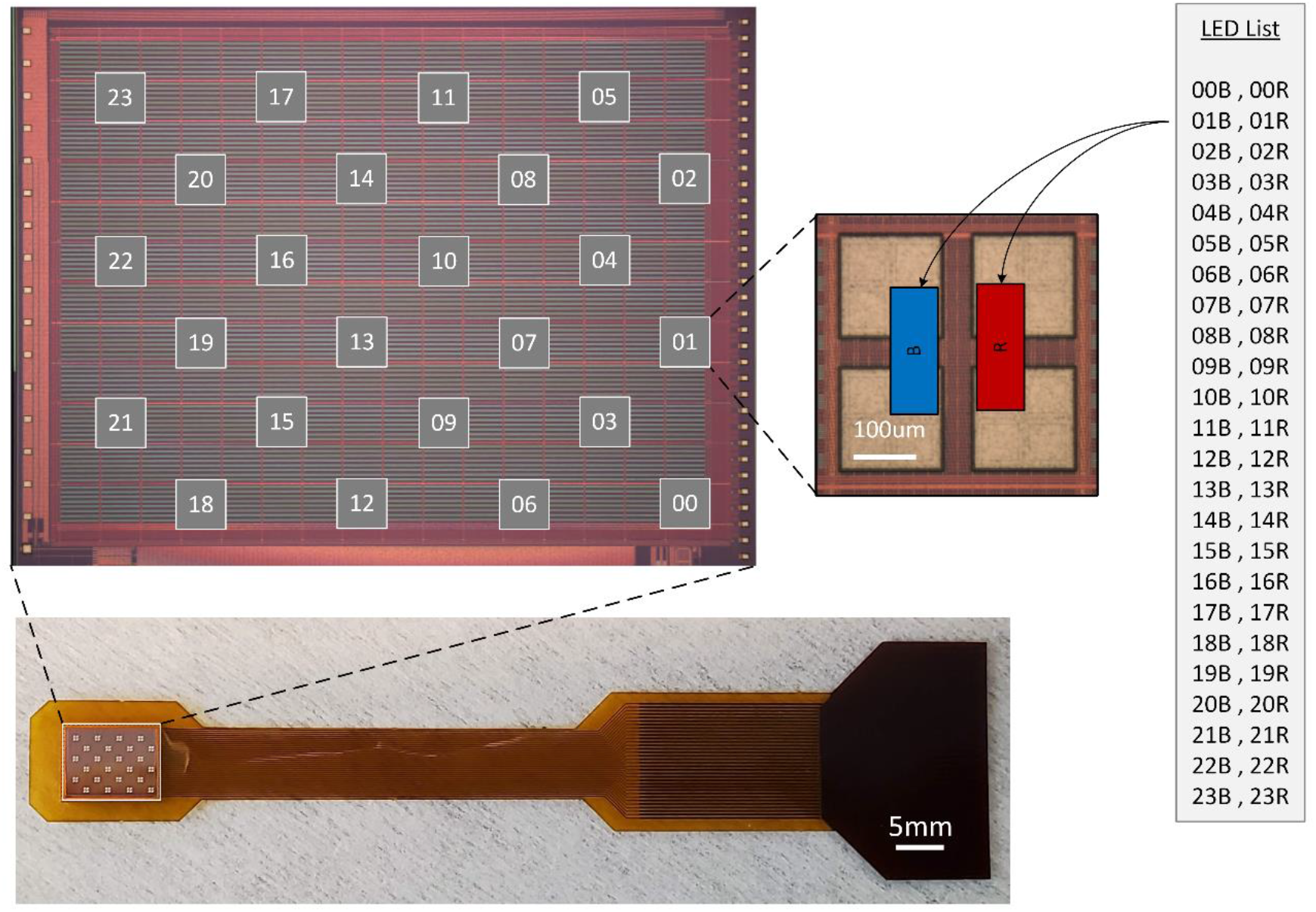
LED map for orientation and localization in animal experiments. The LEDs are number in pairs from 00 until 23 with each pair containing a blue and red LED. In the electrical stimulation mouse experiment, the flexible shank electrodes provide biphasic current stimulation at LED pair location 10, and the blue recording LEDs are located at 07B, 08B, 09B, 10B, 13B, and 14B. In the optogenetic stimulation and optical recording mouse experiment, the red stimulation LED is located at 08R, and the quad of blue recording LEDs are located at 10B, 13B, 14B, and 16B. In the NHP experiment, all the blue LED locations are enabled for full FoV imaging. The dead LED locations correspond to 02B and 22B.

**Fig. S9.**
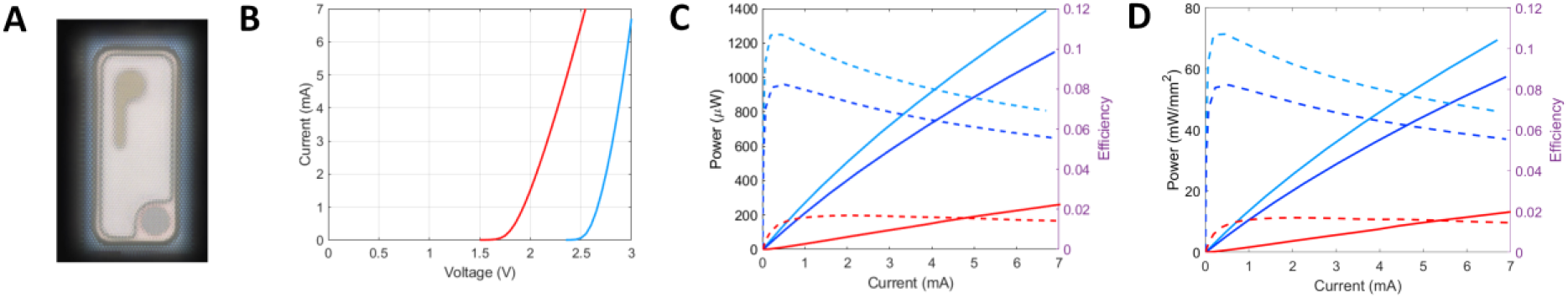
LED electrical and optical characterization of LEDs. (A) Die photo of the LA UB08FP2 blue LED chip with a surface emission area of 0.2mm × 0.1mm. Anode contact is on the top pad and cathode contact is on the bottom pad (B) Current versus voltage sweep for blue and red LEDs using the drivers on the chip. (C) Measured output light power captured using ThorLabs PM100D photodiode module. The efficiency is calculated by taking the light power and normalizing to the electrical power. (D) Output light power density taken by normalizing the output light power to the radiative surface area of the LED chip.

**Fig. S10.**
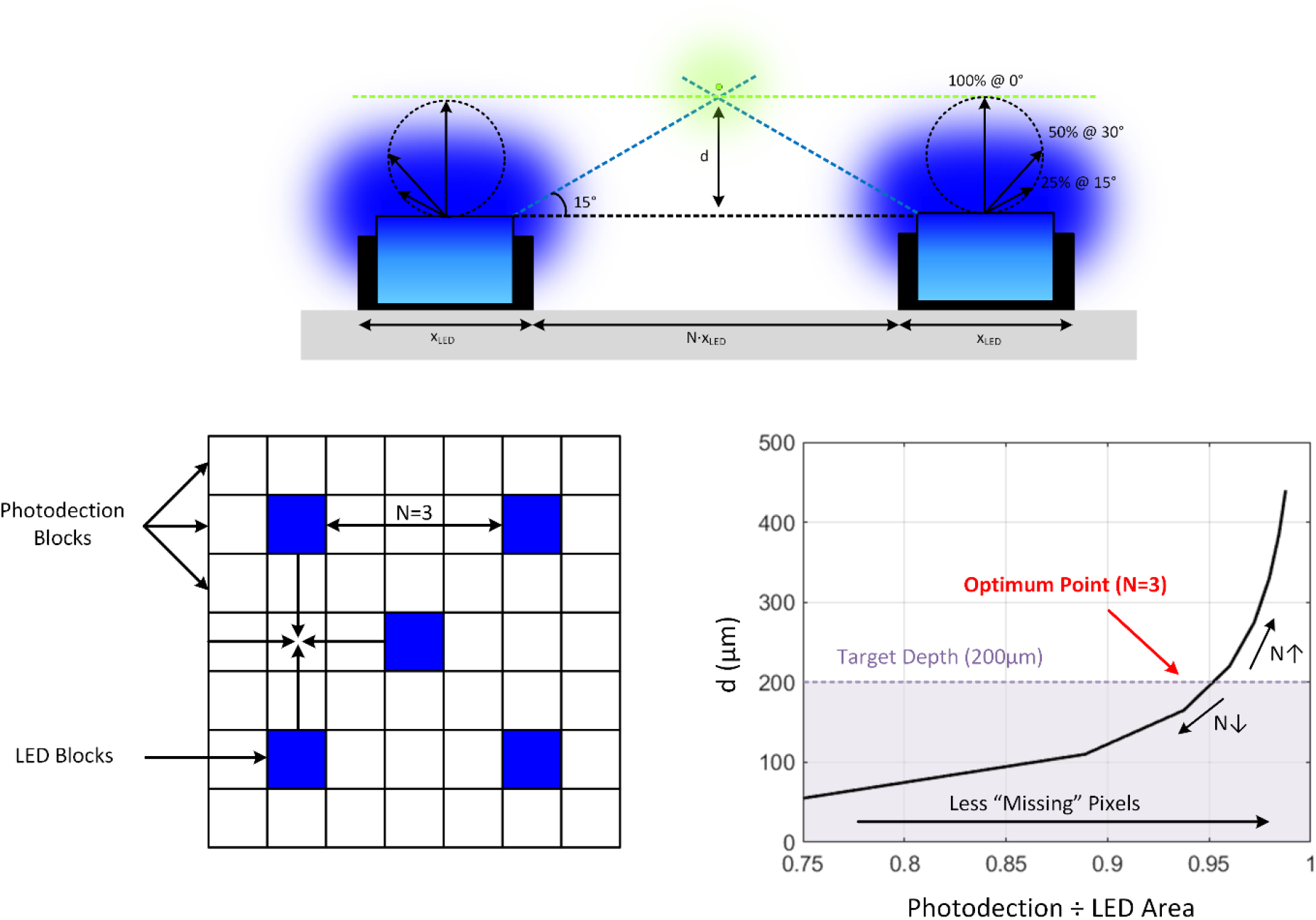
Pitch of light sources. The μLED emission profile is approximately Lambertian meaning that emission angle that corresponds with 25% power output is 75° from normal incidence. The pitch between μLEDs is optimized such that the target depth of 200μm is covered within the FWHM of adjacent μLEDs while allowing for the highest ratio of photodetectors to μLED blocks. In this design, each photodetection block consists of 16×16 arrays of SPADs. Sweeping the number (N) of photodetection blocks between μLEDs gives an optimum point of N=3 where there is FWHM overlap between adjacent μLEDs yet still allows for 87.5% of the array to be covered by photodetectors.

**Fig. S11.**
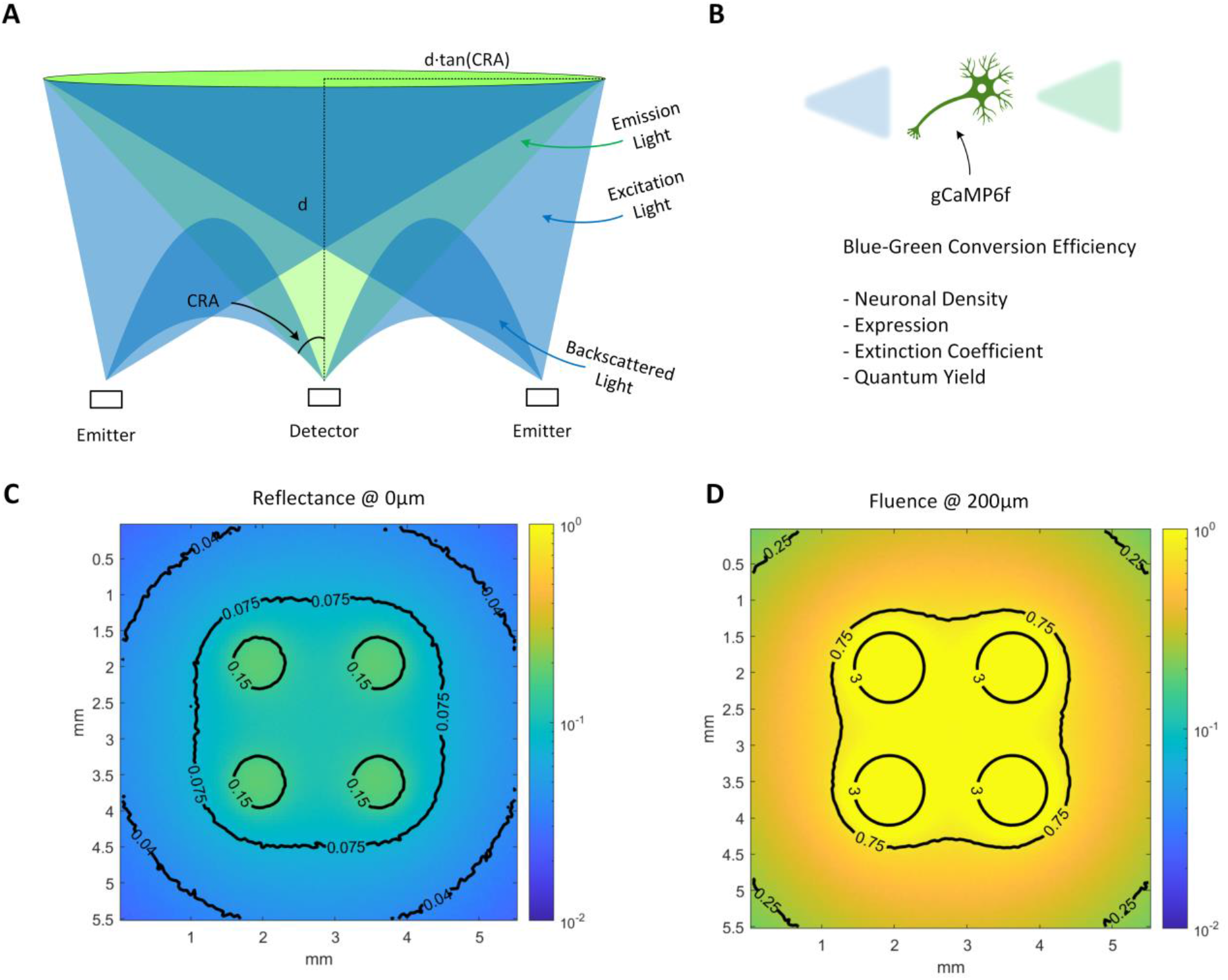
Analysis of biophotonics of SCOPe. (A) Modeling of μLED excitation and backscattered light with respect to the source detector separation and geometry as well as the fluorescence emission and angular acceptance of the SPADs. (B) The fluorescence signal for a given volume is proportional to the intrinsic neuronal density of the tissue, the percentage of gCaMP6f expressing neurons, and the extinction coefficient and quantum yield of gCaMP6f. (C) The reflectance is the amount of light exiting the tissue at z=0μm and is modeled using a Monte Carlo simulation using the μLED pitch and Lambertian source emission profile. (D) The fluence is the amount of light at the z=200μm plane normalized to the μLED light source power. Due to the effects of scattering, there is a focusing of power density to the superficial layers of the tissue resulting in lower required μLED light emission.

**Fig. S12.**
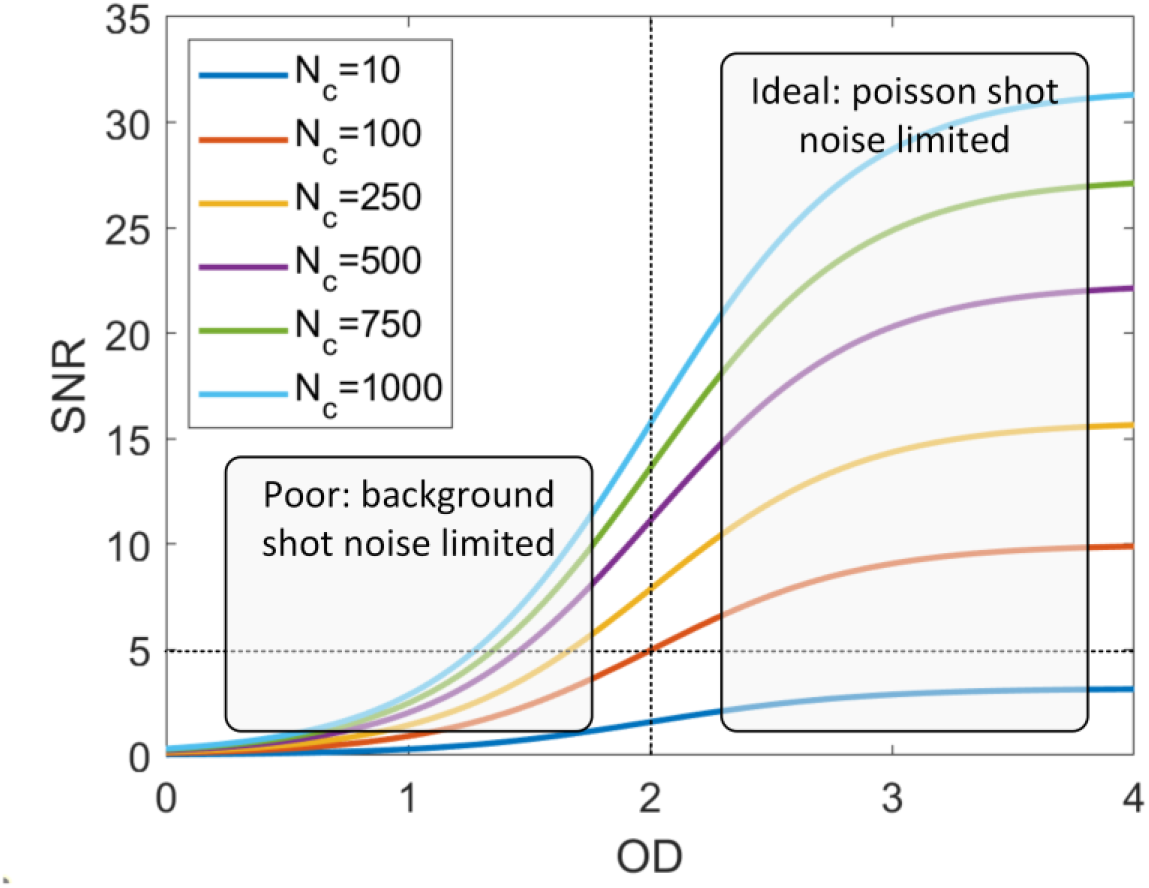
Filtering equations with SNR. The SNR curves corresponding with the equations in **Section S5** for various sensor counts per frame and filtering optical densities. As expected, the SNR curves increase with SPAD sensor counts and with OD filtering improvements. For OD < 2, the SNR is limited by background shot-noise, and for OD > 2, the SNR is limited by Poisson shot noise with limited improvements due to additional filtering.

**Fig. S13.**
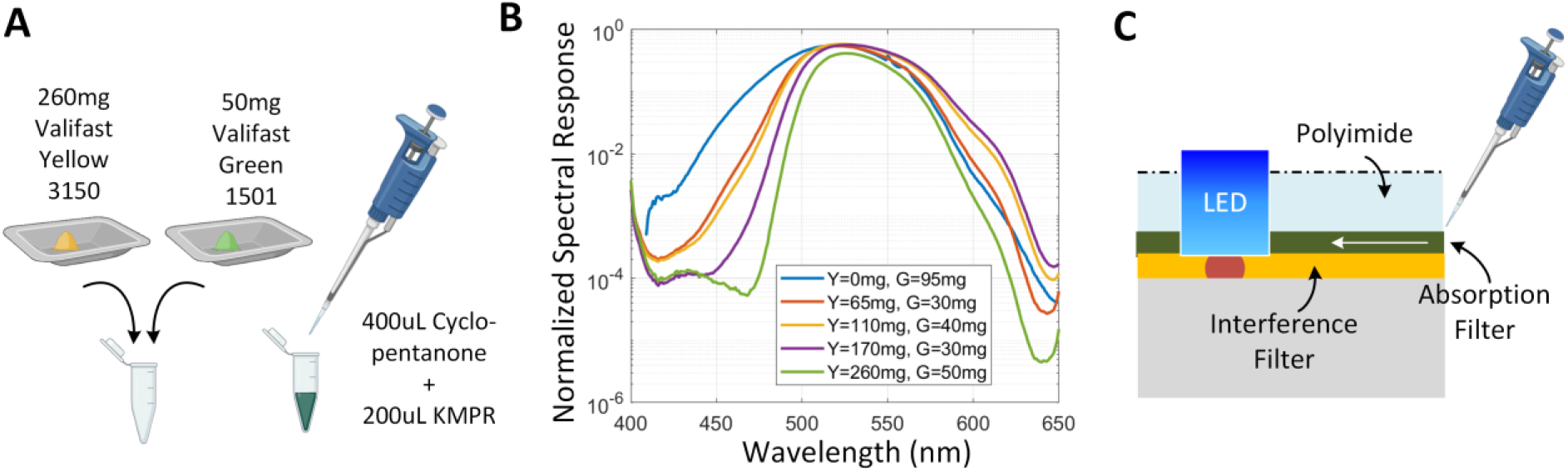
Optimization of absorption filter. (A) The absorption filter is synthesized using a combination of 260mg Valifast Yellow 3150, 50mg Valifast Green 1501, 400μL Cyclopentanone, and 200μL KMPR, (B) Spectral transmission plots for custom absorption filters using various quantities of Valifast Yellow and Green dyes, and (C) The polyimide spacer used for the computational mask also serves as a top substrate for injection of the absorption filter using capillary force for a final thickness of ~10-20μm.

**Fig. S14.**
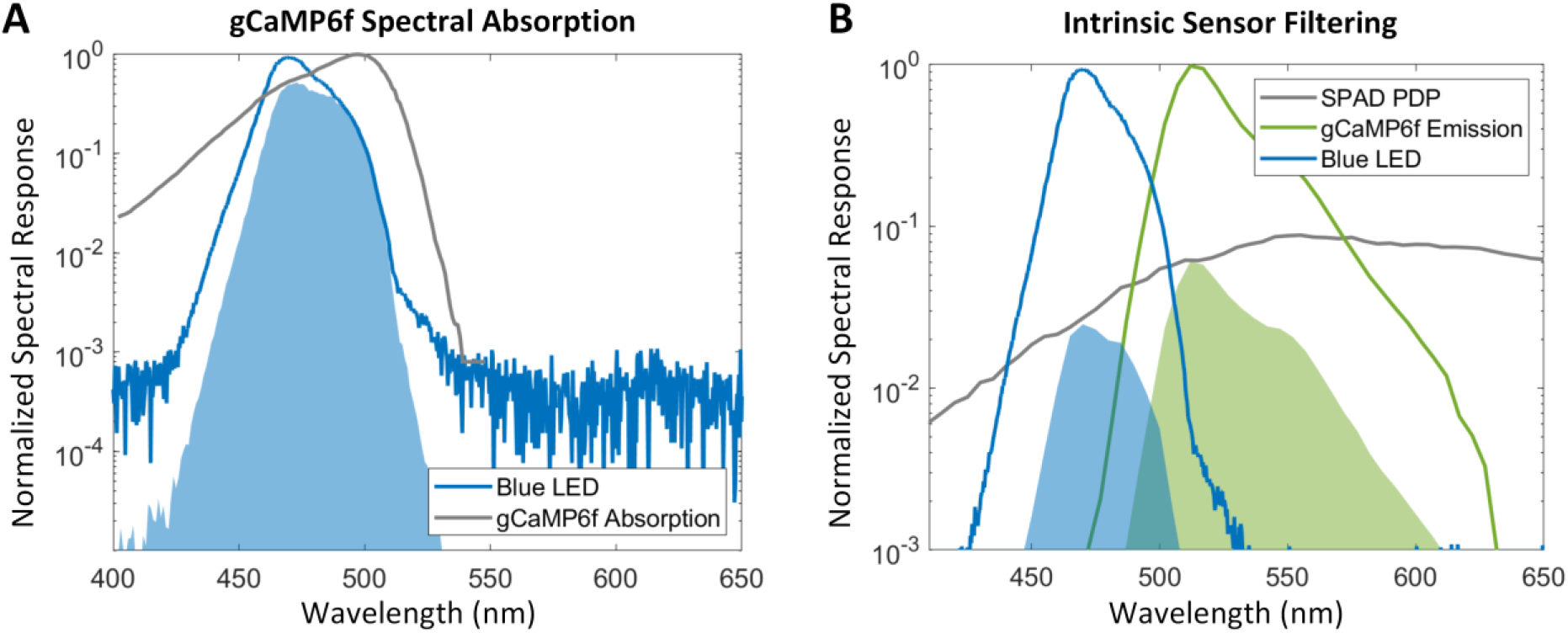
Spectral sensitivity. (A) The spectral overlap of the blue excitation LED with the excitation filter and the gCaMP6f absorption spectrum. Due to the peak mismatch and the spectral bandwidth of the light source, the resulting spectral overlap is 60%. (B) The sensor pixel has a photon detection probability that varies across wavelength with slightly greater sensitivity in the green and red wavelengths. Integrating the blue excitation LED spectrum gives an integrated photon detection probability of 3.1% and the gCaMP6f emission gives an integrated photon detection probability of 6.75%. Taking the ratio of these two integrated PDP values gives the sensor an intrinsic filtering of 2.2:1.

**Fig. S15.**
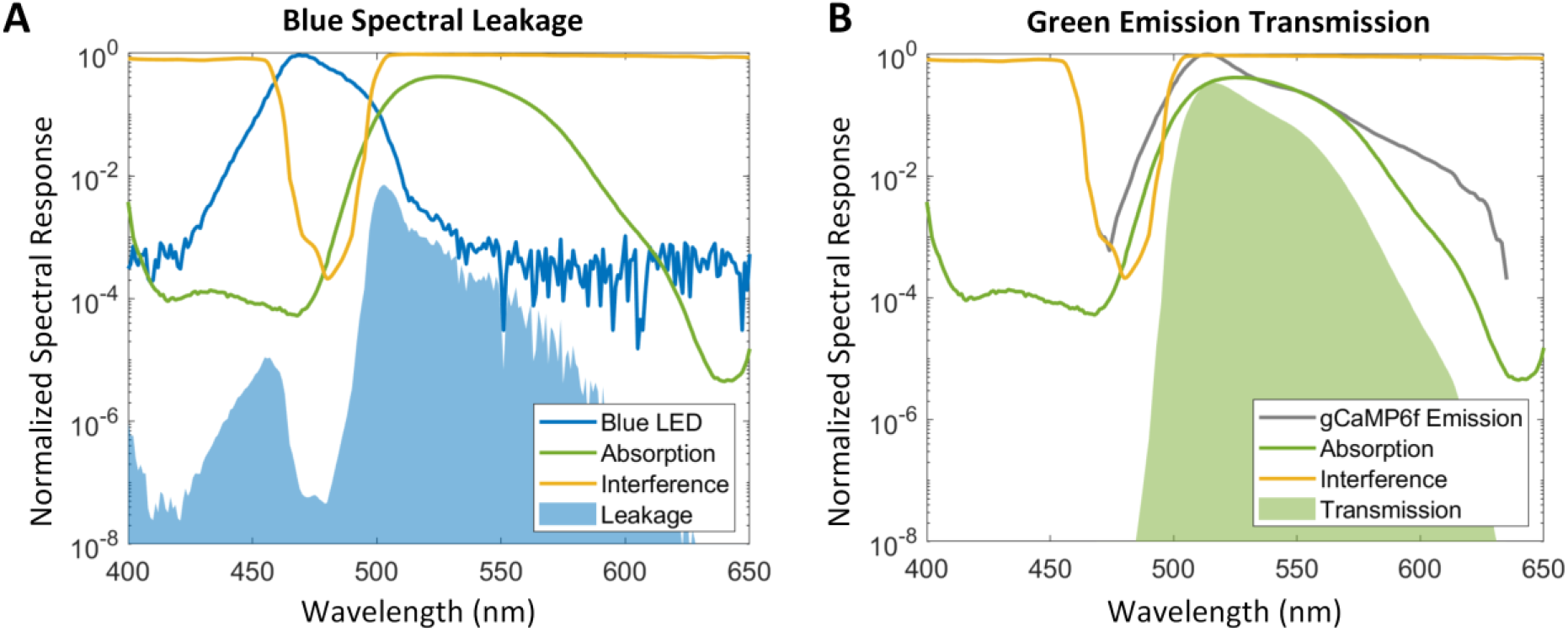
Filter characterization. (A) The total spectral leakage of the blue LED through the excitation and emission filter stack. The interference provides steep transition between the stopband and passband while the absorption filter provides wide stopband rejection across the blue LED spectrum. Leakage is calculated by multiplying the spectral responses together and integrating across the entire frequency domain resulting in a leakage of 0.0037 and contributing to the background in sensor captures. (B) The total spectral transmission of the gCaMP6f fluorescence through the emission filter stack is 0.2677. The total OD filtering is the ratio of these two values which results in OD > 1.9, or 72:1 rejection of light source photons in favor of emission photons.

**Fig. S16.**
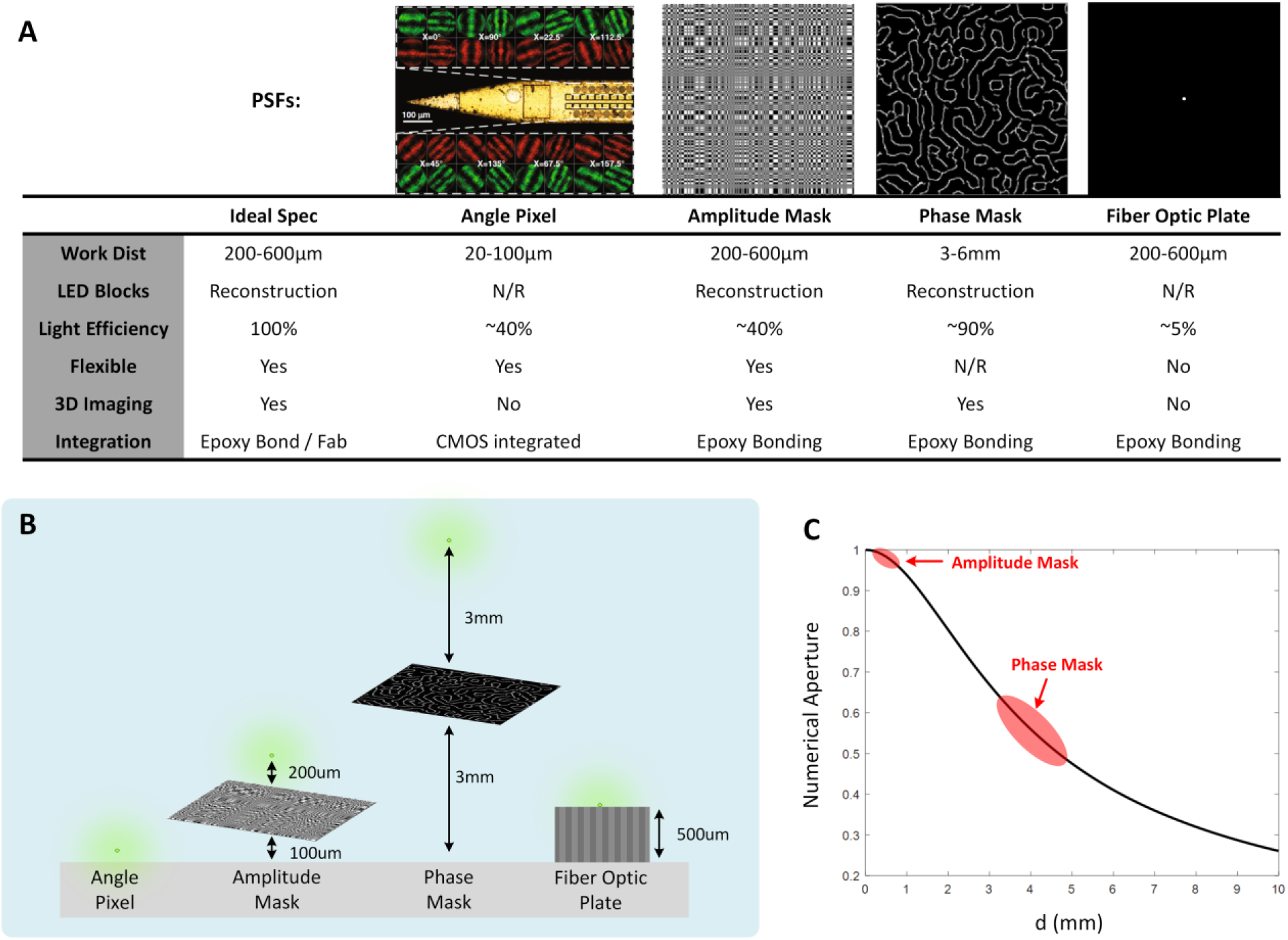
Comparison of computational mask designs. (A) We evaluated various types of computational imaging masks on the basis of working distance, reconstruction with LED blocks, light collection efficiency, flexibility, 3D capabilities, and ease of integration. (B) Comparison of working distances of the angle pixel(*49*), amplitude mask(*16, 19*), phase mask(*17*), and fiber optic plate. (C) Numerical aperture, comparing amplitude and phase masks.

**Fig. S17.**
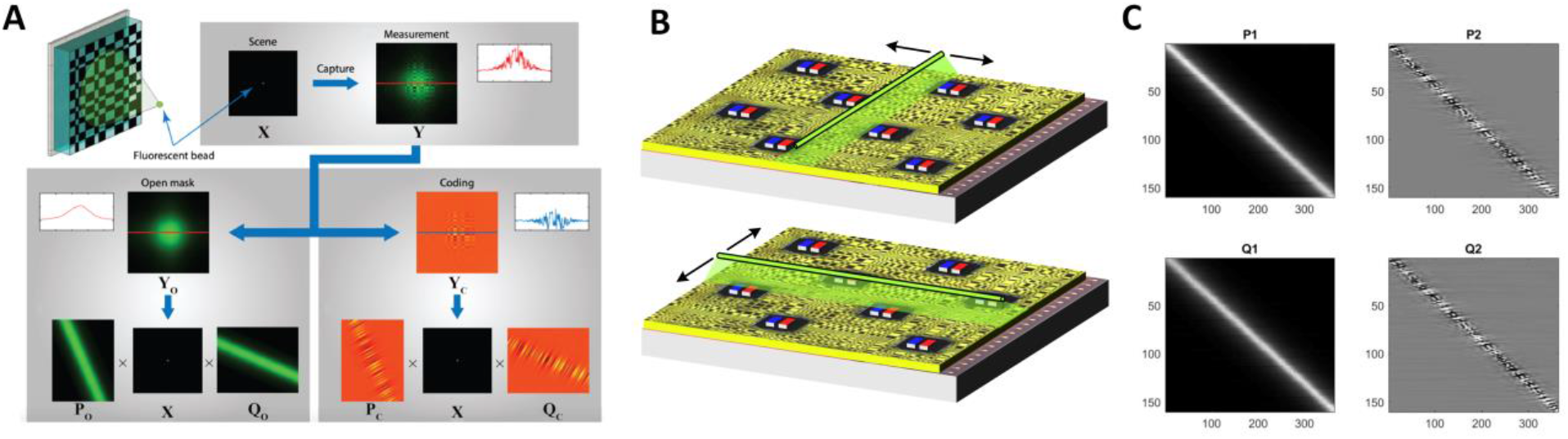
Computational imaging flow – calibration. (A) Texas Two-Step algorithm (adapted from Ref(*16*)) allows for reduction in calibration time and data storage based off of separable mask design, (B) calibration setup showing the sweeping of a line slit in row / col directions, (C) the P_1_, P_2_, Q_1_, and Q_2_ calibration matrices where the P_1_, P_2_ matrices encode the intensity information, and the Q_1_, Q_2_ matrices encode the mask modulation information.

**Fig. S18.**
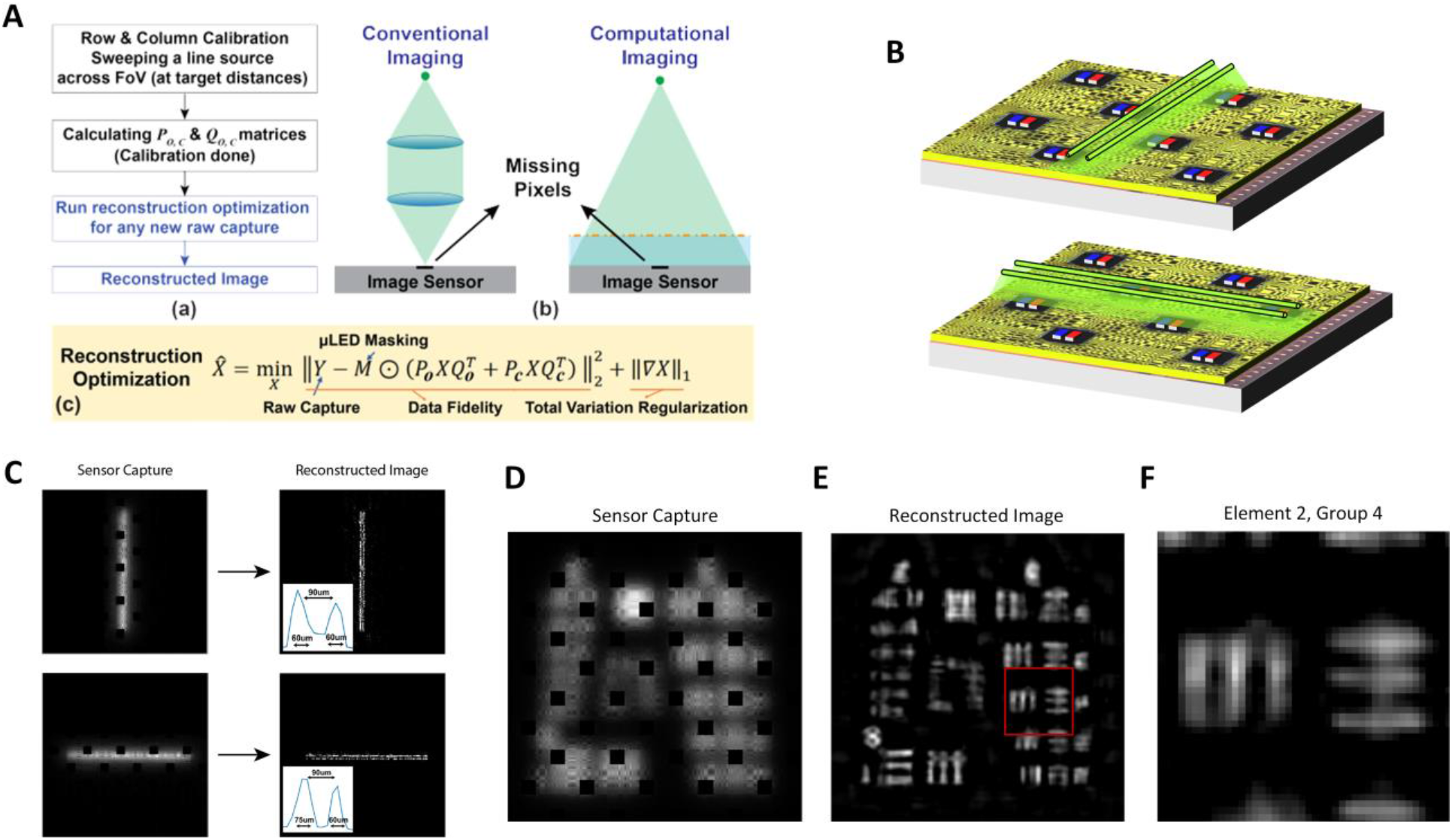
Computational imaging flow – reconstruction. (A) Overall computational imaging flow including calibration, calculation of calibration matrices, raw image capture and computational reconstruction using an inverse optimization algorithm (adapted from Ref(*19*)), (B) Double line slit reconstruction setup for demonstrating resolution in the row and column dimensions, (C) Double line slit reconstruction at 200 μm depth showing the resolution of the sensor to be better than 60 μm (adapted from Ref(*19*)), (D) The raw sensor capture for a USAF resolution target showing the missing information due to LED blocks, (E) Reconstructed image capture of the USAF resolution target at 200 μm depth (F) Magnified inset of element 2, group 4 of the resolution target showing better than 60 μm resolution.

**Fig. S19.**
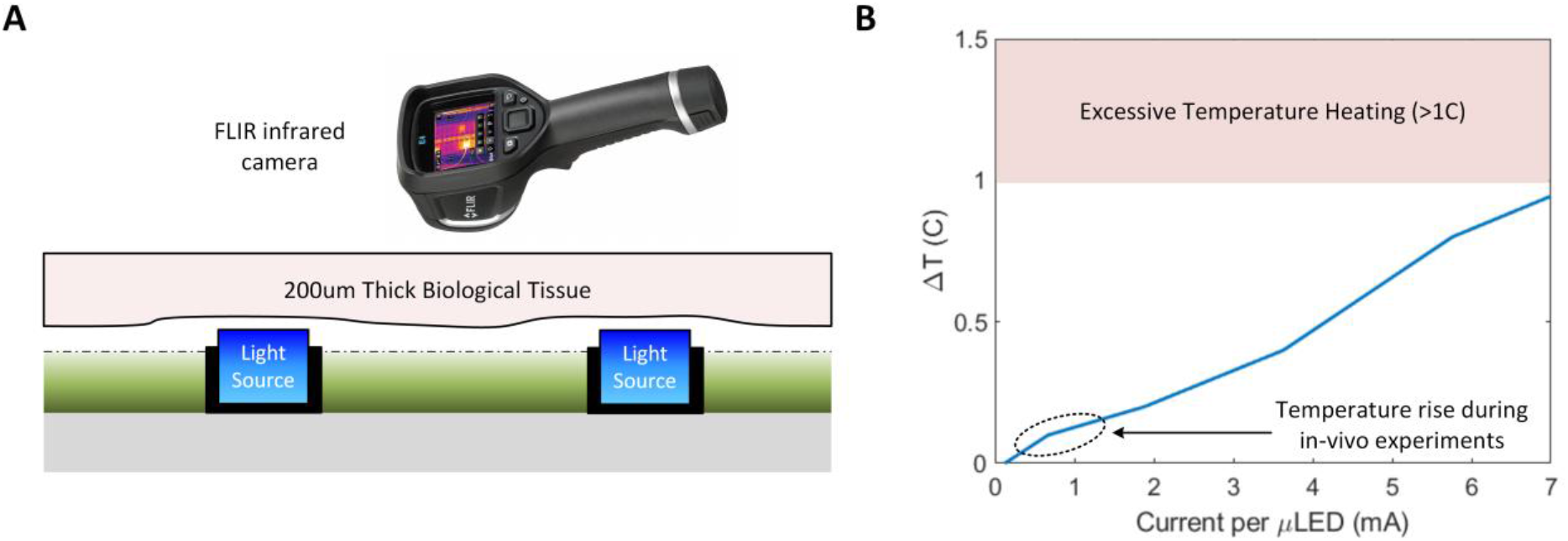
Tissue heating. (A) Cross-section of the CMOS sensor with integrated light sources showing the heating of a 200 μm thick tissue sample due to LED light generation and power dissipation and measured using an FLIR infrared camera. (B) Plot with the rise in temperature versus the current consumed in each μLED; the temperature rise during in-vivo experiments in circled showing that the heating is negligible compared with the safety limits.

**Fig. S20.**
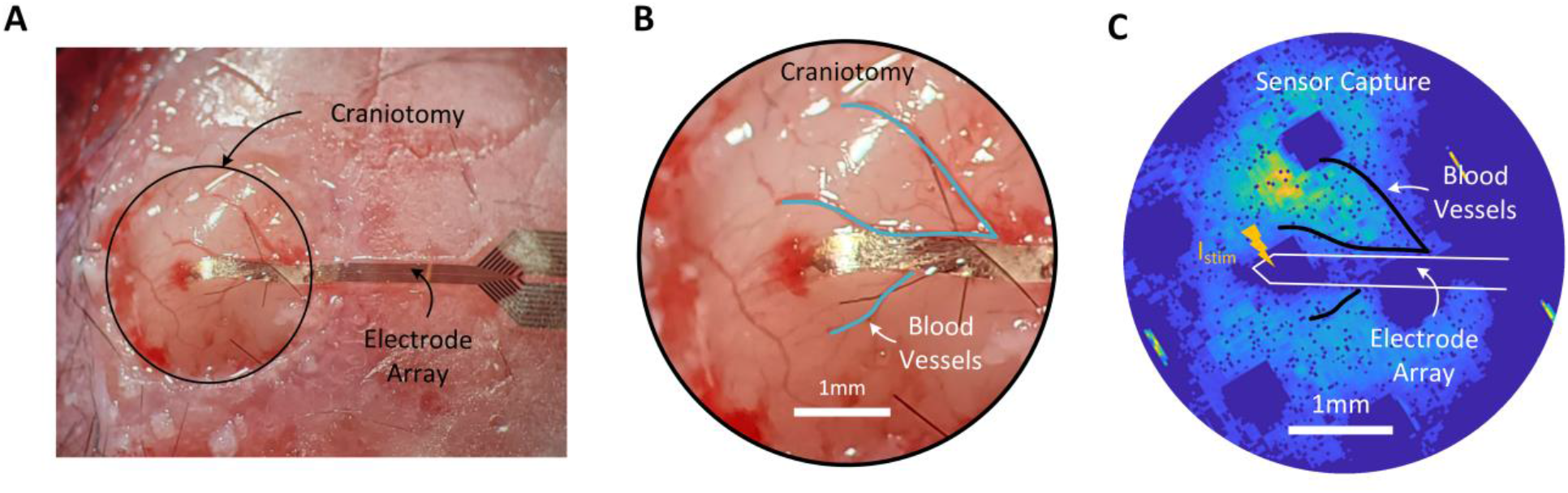
Surgical preparation for electrical stimulation in mouse experiment. (A) The craniotomy preparation including a 6×6 mm^2^ cranial window over the SSC and VC of the left hemisphere of the anesthetized mouse. The flexible shanks are laminated across the cortex and implanted deep into the SC down to Layer 5 for application of electrical current stimulation with varying amplitudes. (B) White light microscope image of the cranial window showing the insertion location of the flexible shanks relative to major blood vessels. (C) The SCOPe image capture is aligned according to these experimental and physiological features to locate the stimulus location. Images are captured at 40 Hz and the full image is cropped to only pixel locations that are in contact with the neural tissue in the cranial window.

**Fig. S21.**
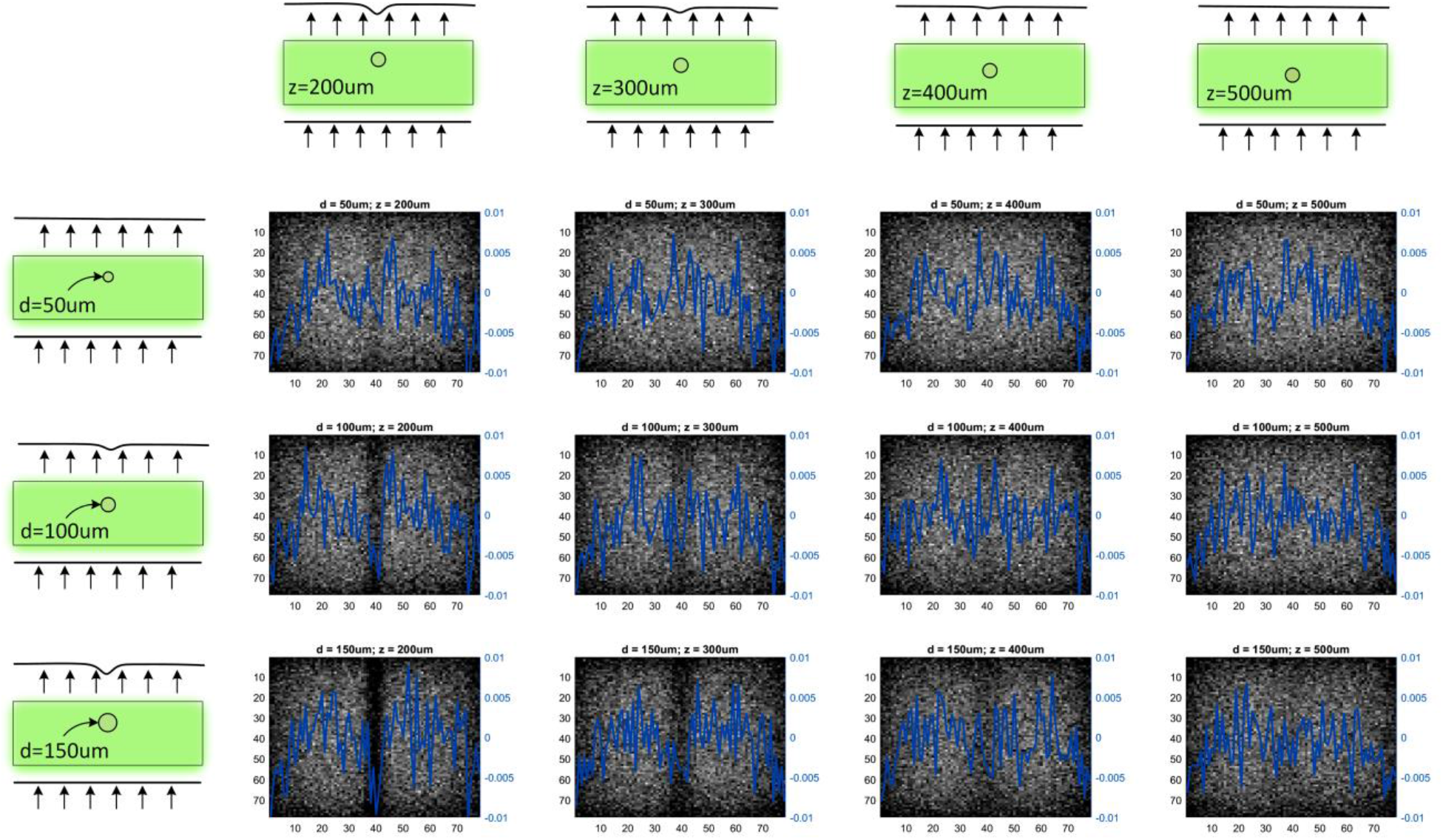
Blood vessel imaging. Monte Carlo setup of a dense fluorescence scene and the occlusion of a single blood vessel of diameters of 50, 100, and 150 μA (along the row axis) and z-depths of 200, 300, 400, and 500 μm (along the column axis). A 3×4 matrix of the transmittance at the tissue-air interface of the fluorescent scene. The blue line overlay shows the cross-sectional counts at the middle row. To image a contrast of ~1% with an SNR > 3 necessitates the detection of 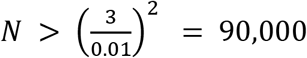 photons.

**Fig. S22.**
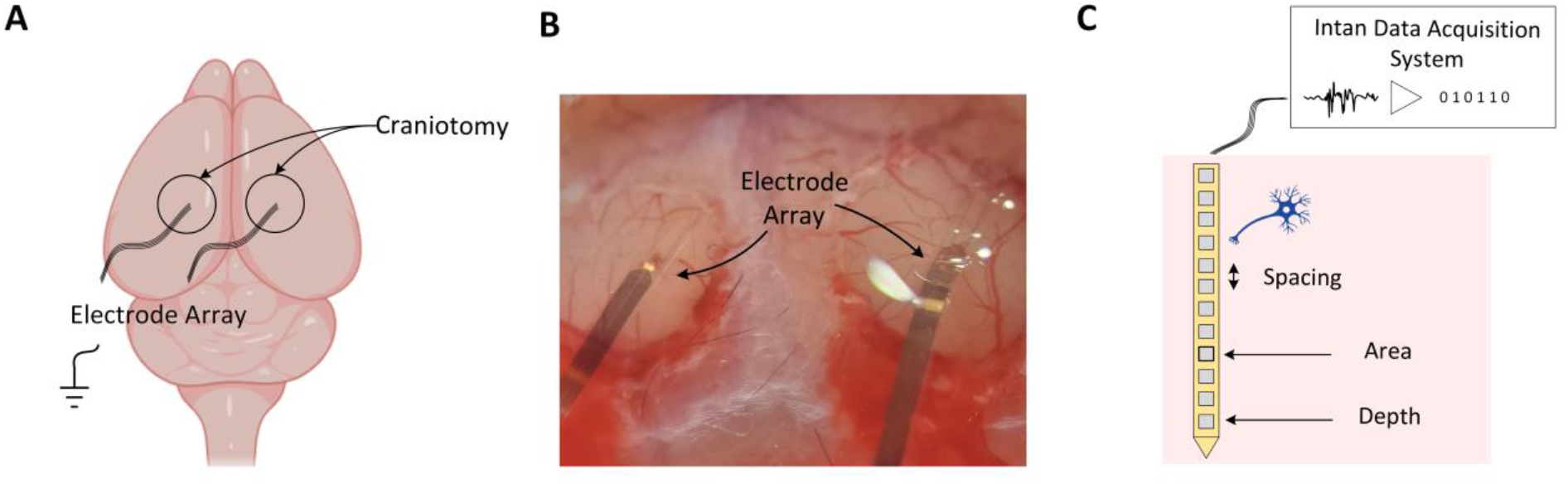
Surgical preparation for optogenetic stimulation in mouse experiment. (A) The craniotomy preparation including a 6×6 mm^2^ cranial window over the SSC and VC of the left hemisphere of the anesthetized mouse. The flexible shanks are laminated across the cortex and implanted deep into the SSC down to Layer 5 for application of electrical current stimulation with varying amplitudes. (B) White light microscope image of the cranial window showing the insertion location of the flexible shanks relative to major blood vessels. (C) The sensor image capture is aligned according to these experimental and physiological features to locate the stimulus location. Images are captured at 40 Hz and the full image is cropped to only pixel locations that are in contact with the neural tissue in the cranial window.

**Fig. S23.**
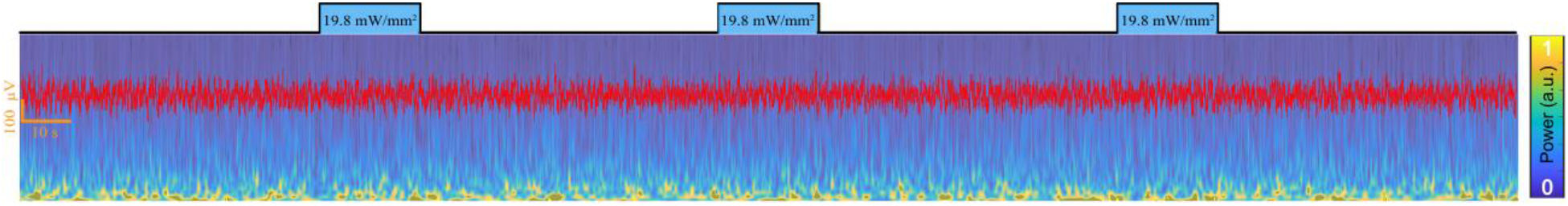
Control experiments for neural activity modulation *in vivo* in wild type mice. LED activation over the implanted electrode array does not show any LFP modulation or artifact generation before and after stimulation.

**Fig. S24.**
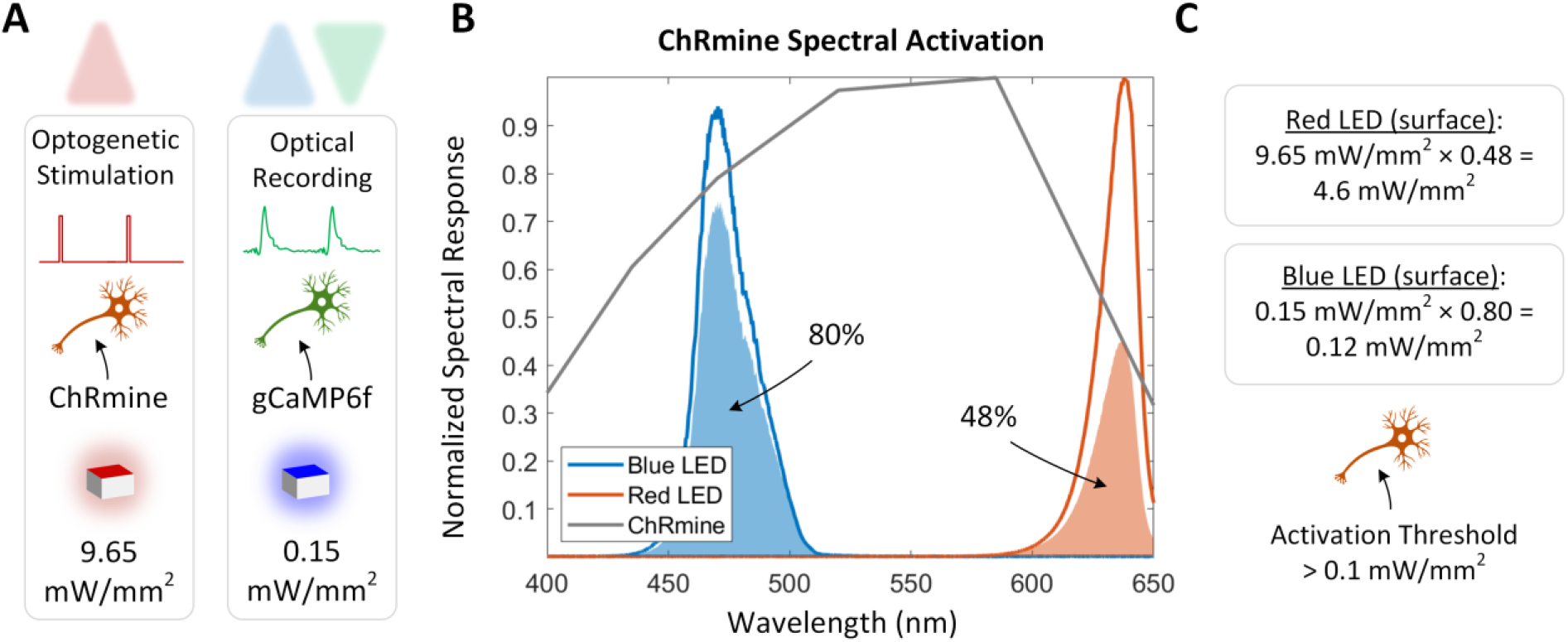
Spectral overlap of LEDs and ChRmine opsin. (A) In the optogenetic stimulation, optical recording in vivo mouse experiment, the operating point of the red LED for stimulation was set at 9.65mW/mm^2^ and blue LED for recording was set at 0.15mW/mm^2^ (**Table S4**). (B) The spectral overlap of the blue excitation LED with the excitation filter and the ChRmine activation spectrum is 80%. The spectral overlap of the blue excitation LED with the excitation filter and the ChRmine activation spectrum is 48%. (C) This corresponds to an emitted red stimulation light power density of 4.6mW/mm^2^ and blue recording light power density of 0.12mW/mm^2^. The activation threshold for ChRmine is 0.1 mW/mm^2^(*46*).

**Fig. S25.**
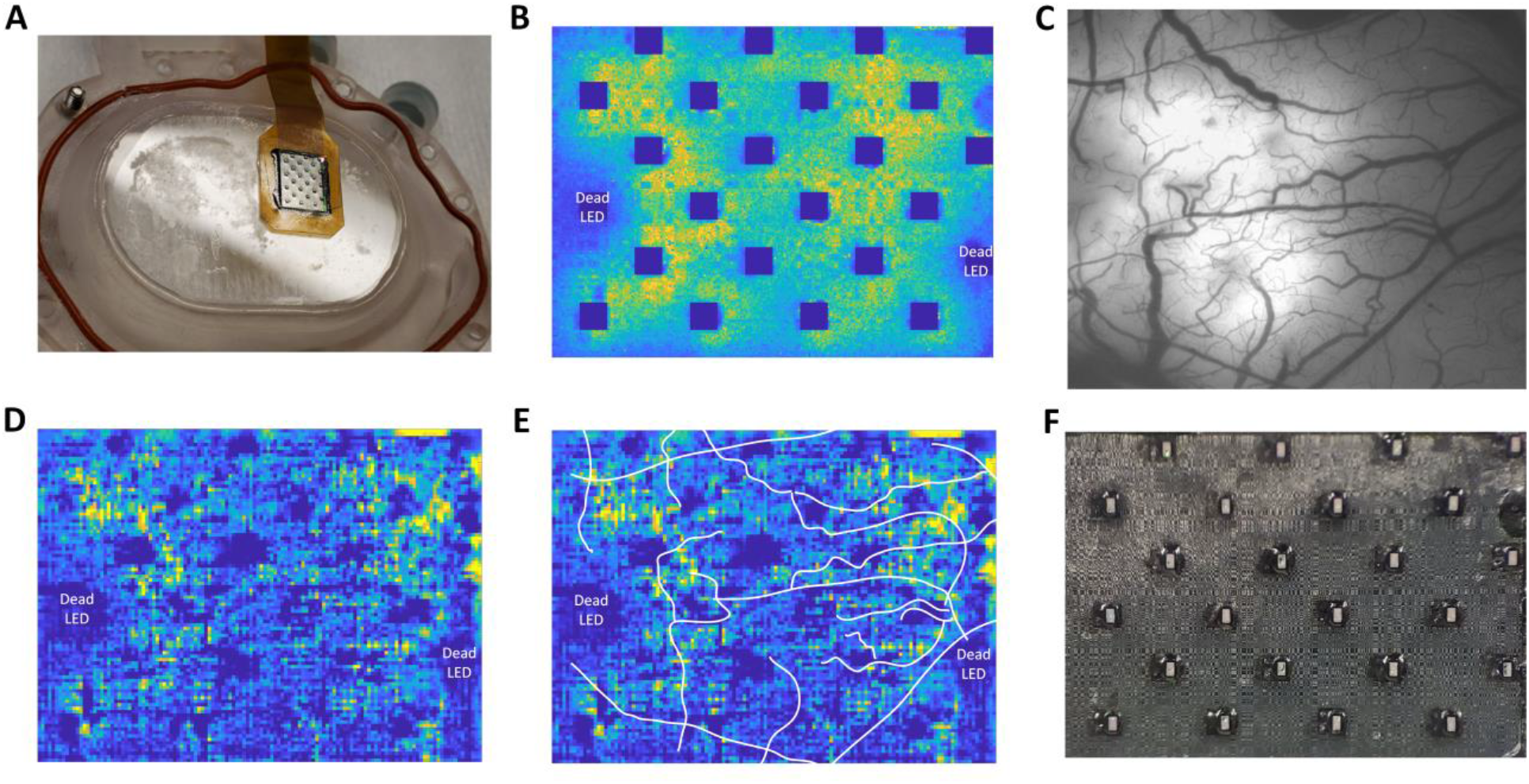
NHP reconstruction. The device is packaged with all the necessary (24) blue μLEDs, excitation and emission filters, and biocompatible parylene coating. The targeted location on the brain port is for the PMd and M1 brain regions. (A) Camera photo taken of the device prior to implantation. The SCOPe is epoxy bonded on top of the cranial window and sterilized. (B) The non-binned spatial capture (192 × 256 pixels) averaged over the entire 50 seconds of a reaching recording. There are 22/24 functional μLEDs on the chip allowing to take advantage of the entire sensor for imaging. (C) Ground truth image of gCaMP8m fluorescence expression in the cortical region under the sensor showing the blood vasculature. (D) Sensor reconstruction image refocused to a calibration depth of 200μm. (E) The refocused image with the blood vasculature overlaid in white lines. (F) Microscope capture of the SCOPe sensor post-experiment showing fully intact device capable of repeat experiment after another sterilization process. There is some biological residue in the top left corner of the chip which also will account for reduced performance in the image reconstruction process.

**Fig. S26.**
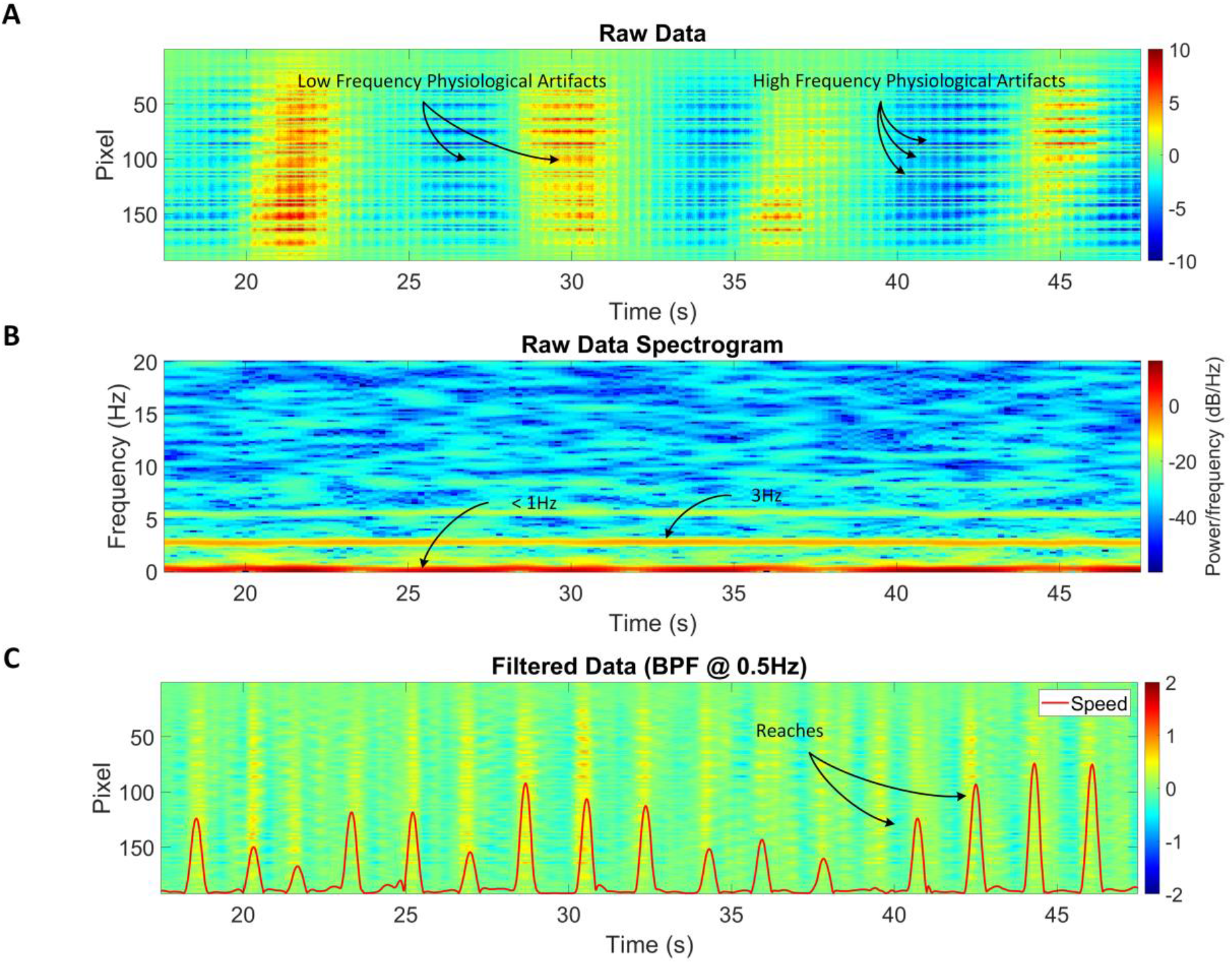
Spectrogram. (A) Example of the space-time matrix of the raw data captured by the sensor containing physiological artifacts. (B) Low frequency 0.1-Hz components are related to blood oxygenation and high frequency 3Hz components are related to heartbeat. (C) Space-time matrix of the temporally bandpass filtered data showing neural activations that are timed synchronously with the reaching tasks.

**Fig. S27.**
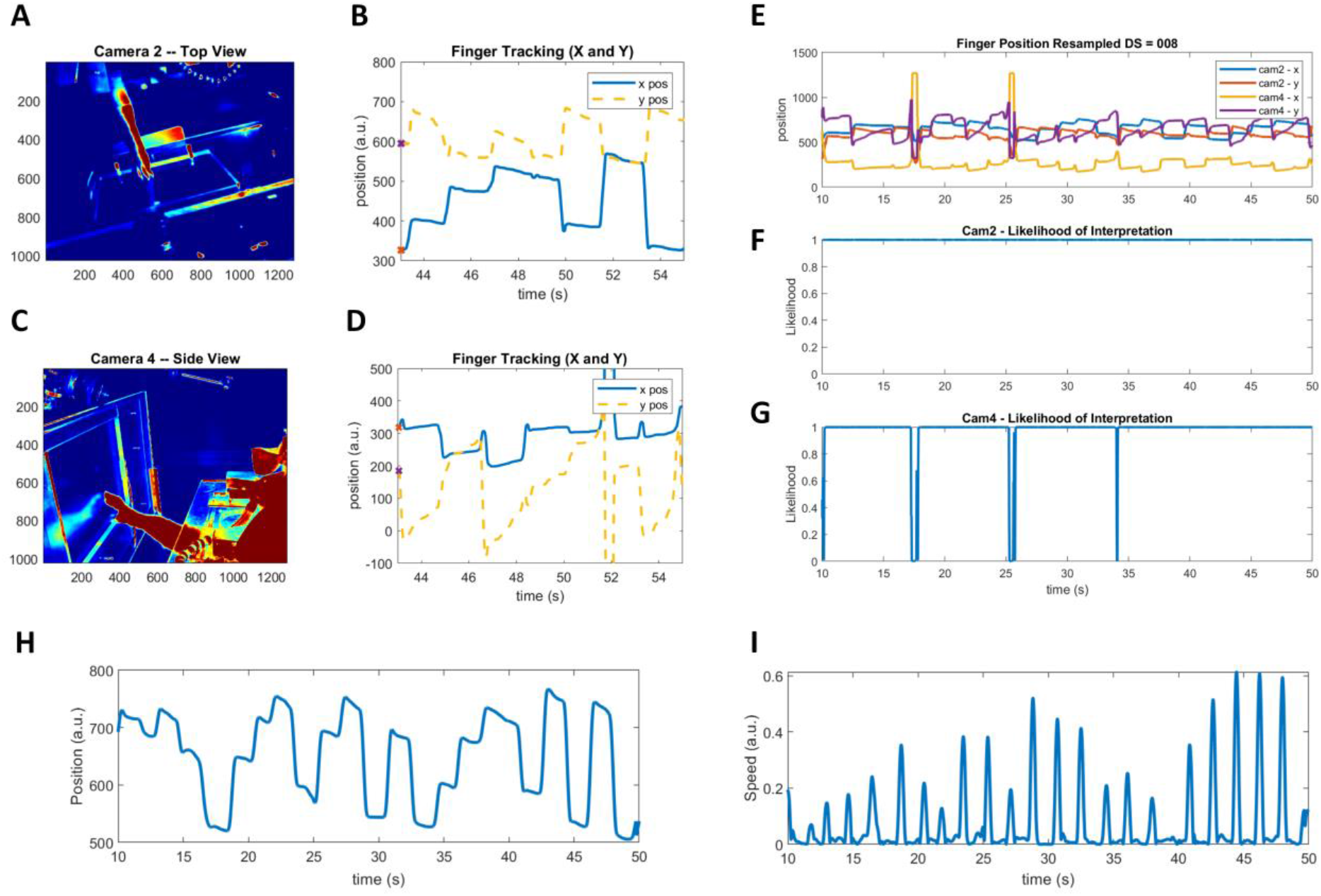
DLC finger tracking. The finger tracking position is provided by synchronizing and mapping in the x and y direction the position of the finger in each of the successive frames, and it is used to identify the onset and completion of reaching tasks and map to the sensor data capture. (A) Camera two top view infrared camera for finger tracking in left-right and forward-backward dimensions. (B) Finger tracking positions as output from the DLC pipeline with the left-right positions (blue line) and the forward-backward positions (yellow dotted line). (C) Camera four side view infrared camera for finger tracking in the up-down dimension and forward-backward dimensions. (D) Finger tracking positions with the up-down positions (blue line) and the forward-backward dimension (yellow dotted line). (E) The finger position given for 40 seconds of experimental data for camera 2, x and y positions, and camera 4, x and y positions. (F) The DLC likelihood of interpretation for camera 2; a likelihood of 1 represents a correct positional estimation, and a likelihood of 0 represents an incorrect positional estimation. Incorrect positional estimations occur due to moments when the finger position falls outside the field of view of the camera. (G) The DLC likelihood of interpretation for camera 4. (H) The estimated finger position given for 40 seconds of experimental data for camera 2, x position and temporally smoothed in time by 10 samples. (I) The corresponding speed calculated from the absolute value of the first derivative of position and temporally smoothed in time by 10 samples.

**Fig. S28.**
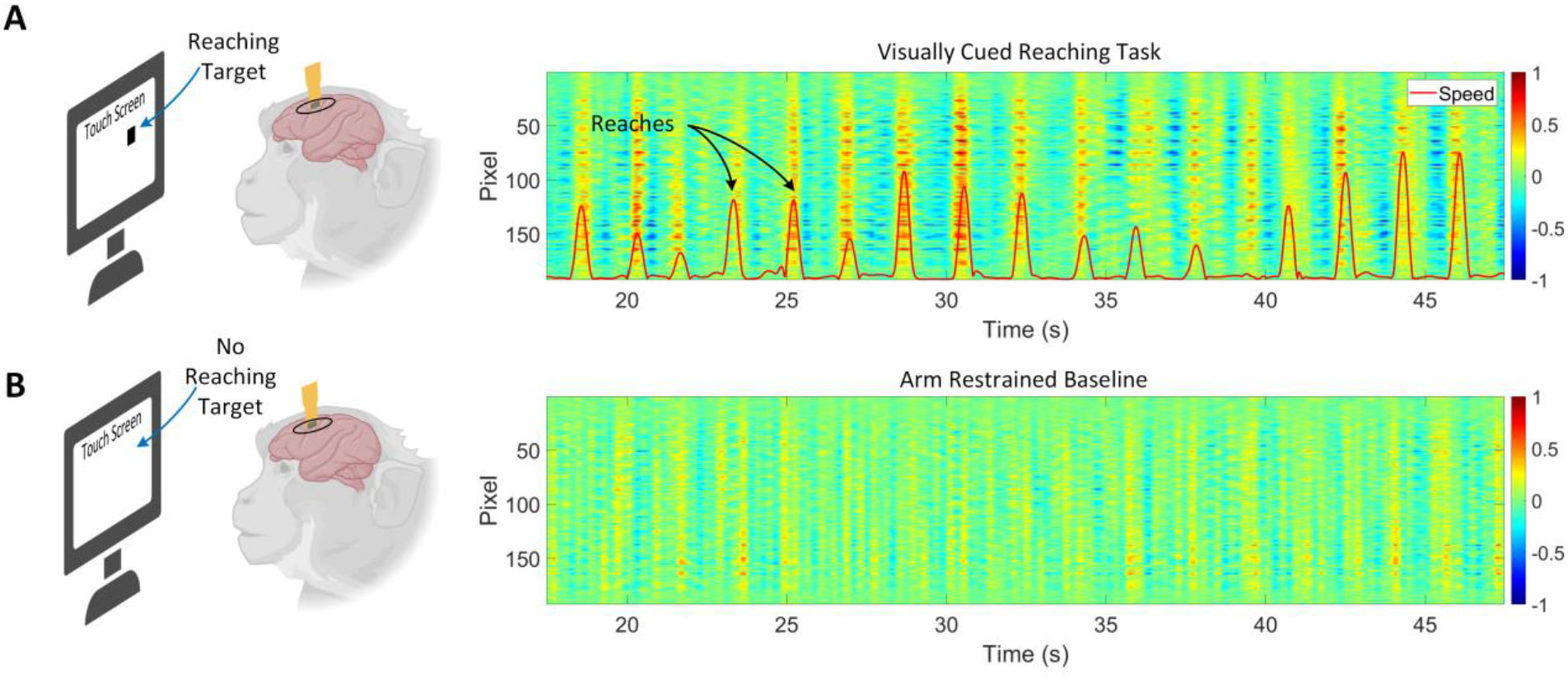
Reaching versus arm-restrained baseline activity. (A) With no arm restraints, activity is synchronized to reaching events and shows good correlation between neural activity and finger speed. (B) The same activations in neural response do not exist in the arm-restrained baseline experiment.

**Fig. S29.**
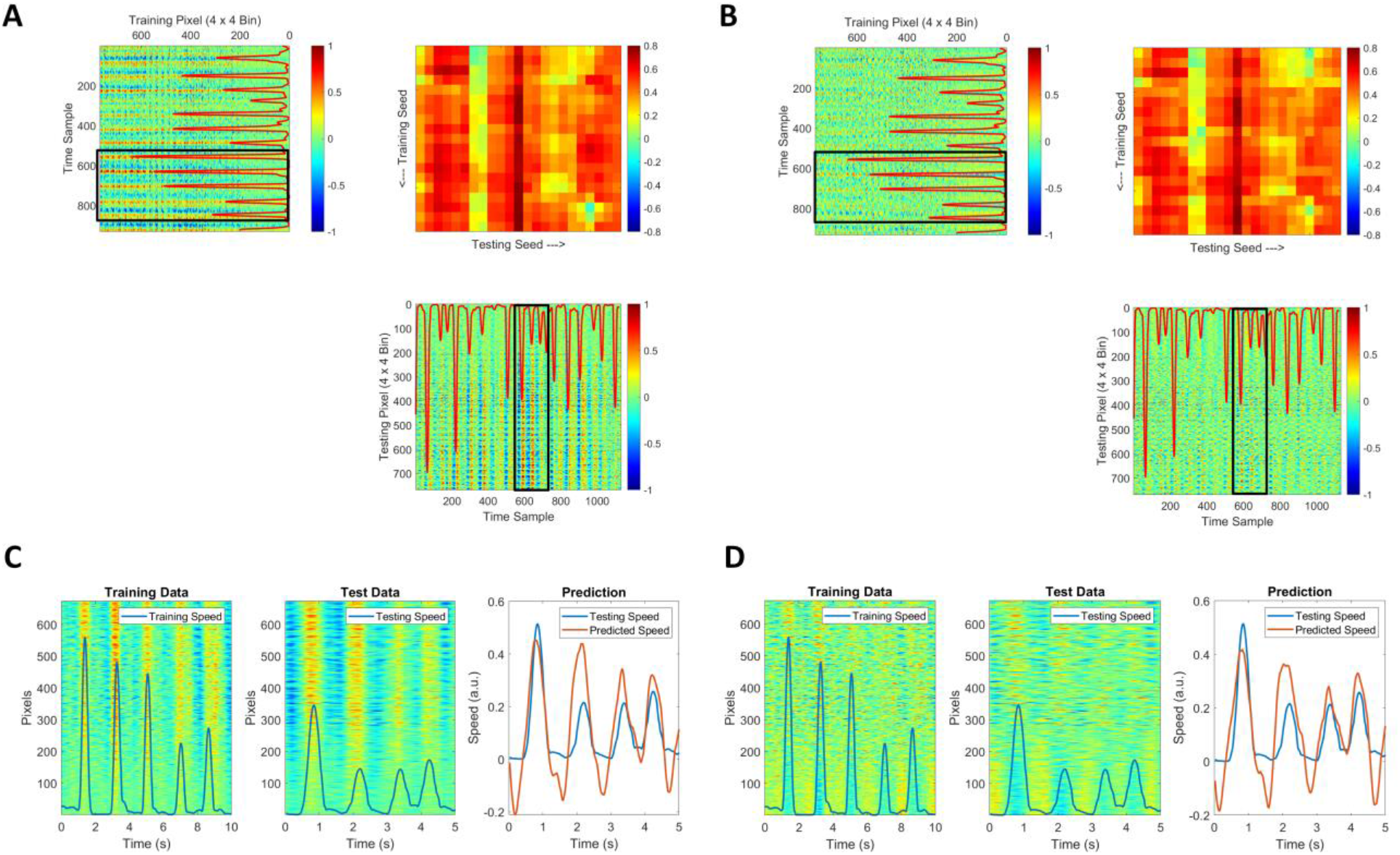
Speed feature decoder correlation matrix. (A) Speed Feature Decoder matrix with training seed on the rows and testing seed on the columns. Speed (blue line, arbitrary units) at each time sample overlaid with the training and testing space-time datasets. The decoder correlation matrix with respect to speed gives good correlation (*ρ* > 0.4) throughout the training and testing datasets. (B) The same Speed Feature Decoder matrix with the sensor-wide mean subtracted from the sensor data. This shows that the performance of the decoder is not affected by subtraction of the sensor-wide mean and that there is spatial structure encoded in the sensor data corresponding to speed prediction. (C) Example training data and testing data snippets represented by the corresponding red rectangles in the training and testing matrices from (A). The example testing and prediction speeds show the good correlation which maps to a single pixel in the correlation matrix. (D) Same as (C) except the sensor-wide mean is subtracted from each of the time samples.

**Fig. S30.**
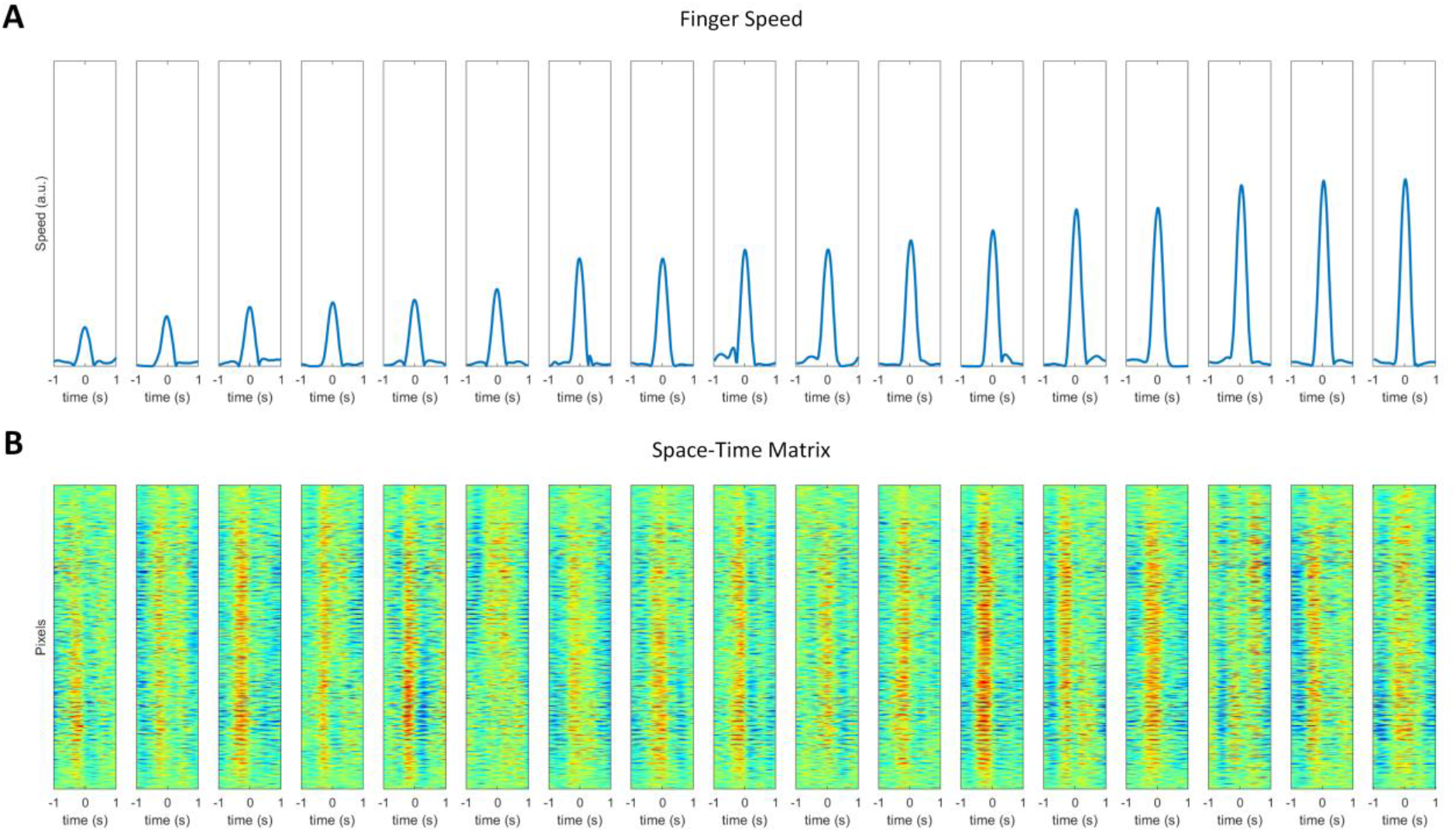
Individual reaching events from a single recording. (A) The 17 individual reaching events are extracted and synchronized by indexing the time sample of the peak speed of the DLC tracked finger. (B) A window of +-40 samples from the seed time sample are used to show the neural activity within +- 1 second of onset and termination of reaching movement.

**Fig. S31.**
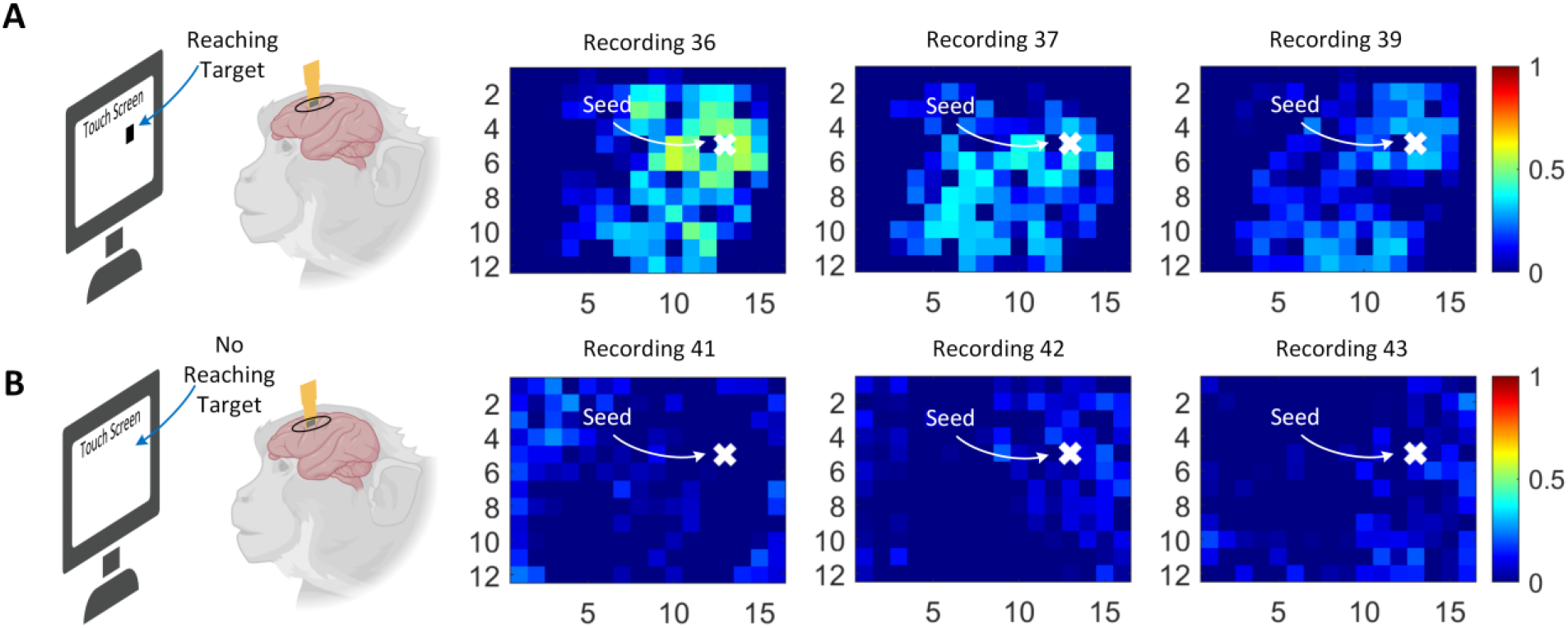
Spatiotemporal correlations. The 2D spatial image showing the correlations at a time lag of 250ms (10 samples) between the seed pixel and the rest of the pixels in the array. The seed pixel is denoted with a white ‘X’ located in the top right of the array corresponding to the PMd brain region on the image sensor. The white circle corresponds to the M1 region and emphasizes the correlation at a neural lag of 250ms. (A) Recordings 36, 37, and 39 correspond with reaching experiments. The white circle is highlighting the positive correlations in neural activations as they travel from PMd to M1. (B) Recordings 41, 42, and 43 correspond with baseline experiments with no visually cued reaches. The white circle is highlighting the no correlations in neural activations between PMd to M1.

**Table S4.** LED optical powers used *in vivo*.

|  | Mouse Exp 1 | Mouse Exp 2 | NHP Exp 1 |
| --- | --- | --- | --- |
| # Of Blue LEDs | 6 | 4 | 22 |
| Optical Power per Blue LED | 4.5μW | 3μW | 9μW |
| # Of Red LEDs | 0 | 1 | 0 |
| Optical Power per Red LED | - | 193μW | - |

**Table S5.** Recordings in nonhuman primate experiment.

| Recording | LED Voltage | Comments |
| --- | --- | --- |
| 0-5 | 0 | Dark |
| 6-9 | 2.48 | Simple Touch |
| 10-11 | 2.49 | Simple Touch |
| 12 | 2.5 | Simple Touch |
| 13 | 2.48 | Baseline |
| 14-22 | 2.48 | Contralateral |
| 23-26 | 2.48 | Ipsilateral |
| 27-28 | 2.48 | Baseline |
| 29-33 | 2.48 | Ipsilateral |
| 34-35 | 2.48 | Contralateral |
| 36-40 | 2.48 | Simple Touch |
| 41 | 2.48 | Baseline |
| 42-43 | 2.48 | Arm-restrained baseline |

**Movie S1.**
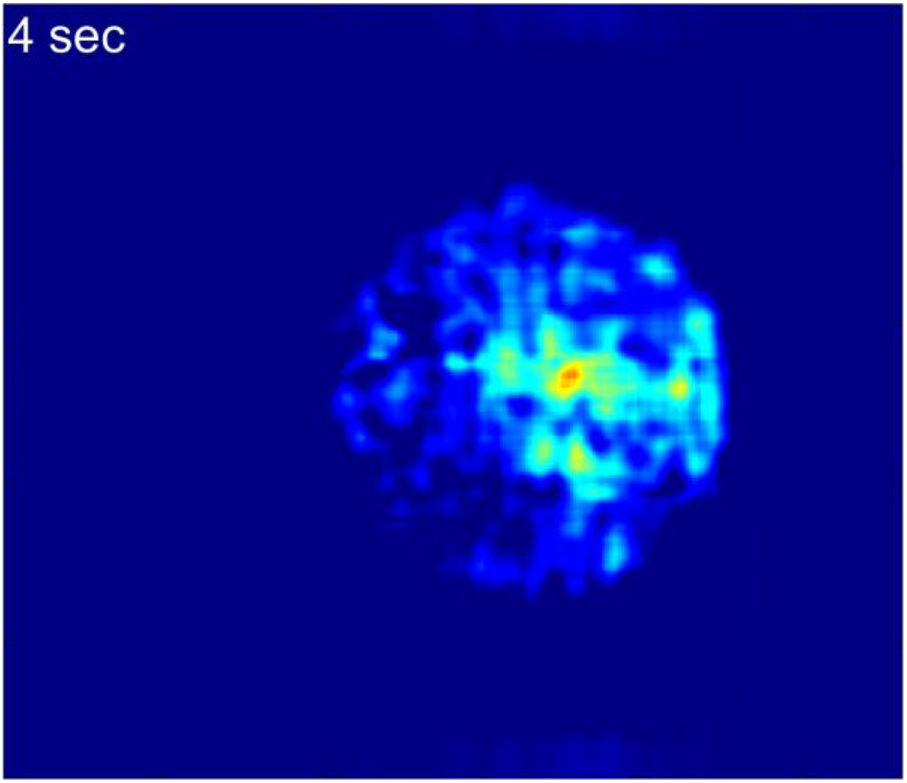
In vivo mouse experiment – electrical stimulation and optical recording.

**Movie S2.**
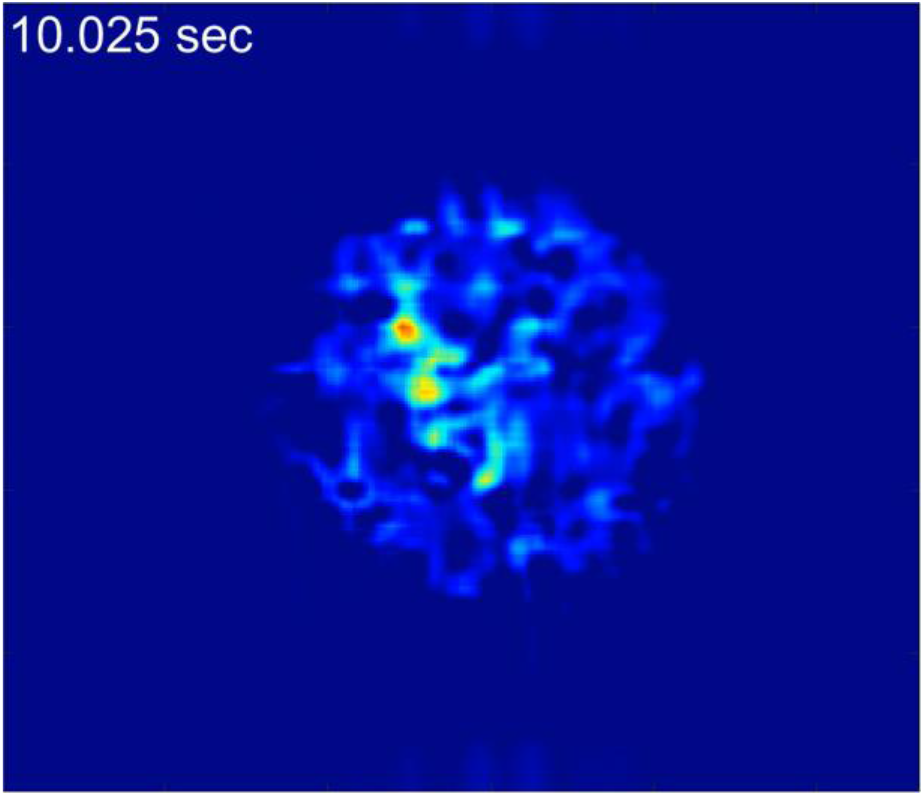
In vivo mouse experiment – optical stimulus and optical recording.

**Movie S3.**
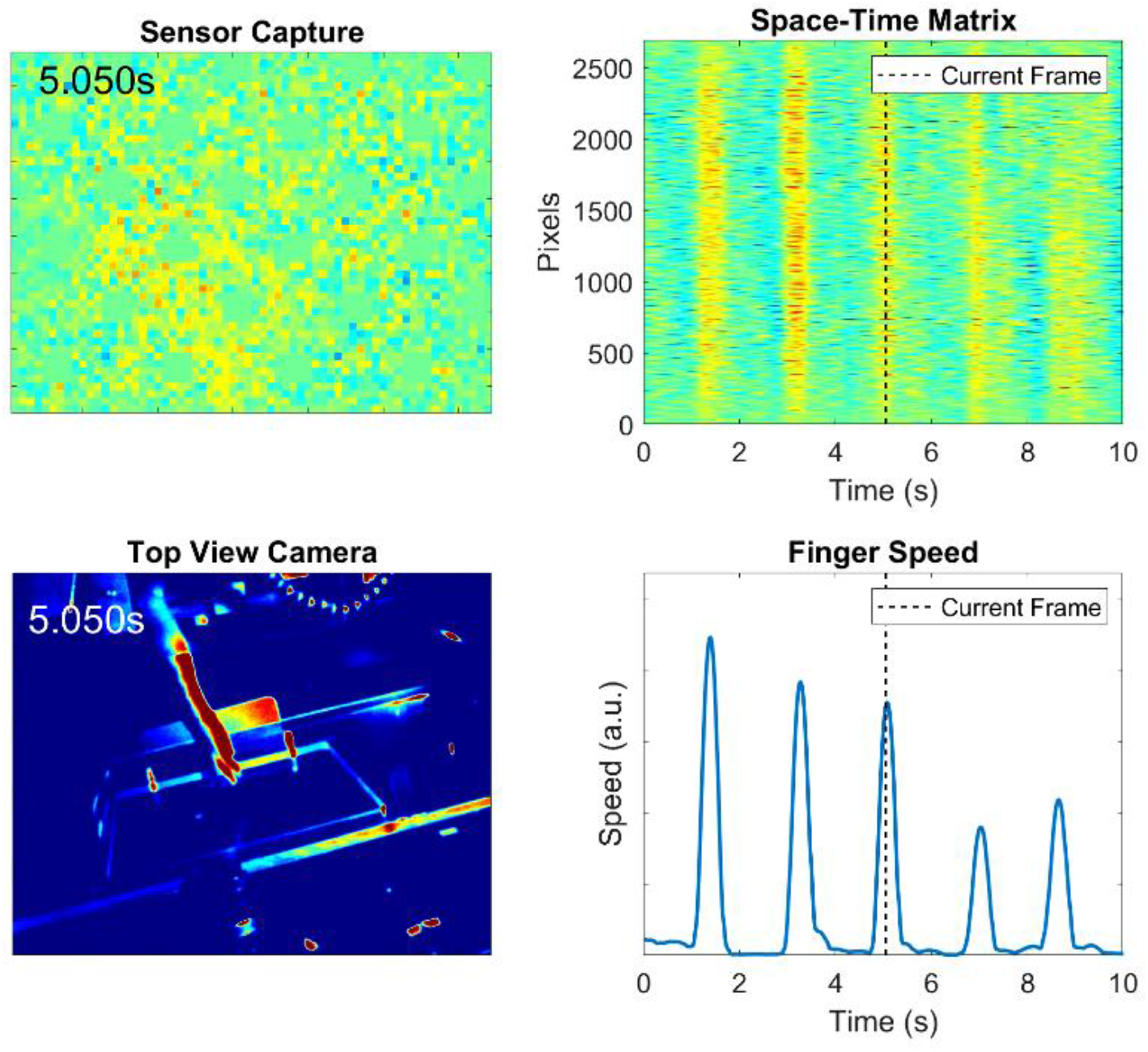
In vivo NHP experiment – behavioral stimulus and optical recording.

